# Evo-devo dynamics of hominin brain size

**DOI:** 10.1101/2023.03.20.533421

**Authors:** Mauricio González-Forero

## Abstract

Brain size tripled in the human lineage over four million years, but why this occurred remains uncertain. To advance our understanding of what caused hominin-brain expansion, I mechanistically replicate it in-silico by modelling the evolutionary and developmental (evo-devo) dynamics of hominin-brain size. I show that, starting from australopithecine brain and body sizes, the model recovers the evolution of brain and body sizes of seven hominin species, the evolution of the hominin brain-body allometry, and major patterns of human development and evolution. Analysis shows that in this model the brain expands because it is “socio-genetically” correlated with developmentally late preovulatory ovarian follicles, not because brain size is directly selected for. The socio-genetic correlation causing the recovered hominin brain expansion is generated over development by ecology and possibly culture. Thus, in this model, direct selection that does not favour brain expansion provides a force that developmental constraints divert causing hominin-brain expansion.

## Introduction

The human brain provides the hardware for stunning achievements, but why it evolved remains unresolved. The fossil record shows a sharp expansion in hominin brain size, tripling over the last four million years from australopithecines to modern humans ^1^ while some *Homo* were small-brained ^2;3^. Many hypotheses exist for why such hominin brain expansion occurred ^4–22^ and they are actively tested, often with correlative ^23–25^ or comparative studies in non-hominin species ^26–29^. Yet, establishing what were the causes of hominin brain expansion remains a major multidisciplinary challenge.

An underexploited approach to identifying the causes of hominin brain expansion is by means of mechanistic modelling. Causes can be inferred from the effects of intervention ^30^, but interventions are mostly impossible or impractical for hominin brain expansion. Despite this challenge, interventions are possible in silico in models that mechanistically replicate hominin brain expansion, enabling inference of the causes of the event ^31^. Models of brain evolution often give qualitative predictions for conditions that favour the evolution of large brains ^32–35^. Yet, it is of particular interest that models make quantitative predictions for conditions under which a human-sized brain evolves (e.g., of 1.3 kg) since what favours a large brain may not necessarily yield a human-sized brain, but possibly one too small or too large for a human.

A recent mathematical model — hereafter, the brain model — makes quantitative predictions for conditions under which a given brain size evolves ^36^. The brain model mechanistically replicates the evolution of adult brain and body sizes of six *Homo* species and much of the timing of human development including the length of childhood, adolescence, and adulthood ^37^. Analysis of the brain model ^37^ has found causal, computational evidence that a challenging ecology ^7;15;22^ and possibly culture ^14;19;21^ rather than social interactions ^6;9;12;16^ could have caused hominin brain expansion. In the model, a challenging ecology, where individuals need brain-supported skills to obtain energy, promotes brain expansion ^36^. If additionally, learning has weakly, not strongly, diminishing returns, then human-sized brains and bodies can evolve ^37^. Although the model does not explicitly model cultural dynamics, weakly diminishing returns of learning could in principle arise from culture if skilled individuals can keep learning from accumulated knowledge in the population ^37^. Thus, in the model, hominin brain expansion needs both a challenging ecology and possibly culture, presumably to reap the benefits in adulthood of investing in growing large brains during childhood. In contrast, conflicting interests between social partners enable evolutionary arms races in brain size as proposed by influential hypotheses ^6;9;16^, but the arms races fail to yield evolutionarily stable human-sized brains and bodies given their metabolic costs ^37^. In turn, cooperation ^12^ disfavours brain size evolution as individuals can rely on social partners’ brains to overcome ecological challenges and so can avoid investing in growing an expensive brain ^37^. The model has incorporated basic aspects of leading hypotheses without explicitly modelling every aspect such as information manipulation or relationship management ^12^. Doing so has not been necessary to obtain the evolution of human-sized brains and bodies given the data used for parameter values.

The brain model makes quantitative predictions by explicitly considering development, that is, the construction of the phenotype over life. In particular, the model describes the construction of brain and body sizes over life using energy conservation analysis following the approach of West et al. ^38^. West et al. use energy conservation analysis to obtain an equation describing the developmental dynamics of body size depending on param-eters measuring metabolic costs that can be easily estimated from data ^38^. The brain model implements West et al.’s approach to obtain equations describing the developmental dynamics of brain, reproductive, and somatic tissue sizes depending additionally on genotypic traits controlling energy allocation to the production of each tissue at each age ^36^. For simplicity, reproductive tissue is defined in the model as preovulatory ovarian follicles which determine fertility given that the model considers only females. The developmentally dynamic equations define the developmental constraints, as the phenotype is constrained to satisfy such equations. The brain model thus depends on parameters measuring brain metabolic costs, which are thought to be a key reason not to evolve large brains ^11;17^ and which are easily estimated from existing data ^39^. In the model, the genotypic traits evolve, which leads to the evolution of brain and body sizes in kg, whose units arise from the empirically estimated metabolic costs. The model has identified key parameters that have strong effects on brain size evolution and particular parameter values that enable the evolution of human-scale brains and bodies ^36;37^ (Table 1).

**Table 1:** Key parameters. Human-scale brains and bodies evolve under these parameter values in the brain model. Changing one of these parameters at a time may substantially change the evolved brain or body sizes or their ontogenetic growth, even causing the evolutionary collapse of brain size (Figs. 6 and R of ref. ^36^ and Extended Data Fig. 3 of ref. ^37^). *Estimated from empirical data ^97;39^. †Empirically informed ^98;79^. Mt.: maintenance. Al.: allocation. DRL.: diminishing returns of learning.

| Parameter | Value | Interpretation |
| --- | --- | --- |
| *Brain mt. cost, $B_b$ | 313 MJ/(kg y) | mid |
| †Follicle mt. cost, $B_r$ | 2697 MJ/(kg y) | high |
| *Soma mt. cost, $B_s$ | 30 MJ/(kg y) | low |
| †Memory cost, $B_k$ | 50 MJ/(TB y) | mildly high |
| Learning cost, $E_k$ | 250 MJ/TB | mildly low |
| Brain al. to skill, $s_k$ | 0.5 | mid |
| Newborn skill, $x_{k1}$ | 0 TB | low |
| Env. difficulty, $\alpha$ | 1.15 | mildly high |
| Skill effectiveness, $\gamma$ | 0.6/TB | mildly low |
| Competence, $c(\text{skill})$ | exponential | weakly DRL |

However, further understanding from the brain model has been hindered by the long-standing lack of mathematical synthesis between development and evolution ^40–42^. To consider developmental dynamics, the brain model was evolutionarily static: it had to assume evolutionary equilibrium where fitness is maximised and so was analysed using dynamic optimisation, specifically using optimal control theory as is standard in life history theory ^43–46^. This was done because of the long-standing lack of mathematical integration of development and evolution, which meant that there were no tractable methods to mathematically model the evolutionary and developmental dynamics of the brain model. Indeed, approaches available at the time that mathematically integrated developmental and evolutionary dynamics required computation of functional derivatives and solution of integro-differential equations ^47;48^, both of which are prohibitively challenging for the relatively complex brain model. Yet, considering the evolutionary dynamics could yield richer insight. For instance, a debated topic is the roles of selection and constraint in brain evolution, often studied with correlational approaches ^49–53^. Considering the evolutionary dynamics in the brain model could enable causal analyses of these roles in hominin brain expansion. Indeed, the short-term evolutionary dynamics can be described as the product of direct selection and genetic covariation assuming negligible genetic evolution ^54;55^, where genetic covariation is a key descriptor of evolutionary constraints ^54;56;57^. Using this separation, Grabowski found that selection for brain size drove brain and body size increases from *A. afarensis* to *H. sapiens* assuming constant genetic covariation for the duration of a species existence ^58^. Yet, the lack of mathematical integration of developmental and evolutionary dynamics has meant that there is a lack of tools to separate selection from constraint in long-term evolution, without assuming negligible genetic evolution.

A solution to these difficulties is offered by a recent mathematical framework — hereafter, evo-devo dynamics framework — that integrates evolutionary and developmental (evo-devo) dynamics allowing for mathematically modelling the evo-devo dynamics for a broad class of models ^59^. This framework provides equations that separate the effects of selection and constraint for longterm evolution under non-negligible genetic evolution and evolving genetic covariation. Moreover, the framework provides equations to analyse what selection acts on in the model, how brain metabolic costs translate into fitness costs, and how brain size development translates into genetic covariation.

To gain a deeper understanding of why hominin brain expansion could have occurred, here I implement the brain model ^37^ in the evo-devo dynamics framework ^59^. This yields a model of the evo-devo dynamics of hominin brain size that mechanistically recovers in silico the hominin brain expansion from australopithecines to modern humans and multiple observations of human evolution and development. This evo-devo dynamics approach enables deeper analysis showing that hominin brain expansion occurs in the model because of direct selection on follicle count rather than on brain size (Extended Data Fig. 1). The brain expands in the model because ecology and possibly culture make brain size and developmentally late follicle count “mechanistically socio-genetically” correlated. This notion is similar to that of genetic covariation in quantitative genetics but differs in two aspects. First, “mechanistic” genetic covariation arises from a mechanistic description of development rather than from a regression-based description as in quantitative genetics, which allows one to model long-term rather than only short-term phenotypic evolution ^59;60^. Second, “socio-genetic” covariation considers not only heredity but also the stabilization (legacy) of the phenotype due to social development, that is, how phenotype construction depends on social partners ^59;60^, which includes a mechanistic description of indirect genetic effects ^61^. Social development in the brain model occurs because of cooperation and competition for energy extraction. This mechanistic treatment shows that brain metabolic costs in the model are not direct fitness costs but affect mechanistic socio-genetic covariation, and that the evolutionary role of ecology and culture in the recovered hominin brain expansion is not to affect direct fitness costs or benefits but to generate the socio-genetic covariation that causes brain expansion.

I provide an overview of the model in Methods. I describe the model in detail and derive the necessary equations for the evo-devo analysis in the Supplementary Information (SI). I provide in the SI the computer code ^62^ written in the freely accessible and computationally fast Julia programming language ^63^.

## Results

### Evolution of brain and body sizes of seven hominins

In the brain model, each individual obtains energy by using her skills to overcome energy-extraction challenges that can be of four types: ecological (e.g., foraging alone), cooperative (e.g., foraging with a peer), between-individual competitive (e.g., scrounging from a peer), and between-group competitive (e.g., scrounging with a peer from two peers). The probability of facing a challenge of type *j* at a given age is 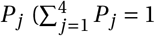, where *j* ∈ {1,…, 4} indexes the respective challenge types). Assuming evolutionary equilibrium, the brain model was previously found to recover the evolution of the adult brain and body sizes of six *Homo* species and less accurately of *Australopithecus afarensis* by varying only the energy extraction time budget (EETB: the proportion of the different types of energy extraction challenges faced) and the shape of energy extraction efficiency (EEE) with respect to one’s own or social partner’s skills ^37^. I recover these results with the evo-devo dynamics approach (Fig. 1). In these results, brain expansion from one evolutionary equilibrium to another is caused by an increasing proportion of ecological challenges and a switch from strongly to weakly diminishing returns of learning. As weakly diminishing returns of learning might arise from accumulated cultural knowledge in the population, this indicates that ecology and possibly culture cause hominin brain expansion in the model ^37^. Below, I analyse further the factors causing such brain expansion.

**Figure 1:**
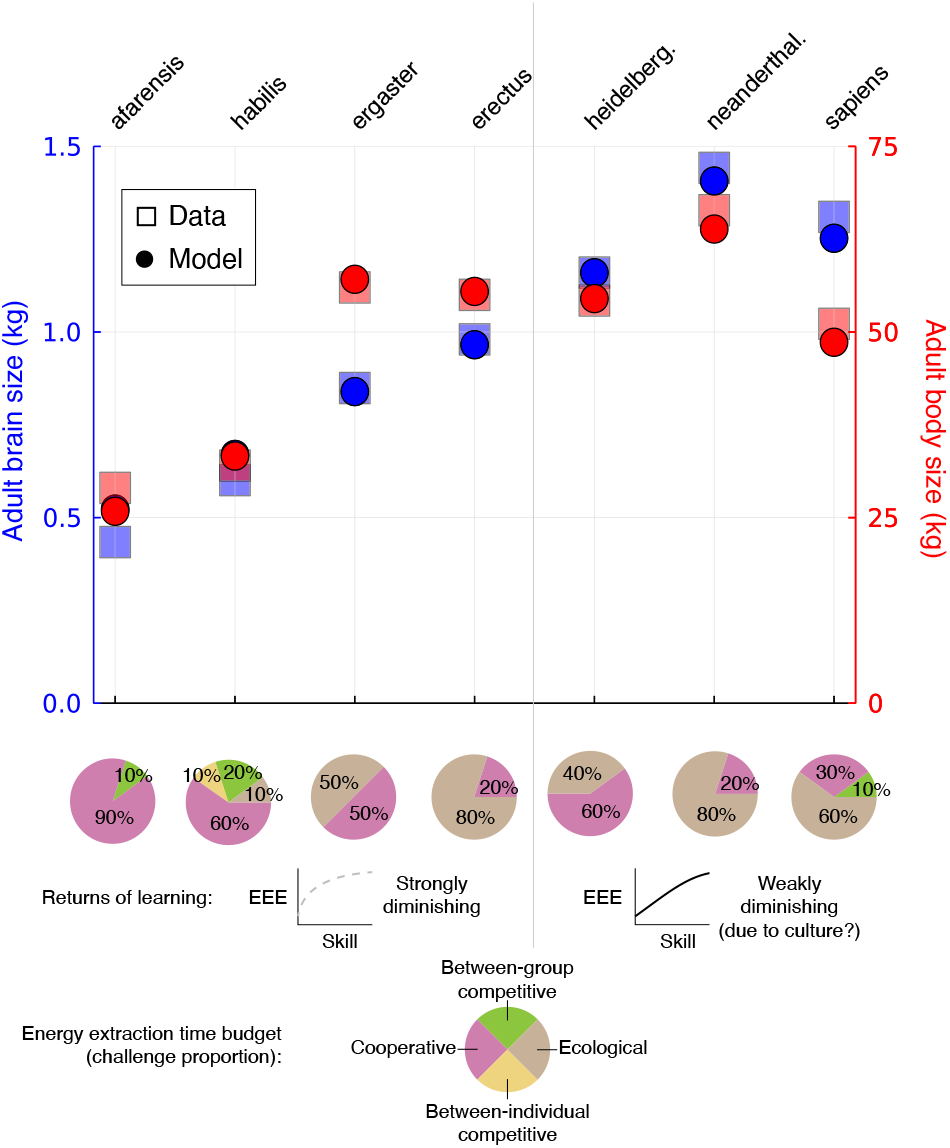
Evolution of brain and body sizes of seven hominin species solely by changing socio-genetic covariation. Adult brain and body sizes of seven hominin species evolve in the model only by changing the EETB, the returns of learning, and how the skills of cooperating partners interact. Squares are the observed adult brain and body sizes for the species at the top (data from refs. ^39;69;99–103^). Dots are the evolved values in the model for a 40-year-old using the evo-devo dynamics approach. Pie charts give the EETB used in each scenario. The returns of learning are either strongly diminishing (power competence) for the left 4 scenarios or weakly diminishing (exponential competence) for the right 3 scenarios. Cooperation is either submultiplicative for the *afarensis* and right 3 scenarios, or additive for the remaining scenarios. These EETBs and shapes of EEE were previously identified as evolving best fitting adult brain and body sizes for the corresponding species assuming evolutionary equilibrium ^37^. In principle, weakly diminishing returns of learning might arise from culture. I will show that varying EETBs and the shape of EEE only varies sociogenetic covariation **L**_**z**_, but not the direction of direct selection *∂w* /*∂***z** or where it is zero (it never is). I refer to the particular EETB and shape of EEE yielding the evolution of adult brain and body sizes of a given species as the species scenario. For the *afarensis* scenario, the ancestral genotypic traits are somewhatNaive2 (Eqs. S46). For the remaining six scenarios, the ancestral genotypic traits are the final genotypic traits of the *afarensis* scenario started from the somewhatNaive2 genotypic traits. The final evolutionary time is 500 for all seven scenarios.

### Emergence of hominin brain-body allometry

To examine the influence of development alone on the developed brain and body sizes, I consider genotypic variation without evolution as follows. Consider the parameter values in the *sapiens* scenario of Fig. 1, which yield the evolution of brain and body sizes of *H. sapiens*. Under those parameter values and without evolution, randomly sampled genotypes develop adult brain and body sizes generating a tight brain-body allometry with slope 0.54 (*R*^2^ = 0.95; Fig. 2a). A similar slope but with a lower placement is found in other primates and mammals ^64^ (Fig. 2a). As there is only development but no evolution in Fig. 2a, this 0.54 slope arises purely from developmental canalization sensu Waddington ^65^. For the sample size used, no organism with random genotype reaches hominin brain and body sizes (no black dot in green region in Fig. 2a). The recovered brain-body allometry from developmental canalization has a high placement (“intercept”), so that the developed brain size is relatively large for the developed body size. In simpler models of development, an allometry with high placement is known to arise with a growth rate, developmentally initial size, or growth duration that is high for the predicted variable (here adult brain size) relative to the predictor (here adult body size) ^66^. In Fig. 2a, brain size can have a high growth rate and growth duration because of the parameter values in the *sapiens* scenario including a high proportion of difficult ecological challenges, weakly diminishing returns of learning, and a high metabolic cost of memory (Figs. 3 and 6 of ref. ^36^ and Extended Data Fig. 1 of ref. ^37^). Hence, in the brain model under the *sapiens* scenario, development alone has a strong influence on the developed brain and body sizes, with a bias ^57^ toward large brains, but is unlikely to yield hominin brain sizes without selection.

**Figure 2:**
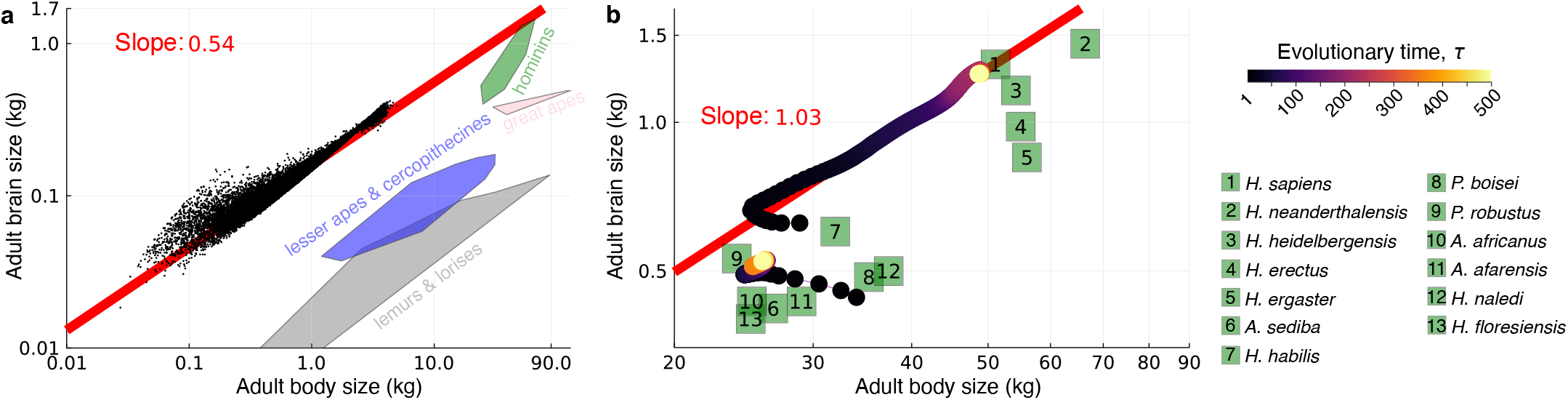
Static, non-evolutionary and evo-devo dynamic brain-body allometries. **a**, Brain-body allometry without evolution. Dots are the brain size at 40 years of age vs body size at 40 years of age in log-log scale, developed under the brain model from 10^6^ randomly sampled genotypes (i.e., growth efforts, drawn from the normal distribution with mean 0 and standard deviation 4) using the parameter values of the *sapiens* scenario. Only “non-failed” organism are shown, that is, those having a body not entirely composed of brain at 40 years of age, which are approximately 4% of 10^6^. The remaining 96% are “failed” organisms (not shown) at 40 years of age, having small bodies (< 100 grams) entirely composed of brain tissue due to tissue decay from birth (Fig. S8). Coloured regions encompass extant and fossil primate species. **b**, Brain-body allometry with evolution. Dots are the brain size at 40 years of age vs body size at 40 years of age over evolutionary time in log-log scale for two trajectories. The bottom trajectory uses the parameter values of the *afarensis* scenario (Fig. 1) and somewhatNaive2 ancestral genotypic traits; in the bottom trajectory, adult and brain body sizes evolve from those approaching *P. boisei* to those approaching *P. robustus*. The top trajectory uses the parameter values of the *sapiens* scenario (Fig. 1) and the evolved genotypic traits of the bottom trajectory as ancestral genotypic traits; in the top trajectory, adult and brain body sizes evolve from those approaching *H. habilis* toward those of *H. sapiens*. A linear regression over this trajectory yields a slope of 1.03 (red line). Adult values for 13 hominin species are shown in green squares. Brain and body size data for non-hominins are from ref. ^64^ excluding three fossil, outlier cercophitecines; brain and body size data for hominins are from refs. ^2;3;39;69;99–103;64;104;105^ using only female data when possible. Fossil data may come from a single individual and body size estimates from fossils are subject to additional error. *H*.: *Homo, A*.: *Australopithecus*, and *P*.: *Paranthropus*.

**Figure 3:**
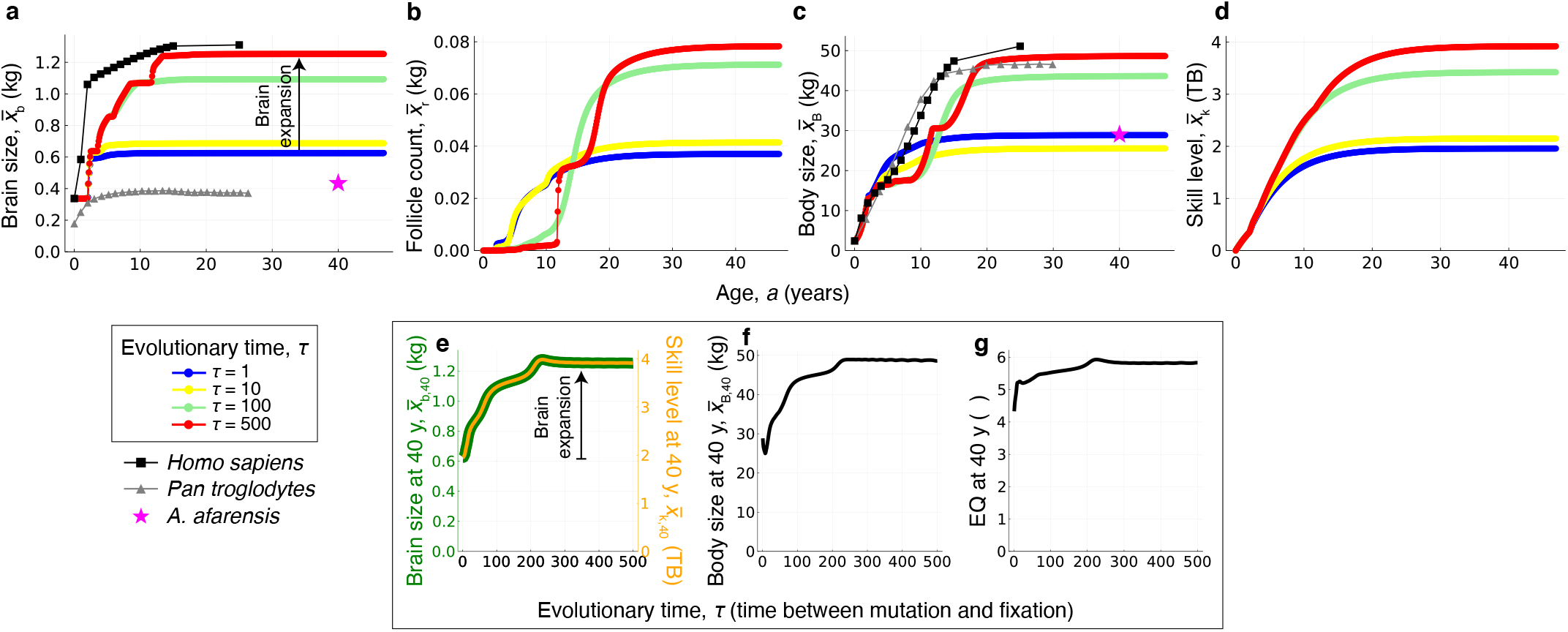
Evo-devo dynamics of hominin brain size. **a-d**, Developmental dynamics over age (horizontal axis) and evolutionary dynamics over evolutionary time (differently coloured dots; bottom left label). Evo-devo dynamics of: **a**, brain size; **b**, follicle count; **c**, somatic tissue size; and **d**, skill level. Evolutionary dynamics of (**e**) brain size (green),(**e**) skill level (orange), (**f**) body size, and (**g**) encephalisation quotient (EQ) at 40 years of age. **a**,**c**, The mean observed values in a modern human female sample are shown in black squares (data from Table S2 of ref. ^39^ who fitted data from ref. ^69^). The mean observed values in *Pan troglodytes* female samples are shown in gray triangles (body size data from Fig. 2 of ref. ^106^; brain size data from Fig. 6 of ref. ^70^). The mean observed values in *A. afarensis* female samples are shown in pink stars (data from Table. 1 of ref. ^103^). One evolutionary time unit is the time from mutation to fixation. If gene fixation takes 500 generations and one generation for females is 23 years ^107^, then 300 evolutionary time steps are 3.4 million years. The age bin size is 0.1 year. Halving age bin size (0.05 year) makes the evolutionary dynamics twice as slow but the system converges to the same evolutionary equilibrium (Fig. S3). I take adult phenotypes to be those at 40 years of age as phenotypes have typically plateaued by that age in the model. All plots are for the *sapiens* trajectory of Fig. 2b.

Letting evolution proceed, I find that the evolved brain and body sizes strongly depend on the ancestral genotypic traits. For instance, under the *sapiens* scenario, the ancestral genotypic traits must develop large bodies, otherwise brain size may collapse over evolution (Figs. S4). This may be interpreted as a requirement to evolve from ancestors that had a genotype capable of leading to body growth while facing cooperative and between-group competition challenges. Moreover, the developmental patterns that evolve strongly depend on the ancestral genotypic traits, even if the evolved adult brain and body sizes are the same. The dependence of the evolved traits on ancestral conditions is sometimes called phylogenetic constraints, which are typically assumed to disappear with enough evolutionary time ^67^. The evodevo dynamics framework finds that phylogenetic constraints do not necessarily disappear with enough time because genetic constraints are necessarily absolute in long-term evolution ^59^. This is because there is sociogenetic covariation only along the path where the developmental constraint is met (so **L**_**z**_ in Eq. M5, a mechanistic, generalised analogue of Lande’s ^54^ **G** matrix, is singular) which means that the evolutionary outcome depends on the evolutionarily initial conditions ^68;59^.

To identify suitable ancestral genotypic traits to model hominin brain expansion, I consider naive ancestral genotypic traits (termed somewhatNaive2) under the *afarensis* scenario. These naive ancestral genotypic traits are such that ancestrally each individual has a high energy allocation to somatic growth at birth and developmentally increasing thereafter, a small allocation to brain growth at birth and developmentally decreasing thereafter, nearly zero allocation to follicle production from birth to 10 years of age, and very small but larger allocation to follicle production from 10 years of age onwards (blue dots in Fig. S1d-f). These ancestral genotypic traits cause individuals to develop brain and body sizes of australopithecine scale, most closely approaching those of *Paranthropus boisei* (initial evolutionary time of bottom trajectory in Fig. 2b and blue dots in Extended Fig. 2a,c). With this ancestral genotype, letting evolution proceed under the *afarensis* scenario yields the evolution of australopithecine brain and body sizes, most closely approaching those of *P. robustus* (bottom trajectory in Fig. 2b). Setting the evolved genotypic traits under this *afarensis* scenario as ancestral genotypic traits and switching parameter values to the *sapiens* scenario yields an immediate plastic change in the developed brain and body sizes approaching those seen in *habilis* (initial evolutionary time of top trajectory in Fig. 2b). Letting evolution proceed yields the evolution of *H. sapiens* brain and body sizes (top trajectory in Fig. 2b). This evolutionary trajectory approaches the observed brain-body allometry in hominins starting from brain and body sizes of australopithecine scale, with a slope of 1.03 (Fig. 2b).

The switch from the *afarensis* to the *sapiens* scenario involves a sharp decrease in cooperative challenges, a sharp increase in ecological challenges, and a shift from strongly to weakly diminishing returns of learning (Fig. 1). While these changes are here implemented suddenly and so lead to an immediate plastic response, the changes may be gradual allowing for genetic evolution.

### Evo-devo dynamics of brain size

Further detail of the recovered hominin brain expansion is available by examining the evo-devo dynamics that underlie the *sapiens* trajectory in Fig. 2b. Such trajectory arises from the evolution of genotypic traits controlling energy allocation to growth. This evolution of energy allocation yields the following evo-devo dynamics in the phenotype.

Adult brain size more than doubles from around 0.6 kg to around 1.3 kg closely approaching that observed in modern human females ^69;70;39^ (Fig. 3a).

In the model, the developmental onset of reproduction occurs when follicle count (in mass units) becomes appreciably non-zero and gives the age of “menarche”. Females ancestrally become fertile early in life with low adult fertility and evolve to become fertile later in life with high adult fertility (Fig. 3b), consistent with empirical analyses ^71–75^.

Body size ancestrally grows quickly over development and reaches a small size of around 30 kg (blue dots in Fig. 3c), and then evolves so it grows more slowly to a bigger size of around 50 kg (red dots in Fig. 3c), consistent with empirical analyses ^71;76^. Body size evolves from a smooth developmental pattern with one growth spurt to a kinked pattern with multiple growth spurts, which are most easily seen as peaks in a weight velocity plot ^77;71;78^ (Fig. S2o inset). The evolved number and pattern of growth spurts strongly depend on the ancestral genotype (Fig. S6o inset). The evolved age at menarche occurs before the last growth spurt (Fig. 3b,c), in contrast to observation ^71^ and previous results ^37^.

Adult skill level evolves expanding from around 2 TB to 4 TB, the units of which arise from the used value of the metabolic cost of memory which is within an empirically informed range ^79^ (Fig. 3d).

The evolved developmental growth rates of phenotypic traits are slower than and somewhat different from those observed and those obtained in the previous optimisation approach ^37^, which was already delayed possibly because the developmental Kleiber’s law I use underestimates resting metabolic rate at small body sizes (Fig. C in ref. ^36^; Fig. S2B in ref. ^80^). The added developmental delays could be partly due to my use of relatively coarse age bins (0.1 year) rather than the (nearly) continuous age used previously ^37^, but halving age bin size (0.05 year) yields the same results (Fig. S3). The added developmental delays might also be partly because of slow evolutionary convergence to equilibrium, and because the evolved ontogenetic pattern depends on the ancestral genotypic traits (compare the red dots of Figs. 3a and S6h).

These patterns generate associated expansions in adult brain, body, and encephalisation quotient (EQ) ^49^ (Fig. 3e-g). EQ measures here brain size relative to the expected brain size for a given mammal body size ^81^. Adult brain size expands more sharply than adult body size (Fig. 3e,f). Consequently, adult brain size evolves from being ancestrally slightly over 4 times larger than expected to being about 6 times larger than expected (Fig. 3g). Thus, the brain expands beyond what would be expected from body expansion alone.

The evo-devo dynamics of brain and body sizes that underlie the *afarensis* trajectory in Fig. 2b are shown in Extended Data Fig. 2. The evolved body size under the *afarensis* scenario shows mild indeterminate body growth (red dots in Extended Data Fig. 2c), reminiscent of that in female bonobos (Fig. 6 of ref. ^82^). Such indeterminate body growth disappears with the plastic change induced by changing to the conditions of the *sapiens* scenario (blue dots in Fig. 3c).

### Analysis of the action of selection

To understand what causes the obtained brain expansion, I now analyse direct selection and genetic covariation which formally separate the action of selection and constraint on evolution. Such formal separation was first formulated for short-term evolution under the assumption of negligible genetic evolution ^54;55^ and is now available for long-term evolution under non-negligible genetic evolution ^59^.

I first analyse the action of selection. In the brain model, fertility is proportional to follicle count whereas survival is constant as a first approximation. Then, in the brain model there is always positive direct selection for ever-increasing follicle count, but there is no direct selection for brain size, body size, skill level, or anything else (Fig. 4a-d; Eq. M3; Extended Data Fig. 1). The fitness landscape has no internal peaks and direct selection only favours an ever higher follicle count (Fig. 5). Since there is only direct selection for follicle count, the evolutionary dynamics of brain size 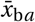 at age *a* satisfy

**Figure 4:**
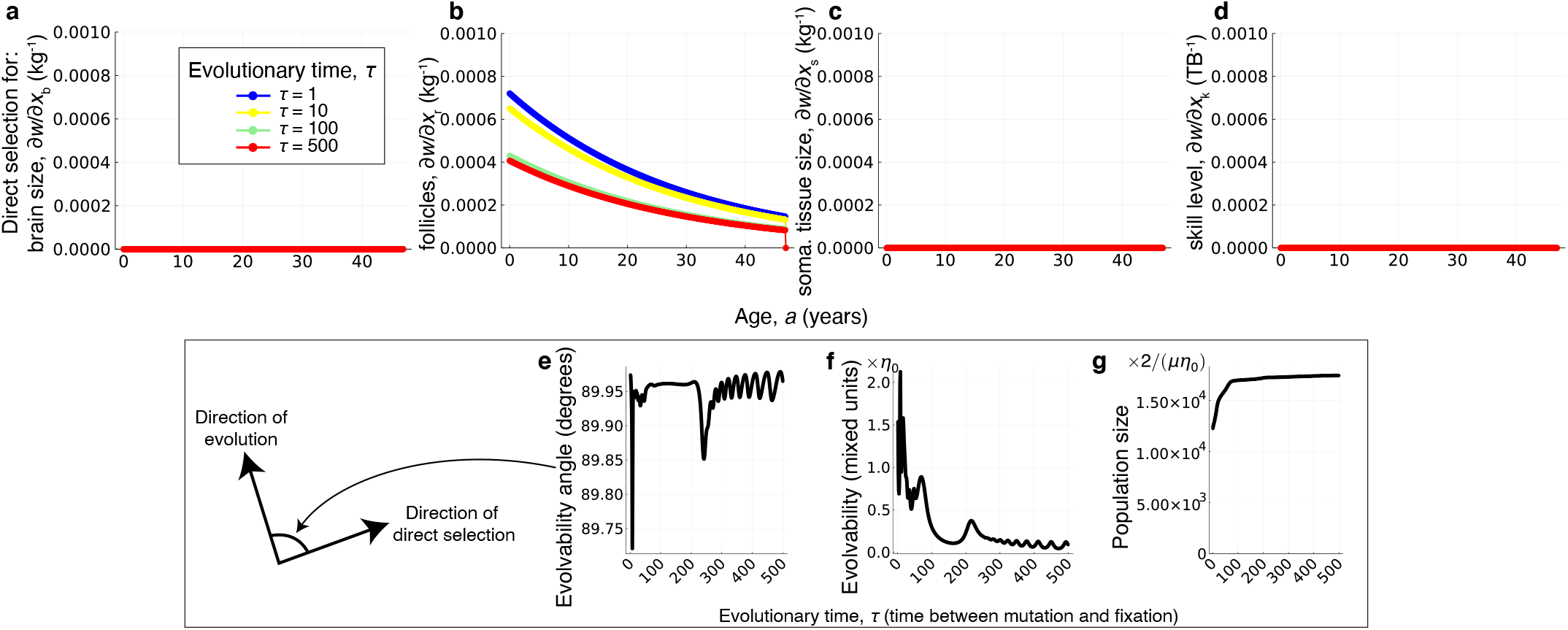
The action of selection. **a-d**, There is only direct selection for follicle count, and such selection decreases with age. **e**, The angle between the direction of evolution and direct selection, both of the geno-phenotype (i.e., genotype and phenotype), is nearly 90 degrees over evolutionary time. **f**, Evolvability is small (*η*_0_ is small) and decreases over evolutionary time. Evolvability equal to 0 here means no evolution despite selection; SI section S7; Eq. 1 of ref. ^83^). **g**, Population size increases over evolutionary time (plot of 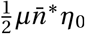, so the indicated multiplication yields population size). Mutation rate *µ* and parameter *η*_0_ can take any value satisfying 0 < *µ* 1 and 0 < *η*_0_ 1/(*N*_g_ *N*_a_), where the number of genotypic traits is *N*_g_ = 3 and the number of age bins is *N*_a_ = 47y/0.1y. If *µ* = 0.01 and *η*_0_ = 1/(3 × 47y/0.1y), then a population size of 1000×2/(*µη*_0_) is 2.82 billion individuals (which is unrealistically large due to the assumption of marginally small mutational variance to facilitate analysis). All plots are for the *sapiens* trajectory of Fig. 2b.

**Figure 5:**
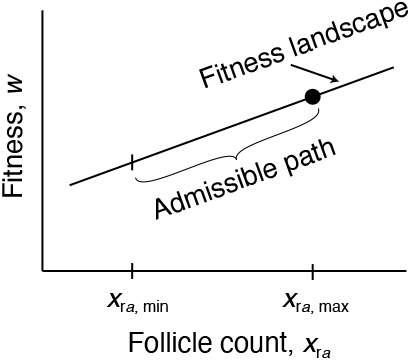
Illustration of the fitness landscape in the brain model. The fitness landscape *w* is a linear function (Eq. M3) of the follicle count 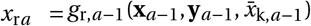, which is a recurrence over age. The slope of the fitness landscape with respect to *x*_r*a*_ is positive and decreases with age *a* (Fig. 4b). Evaluating the recurrence at all possible genotypic trait values 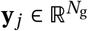 for all ages *j* < *a* gives values *x*_r*a*,min_ and *x*_r*a*,max_ that depend on development *g*_r *j*_ for all ages *j* < *a*, the various parameters influencing it, and the developmentally initial conditions. The admissible follicle count ranges from *x*_r*a*,min_ to *x*_r*a*,max_. The admissible path on the landscape is given by the admissible follicle count. As there are no absolute mutational constraints, evolution converges to the peak of the admissible path ^60^ (dot), where total genotypic selection vanishes, d*w* /d**y** = **0** (Extended Data Fig. 3).

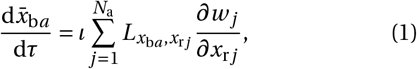

where *ι* is a non-negative scalar measuring mutational input, 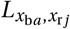 is the mechanistic additive socio-genetic covariance between brain size at age *a* and follicle count at age *j, w*_*j*_ is fitness at age *j*, and 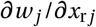 is the direct selection gradient of follicle count at age *j*. Eq. (1) shows that brain size evolves in the brain model because brain size is socio-genetically correlated with follicle count (i.e., setting the socio-genetic covariation between brain and follicle count to zero in Eq. 1, so 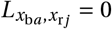 for all ages *a* and *j*, yields no brain size evolution).

Brain size and follicle count are socio-genetically correlated in the model because of development. To see this, consider the mechanistic additive socio-genetic cross-covariance matrix of the phenotype, given by

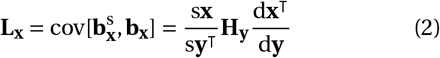

(L for legacy). Here **b**_**x**_ is the mechanistic breeding value of the phenotype and 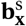 is the stabilised mechanistic breeding value, which is a generalisation of the former and considers the effects of social development. In turn, **H**_**y**_ is the mutational covariance matrix, d**x**^T^/d**y** is the matrix of total effects of the genotype on the phenotype, and s**x**/s**y**^T^ is the matrix of stabilised effects of the genotype on the phenotype, where stabilised effects are the total effects after social development has stabilised in the population. Whereas **H**_**y**_ depends on genotypic traits but not development, both d**x**^T^/d**y** and s**x**/s**y**^T^ depend on development. In the model, there is no mutational covariation (i.e., **H**_**y**_ is diagonal), so **H**_**y**_ does not generate socio-genetic covariation between brain size and follicle count. Hence, such socio-genetic covariation can only arise from the total and stabilised effects of the genotype on the phenotype, which arise from development.

Therefore, the various evolutionary outcomes matching the brain and body sizes of seven hominin species ^37^ (Fig. 1) arise in this model exclusively due to change in developmental constraints and not from change in direct selection on brain size or cognitive abilities. In the model, EETBs and the shape of EEE only directly affect the developmental map (**g**_*a*_) but not fitness, so varying EETBs and the shape of EEE does not affect the direction of direct selection, but only its magnitude (Eqs. S38). Moreover, from the equation that describes the long-term evolutionary dynamics (Eq. M5) it follows that varying EETBs and the shape of EEE only affects evolutionary outcomes (i.e., path peaks; Fig. 5) by affecting the mechanistic socio-genetic covariation **L**_**z**_ (Eq. S29). That sociogenetic covariation determines evolutionary outcomes despite no internal fitness landscape peaks is possible because there is socio-genetic covariation only along the admissible path where the developmental constraint is met (so **L**_**z**_ is singular ^59^) and consequently evolutionary outcomes occur at path peaks rather than landscape peaks ^60^ (Fig. 5). That is, the various evolutionary outcomes matching the brain and body sizes of seven hominin species ^37^ (Fig. 1) are exclusively due to change in mechanistic socio-genetic covariation described by the **L**_**z**_ matrix, by changing the position of path peaks on the peak-invariant fitness landscape. Therefore, ecology and possibly culture cause hominin brain expansion in the model by affecting developmental and consequently socio-genetic constraints rather than direct selection. Additionally, brain metabolic costs directly affect the developmental map (**g**_*a*_) and so affect mechanistic sociogenetic covariation (**L**_**z**_) but do not directly affect fitness (*w*) and so do not constitute direct fitness costs (Eqs. S8, S10, S2, S9, and M3). Yet, in the model, brain metabolic costs often constitute total fitness costs and, occasionally, total fitness benefits (Methods; Fig. S12; SI section S8).

Evolution is almost orthogonal to direct selection throughout hominin brain expansion in the model (Fig. 4e). Evolvability ^83^, measuring the extent to which evolution proceeds in the direction of direct selection, is ancestrally very small and decreases toward zero as evolution proceeds (Fig. 4f). This means evolution stops because there is no longer socio-genetic variation in the direction of direct selection. The population size expands as the brain expands (Fig. 4g), although it decreases when shifting from the *afarensis* scenario to the *sapiens* scenario due to the plastic change in phenotype (Fig. S7n).

### Analysis of the action of constraint

To gain further insight into what causes the recovered brain expansion, I now analyse the action of constraint. Since there is only direct selection for follicle count, the equation describing long-term evolution (Eq. M5) entails that whether or not a trait evolves in the model is dictated by whether or not there is mechanistic socio-genetic covariation between the trait and follicle count (e.g., Eq. 1).

Examination of such covariation shows that brain expansion in the model is caused by positive socio-genetic covariation between brain size and developmentally late follicle count. The mechanistic socio-genetic covariation of brain size with follicle count, and how such covariation evolves, is shown in Fig. 6. Ancestrally, socio-genetic covariation between brain size and developmentally early follicle count is negative (black area in Fig. 6a), but between brain size and developmentally late follicle count is slightly positive (orange area in Fig. 6a). This positive covariation is what causes brain expansion. This pattern of socio-genetic covariation is maintained as brain expansion proceeds, but developmentally early brain size becomes less socio-genetically covariant with follicle count and so stops evolving, whereas developmentally later brain size becomes socio-genetically covariant with increasingly developmentally later follicle count. The magnitude of covariation also evolves (Fig. 6a-d).

**Figure 6:**
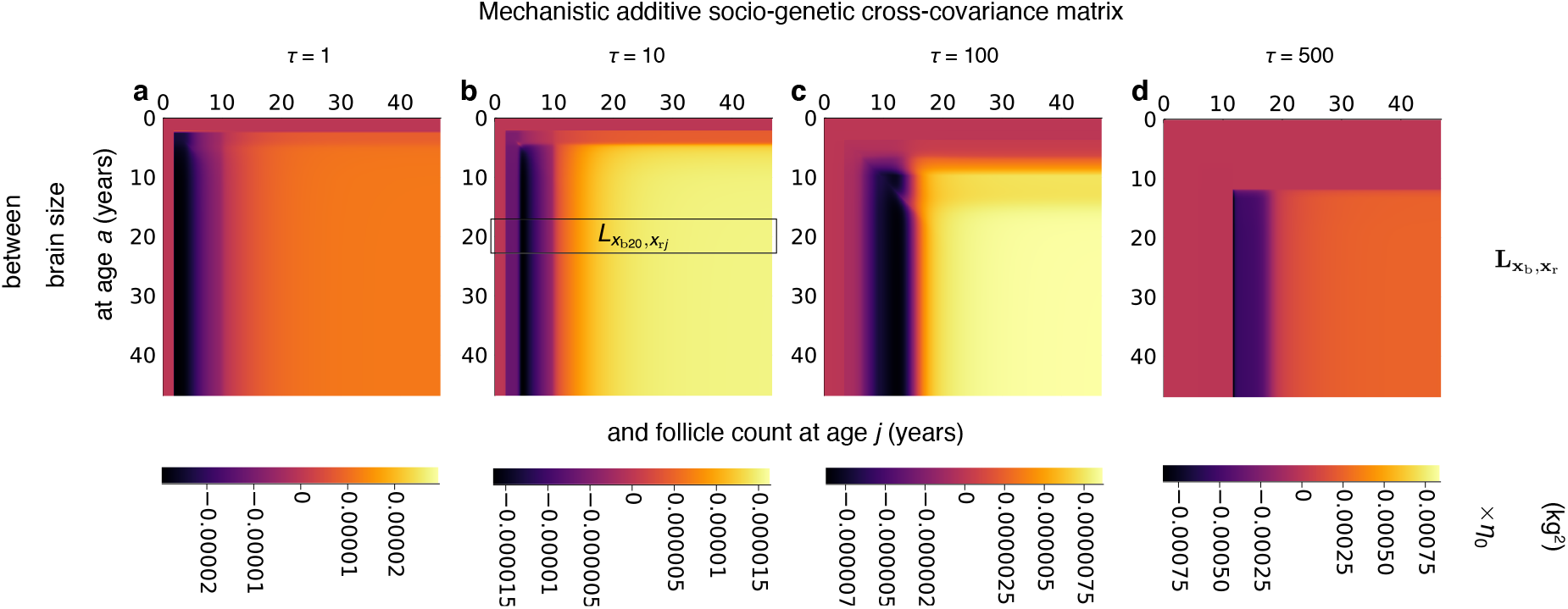
The action of constraint on brain expansion. Mechanistic socio-genetic cross-covariance matrix between brain size and follicle count over evolutionary time *τ*. For instance, in panel **b**, the highlighted box gives the sociogenetic covariance between brain size at 20 years of age and follicle count at each of the ages at the top horizontal axis. Thus, at evolutionary time *τ* = 10, socio-genetic covariation between brain size at 20 years of age and follicle count at 6 years of age is negative (bottom bar legend), but between brain size at 20 years of age and follicle count at 30 years of age is positive. The positive socio-genetic covariation between brain size and follicle count (e.g., yellow areas in **b** and **c**) causes brain expansion. Bar legends have different limits so that patterns are visible (bar legend limits are {−*l, l* }, where 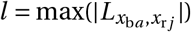 over *a* and *j* for each *τ*). All plots are for the *sapiens* trajectory of Fig. 2b.

Hence, direct selection on developmentally late follicle count provides a force for follicle count increase, and socio-genetic covariation between brain size and developmentally late follicle count diverts this force and causes brain expansion. This occurs even though the force of selection is weaker at advanced ages ^84^ (i.e., slopes are negative in Fig. 4b), which is compensated by developmentally increasing socio-genetic covariation with follicle count. Such covariation can arise because of developmental propagation of phenotypic effects of mutations ^60^. Therefore, ecology and culture cause brain expansion in the model by generating positive socio-genetic covariation over development between brain size and developmentally late follicle count.

The socio-genetic covariation between body size and follicle count, and between skill level and follicle count follow a similar pattern (Extended Data Fig. 4a-h). Hence, the evolutionary expansion in body size and skill level in the model are also caused by their positive socio-genetic covariation with developmentally late follicle count.

The evolution of follicle count is governed by a different pattern of socio-genetic covariation. Developmentally early follicle count evolves smaller values because of negative socio-genetic covariation with developmentally late follicle count (Extended Data Fig. 4i-l). In turn, developmentally late follicle count evolves higher values because of positive socio-genetic covariation with developmentally late follicle count (Extended Data Fig. 4i-l). Positive socio-genetic covariance between follicle count of different ages is clustered at the ages where follicle count developmentally increases most sharply (Fig. 3b). This cluster of positive socio-genetic covariation evolves to later ages (Extended Data Fig. 4j-l), corresponding to the evolved ages of peak developmental growth in follicle count (Fig. 3b). This cluster of positive socio-genetic covariation has little effect on follicle count evolution as follicle count around the evolving age of menarche mostly decreases over evolution, so such covariation is mostly compensated by the negative socio-genetic covariation with developmentally later follicle count. Socio-genetic covariation between other phenotypes exists (Figs. S10 and S11) but has no evolutionary effect as only that with follicle count does. Several of the above patterns of sociogenetic covariation emerge during the *afarensis* trajectory (Fig. S9).

## Discussion

I modelled the evo-devo dynamics of hominin brain expansion, recovering major patterns of human development and evolution. I showed that hominin brain expansion occurs in this model because brain size is socio-genetically correlated with developmentally late follicle count as there is only direct selection for follicle count. In other words, mutant alleles coding for increased allocation to brain growth can only increase in frequency in the model by being socio-genetically correlated with developmentally late follicle count which is selected for, rather than due to selection for brain size. This sociogenetic correlation is generated over development by a moderately challenging ecology and possibly cumulative culture. This covariation yields an admissible evolutionary path on the fitness landscape (Fig. 5), a path along which the brain expands because of developmental constraints, as without them there is no direct selection for or against brain expansion. Thus, in this model, hominin brain expansion is caused by unremarkable selection and particular developmental constraints involving a moderately challenging ecology and possibly cumulative culture. This constraint-caused brain expansion occurs despite it generating a brain-body allometry of 1.03 and a duplication of EQ (Extended Data Fig. 2g and Fig. 3g). While cognitive ability in the form of skill level is not directly under selection in the model, the model can be modified to incorporate such widely considered scenario. Yet, as found above, direct selection for cognitive ability is not necessary to recreate hominin brain expansion and multiple aspects of human development and evolution, whereas certain developmental constraints with unexceptional direct selection are sufficient, at least for the parameter values analysed. Change in development without changes in direct selection can thus yield diverse evolutionary outcomes, including the brain and body sizes of seven hominins, rather than only evolutionarily transient effects.

These results show that developmental constraints can play major evolutionary roles by causing hominin brain expansion in this in silico replica. Developmental constraints are traditionally seen as preventing evolutionary change ^12;85;86;52^, effectively without the ability to generate evolutionary change that is not already favoured by selection. Less prevalent views have highlighted the potential relevance of developmental constraints in evolution ^51^ and human brain evolution (e.g., p. 87 of ref. ^87^).

The findings here show that while constraints do prevent evolutionary change in some directions, constraints can be “generative” ^88^ in the sense that they can divert evolutionary change in a direction that causes brain expansion, such that without those constraints brain expansion is not favoured by selection and does not evolve.

The results above contrast with a previous study finding that direct selection on brain size drove brain expansion in hominins ^58^. Such a study used the short-term restricted Lande equation ^54;55^ for this long-term inference. I used analogous equations ^59^ that describe longterm evolution and that separate the evolutionary effects of developmental constraints and direct selection — a separation that has otherwise not been clear-cut ^57^.

My approach illustrates why the human brain size could have evolved, but it has not established why it did. Yet, this approach can be built upon to pursue that goal. There is scope for refinement of the model, for improved parameter estimates, and for other models to improve predictions as those obtained are near but do not exactly match observation, particularly in the ontogenetic patterns. Rapidly advancing techniques of simulation-based inference may then be used for model selection, parameter estimation, and uncertainty quantification ^31^. These techniques have been instrumental in multiple fields such as in the discovery of the Higgs boson ^31^ or in establishing that humans are causing climate change. Such simulation-based inference was impractical with the previous dynamic optimisation approach, as a single run took approximately 3 days ^37^, the runs are not easy to parallelise as suitable initial guesses are needed for the genotypic traits, and simulation-based inference needs hundreds of thousands of runs. This meant that simulation-based inference would have taken about 800 years. In contrast, a run here took approximately 4 minutes, indicating that simulation-based inference with the evo-devo dynamics approach could take months. This computational speed suggests that simulation-based inference ^31^ of human brain size evolution may now be feasible.

### Methods

#### Model overview

The evo-devo dynamics framework I use ^59^ is based on standard adaptive dynamics assumptions ^89;90^. The framework considers a resident, well-mixed, finite population with deterministic population dynamics where individuals can be of different ages, reproduction is clonal, and mutation is rare (mutants arise after previous mutants have fixed) and weak (mutant genotypes are marginally different from the resident genotype). Under these assumptions, population dynamics occur in a fast ecological timescale and evolutionary dynamics occur in a slow evolutionary timescale. Individuals have genotypic traits, collectively called the genotype, that are under direct genetic control. As mutation is weak, there is vanishingly small variation in genotypic traits (marginally small mutational variance). Also, individuals have phenotypic traits, collectively called the phenotype, that are developed, that is, constructed over life. A function **g**_*a*_, called the developmental map, describes how the phenotype is constructed over life and gives the developmental constraint. The developmental map can be non-linear, evolve, change over development, and take any differentiable form with respect to its arguments, but the phenotype at the initial age (here, newborns) is constant and does not evolve as is standard in life history theory. Mutant individuals of age *a* have fertility *f*_*a*_ (rate of offspring production) and survive to the next age with probability *p*_*a*_. The evo-devo dynamics framework provides equations describing the evolutionary dynamics of genotypic and phenotypic traits in gradient form, thus describing long-term genotypic and phenotypic evolution as the climbing of a fitness landscape while guaranteeing that the developmental constraint is met at all times.

The brain model ^36;37^ provides a specific developmental map **g**_*a*_, fertility *f*_*a*_, and survival *p*_*a*_, which can be fed into the evo-devo dynamics framework to model the evolutionary dynamics of the developed traits studied. More specifically, the brain model considers a female population, where each individual at each age has three tissue types — brain, reproductive, and remaining somatic tissues — and a skill level. Reproductive tissue is defined as referring to pre-ovulatory ovarian follicles, so that reproductive tissue is not involved in offspring maintenance, which allows for writing fertility as being proportional to follicle count (in mass units), in accordance to observation ^91^. As a first approximation, the brain model lets the survival probability at each age be constant. At each age, each individual has an energy budget per unit time, her resting metabolic rate *B*_rest_, that she uses to grow and maintain her tissues. The part of this energy budget used in growing her tissues is her growth metabolic rate *B*_syn_. A fraction of the energy consumed by the preovulatory follicles is for producing offspring, whereas a fraction of the energy consumed by the brain is for gaining (learning) and maintaining (memory) skills. Each individual’s skill level emerges from this energy bookkeeping rather than being assumed as given by brain size. Somatic tissue does not have a specific function but it affects body size, thus affecting the energy budget because of Kleiber’s law ^92^ which relates resting metabolic rate to body size by a power law. Genes control the individual’s energy allocation effort into producing brain tissue, preovulatory follicles, and somatic tissue at each age. The causal dependencies in the brain model are described in Extended Data Fig. 1, which uses the insights from the evo-devo dynamics framework, in particular, the separation of direct and total effects on fitness in the model.

I write the brain model with the notation of the evodevo dynamics framework as follows. The model considers four phenotypic traits (i.e., *N*_p_ = 4): brain mass, follicle count (in mass units), somatic tissue mass, and skill level at each age. For a mutant individual, the brain size at age *a* ∈ {1,…, *N*_a_} is *x*_b*a*_ (in kg), the follicle count at age *a* is *x*_r*a*_ (in kg), the size of the remaining somatic tissue at age *a* is *x*_s*a*_ (in kg), and the skill level at age *a* is *x*_k*a*_ (in terabytes, TB). The units of phenotypic traits (kg and TB) arise from the units of the parameters measuring the unit-specific metabolic costs of maintenance and growth of the respective trait. The vector **x**_*a*_ = (*x*_b*a*_, *x*_r*a*_, *x*_s*a*_, *x*_k*a*_)^T^ is the mutant phenotype at age *a*. Additionally, the model considers three genotypic traits (i.e., *N*_g_ = 3): the effort to produce brain tissue, preovulatory follicles, and somatic tissue at each age. For a mutant individual, the effort at age *a* to produce: brain tissue is *y*_b*a*_, follicles is *y*_r*a*_, and somatic tissue is *y*_s*a*_. These growth efforts are dimensionless and can be positive or negative, so they can be seen as measured as the difference from a baseline growth effort. The vector **y**_*a*_ = (*y*_b*a*_, *y*_r*a*_, *y*_s*a*_)^T^ is the mutant growth effort at age *a*, which describes the mutant genotypic traits at that age. The growth efforts generate the fraction *q*_*ia*_ (**y**_*a*_) of the growth metabolic rate *B*_syn_ allocated to growth of tissue *I* ∈ {b, r, s} at age *a* (*q*_*ia*_ corresponds to the control variables *u* in refs. ^36;37^; we consider *y*’s rather than *q*’s as the genotypic traits as the *y*’s do not need to be between zero and one nor add up to one, so numerical solution is simpler). To describe the evolutionary dynamics of the phenotype as the climbing of a fitness landscape, the evo-devo dynamics framework defines the mutant geno-phenotype at age *a* as the vector **z**_*a*_ = (**x**_*a*_; **y**_*a*_) (the semicolon indicates a linebreak). The mutant phenotype across ages is 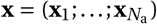, and similarly for the other variables. While **x**_*a*_ is a mutant’s phenotype *across traits* at age *a*, we denote the mutant’s *i* -th phenotype *across ages* as 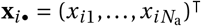 for *i* ∈ {b, r, s, k}. The mutant’s *i* -th genotypic trait across ages is 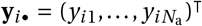 for *i* ∈ {b, r, s}. The resident traits are analogously denoted with an overbar (e.g., 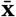).

The brain model describes development by providing equations describing the developmental dynamics of the phenotype. That is, the mutant phenotype at age *a* + 1 is given by the developmental constraint

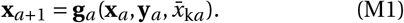

The equations for the developmental map **g**_*a*_ are given in section S1.1 of the SI and were previously derived from mechanistic considerations of energy conservation following the reasoning of West et al.’s metabolic model of ontogenetic growth ^38^ and phenomenological considerations of how skill relates to energy extraction ^36;37^. The developmental map of the brain model depends on the skill level of social partners of the same age (i.e., peers), 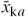, because of social challenges of energy extraction (where *P*_1_ < 1) so we say that development is social. When individuals face only ecological challenges (i.e., *P*_1_ = 1), development is not social.

The evo-devo dynamics are described by the developmental dynamics of the phenotypic traits given by Eq. (M1) and by the evolutionary dynamics of the genotypic traits. The latter are given by the canonical equation of adaptive dynamics ^89^

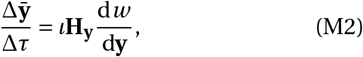

where *τ* is evolutionary time, *ι* is a non-negative scalar measuring mutational input and is proportional to the mutation rate and carrying capacity, and **H**_**y**_ = cov[**y, y**] is the mutational covariance matrix (H for heredity; derivatives are evaluated at resident trait values throughout and I use matrix calculus notation ^93^ as defined in Eq. S1). Due to age-structure, a mutant’s relative fitness is 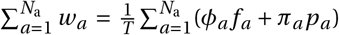 where *f*_*a*_ and *p*_*a*_ are a mutant’s fertility and survival probability at age *a, T* is generation time, and *φ*_*a*_ and *π*_*a*_ are the forces ^84;94;95^ of selection on fertility and survival at that age (*T, φ*_*a*_, and *π*_*a*_ are functions of the resident but not mutant trait values). After substitution and simplification, a mutant’s relative fitness reduces to

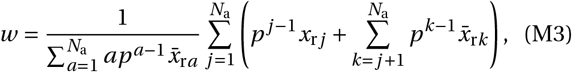

where *p* is the constant probability of surviving from one age to the next. This fitness function depends directly on the mutant’s follicle count, but only indirectly on metabolic costs via the developmental constraint (i.e., after substituting *x*_r *j*_ for the corresponding entry of Eq. M1). Eq. (M2) thus depends on the total selection gradient of genotypic traits d*w* /d**y**, which measures total genotypic selection. While Lande’s ^54^ selection gradient measures direct selection without considering developmental constraints by using partial derivatives (*∂*), total selection gradients measure selection considering developmental constraints by using total derivatives (d). Lande’s selection gradient thus measures the direction in which selection favours evolution to proceed without considering any constraint, whereas total selection gradients measure the direction in which selection favours evolution considering the developmental constraint (M1). From the first and last equalities in Layer 4, Eq. S21 of ref. ^59^, the total selection gradient of genotypic traits for the brain model is

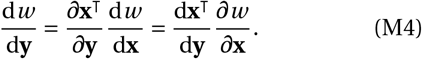

Eq. (M4) shows that total genotypic selection can be written in terms of either total phenotypic selection (d*w* /d**x**) or direct phenotypic selection (*∂w* /*∂***x**). Eqs. (M1) and (M2) together describe the evo-devo dynamics. Eq. (M2) entails that total genotypic selection vanishes at evolutionary equilibria if there are no absolute mutational constraints (i.e., if *ι* > 0 and **H**_**y**_ is non-singular). Moreover, since in the brain model there are more phenotypic traits than genotypic traits (*N*_p_ > *N*_g_), the matrices *∂***x**^T^/*∂***y** and d**x**^T^/d**y** have fewer rows than columns and so are singular; hence, setting Eq. (M4) to zero implies that evolutionary equilibria can occur with persistent direct and total phenotypic selection in the brain model.

While I use Eqs. (M1) and (M2) to compute the evodevo dynamics, those equations do not describe phenotypic evolution as the climbing of an adaptive topography. To analyse phenotypic evolution as the climbing of an adaptive topography, I use the following. The evodevo dynamics framework ^59^ shows that long-term phenotypic evolution can be understood as the climbing of a fitness landscape by simultaneously following genotypic and phenotypic evolution, which for the brain model is given by

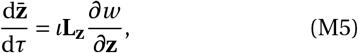

since **z** = (**x**; **y**) includes the phenotype **x** and genotypic traits **y**. The vector *∂w* /*∂***z** is the direct selection gradient of the geno-phenotype (as in Lande’s ^54^ selection gradient of the phenotype). The matrix **L**_**z**_ is the mechanistic additive socio-genetic cross-covariance matrix of the geno-phenotype, for which the evo-devo dynamics framework provides formulas that guarantee that the developmental constraint (M1) is met at all times. The matrix **L**_**z**_ is asymmetric due to social development; if individuals face only ecological challenges, development is not social and **L**_**z**_ reduces to **H**_**z**_, the mechanistic additive genetic covariance matrix of the geno-phenotype, which is symmetric (**H**_**x**_ is a mechanistic version of Lande’s ^54^ **G** matrix: whereas **H**_**x**_ involves total derivatives describing the total effect of genotype on phenotype, **G** is defined in terms of regression of phenotype on genotype; hence, **H**_**x**_ and **G** have different properties including that mechanistic heritability can be greater than one). The matrix **L**_**z**_ is always singular because it considers both the phenotype and genotypic traits, so selection and development jointly define the evolutionary outcomes even with a single fitness peak ^60^. Eq. (M5) and the formulas for **L**_**z**_ entail that evolution proceeds as the climbing of the fitness landscape in geno-phenotype space, where the developmental constraint (M1) provides the admissible evolutionary path, such that evolutionary outcomes occur at path peaks rather than landscape peaks if there are no absolute mutational constraints ^60^.

I implement the developmental map of the brain model into the evo-devo dynamics framework to study the evolutionary dynamics of the resident phenotype 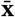, including the resident brain size 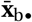.

#### Seven hominin scenarios

It was previously found ^37^ that, at evolutionary equilibrium, the brain model recovers the evolution of the adult brain and body sizes of six *Homo* species and less accurately of *Australopithecus afarensis*. The parameter values yielding these seven outcomes are described in Fig. 1. I call each such parameter combination a scenario. The *sapiens, neardenthalensis*, and *heidelbergensis* scenarios use weakly diminishing returns of learning and submultiplicative cooperation: specifically, these scenarios use exponential competence with parameter values given in Regime 1 of Table S2 (Eq. S5). I call ecological scenario that with such weakly diminishing returns of learning and submultiplicative cooperation but setting the proportion of ecological challenges to one (*P*_1_ = 1), which was previously ^36^ found to yield the evolution of brain and body sizes of Neanderthal scale at evolutionary equilibrium. The *erectus, ergaster*, and *habilis* scenarios use strongly diminishing returns of learning and additive cooperation: specifically, these scenarios use power competence with parameter values given in Regime 2 of Table S2 and with additive cooperation (Eq. S5). The *afarensis* scenario uses strongly diminishing returns of learning and submultiplicative cooperation; that is, power competence with parameter values given in Regime 2 of Table S2 (Eq. S5). In the main text, I primarily describe results under the *sapiens* scenario. In the SI, I also give analogous results under the *afarensis* (Figs. S1, S7, and S9) and ecological (Fig. S5) scenarios.

#### Ancestral genotypic traits

To solve the evo-devo dynamics, one must specify the ancestral resident genotypic traits giving the resident growth efforts 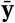 at the initial evolutionary time. I explored nine sets of ancestral genotypes described in SI section S4 and labelled as: naive, somewhatNaive, ecoSols, highlySpecified, afarensisFromHighlySpecified, afarensisFromSomewhatNaive, somewhatNaive2, afarensisFromNaive2, and afarensisFromEcoSols. I find that the outcome depends on the ancestral genotype. For instance, in the *sapiens* scenario, at least two drastically different evolutionary outcomes are possible by changing only the ancestral genotype (i.e., there is bistability in brain size evolution), so there are at least two path peaks on the fitness landscape as follows. Using somewhatNaive ancestral growth efforts in the *sapiens* scenario yields an evolutionary outcome with no brain, where residents have a somewhat semelparous lifehistory reproducing for a short period early in life followed by body shrinkage (Fig. S4). In contrast, using afarensisFromNaive2 ancestral growth efforts in the *sapiens* scenario yields adult brain and body sizes of *H. sapiens* scale (Fig. 3). This bistability does not arise under the ecological scenario which yields brain expansion under somewhatNaive ancestral growth efforts (Fig. S5). Thus, for the *sapiens* scenario to yield brain and body sizes of *H. sapiens* scale it seems to require an ancestral genotype that can develop large bodies under cooperative and between-group competitive social challenges (blue dots in Figs. 3c and S6o), whereas ancestrally large bodies are not needed for brain expansion under purely ecological challenges (blue dots in Fig. S5o). In the main text, I present the results for the *sapiens* scenario with the afarensisFromNaive2 ancestral genotype.

#### The action of total selection

Despite absence of direct selection on brain size or skill level in the model, there is total selection on the various traits. Total selection is measured by total selection gradients that quantify the total effect of a trait on fitness considering the developmental constraints and so how traits affect each other over development ^59;96^. Thus, in contrast to direct selection, total selection confounds the action of selection and constraint. Since I assume there are no absolute mutational constraints (i.e., **H**_**y**_ is non-singular), evolutionary outcomes occur at path peaks in the fitness landscape where total genotypic selection vanishes (d*w* /d**y** = **0**), which are not necessarily fitness landscape peaks where direct selection vanishes (*∂w* /*∂***z** /= **0**).

The following patterns of total selection occur during the *sapiens* trajectory of Fig. 2b. Total selection ancestrally favours increased brain size throughout life (blue circles in Extended Data Fig. 3a). As evolution advances, total selection for brain size decreases and becomes negative early in life, possibly due to my assumption that the brain size of a newborn is fixed and cannot evolve. A similar pattern results for total selection on follicle count (Extended Data Fig. 3b). Somatic tissue is ancestrally totally selected for early in life and against later in life, eventually becoming totally selected for throughout life (Extended Data Fig. 3c). Total selection for skill level ancestrally fluctuates but decreases across life, decreasing but remaining positive throughout life as evolution proceeds (Extended Data Fig. 3d). Thus, total selection still favours evolutionary change in the phenotype at evolutionary equilibrium, but change is no longer possible (red dots in Extended Data Fig. 3a-d are at non-zero values). This means that evolution does not and cannot reach the favoured total level of phenotypic change in the model.

Although evolution does not reach the favoured total level of *phenotypic* change in the model, it does reach the favoured total level of *genotypic* change because of the assumption of no absolute mutational constraints. Total selection for the genotypic trait of brain growth effort is ancestrally positive early in life and evolves toward zero (Extended Data Fig. 3e). Total genotypic selection for follicle production is also ancestrally positive early in life, transiently evolves to negative, and eventually approaches zero (Extended Data Fig. 3f). Total genotypic selection for somatic growth effort is ancestrally negative early in life and evolves toward zero (Extended Data Fig. 3g). The evolved lack of total genotypic selection means that evolution approaches the favoured total level of genotypic change. This also means that evolution stops at a path peak on the fitness landscape (Fig. 5).

The occurrence of total selection for brain size or skill level may suggest that this total selection causes brain expansion in the model, but in the recovered brain expansion total selection can change the evolved brain size only due to change in the developmental constraints. This is because total selection equals the product of direct selection and total developmental bias (Eqs. M4 and S34), and in the model changing EETBs or the shape of EEE does not affect the direction of direct selection but only the direction of total developmental bias by affecting the developmental constraints. Thus, varying EETBs or the shape of EEE affects total selection in the evolved brain and body sizes only because the developmental constraints are changed rather than direct selection.

#### Total fitness effects of metabolic costs

While brain metabolic costs do not entail direct fitness costs in the model (i.e., *∂w* /*∂B*_b_ = 0), they may entail total fitness costs (i.e., d*w* /d*B*_b_ /= 0) and these can be computed using formulas from the evo-devo dynamics framework (SI section S8). Using these formulas shows that metabolic costs of maintenance may be total fitness costs at some ages but benefits at some other ages over the *sapiens* trajectory (Fig. S12). Consequently, the metabolic cost of brain maintenance is a total fitness benefit at evolutionary time 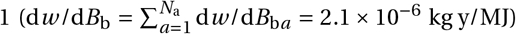, and a total fitness cost at evolutionary times 10, 100, and 500 (−1.5 × 10^−5^ kg y/MJ, −2 × 10^−5^ kg y/MJ, and −1.7 × 10^−5^ kg y/MJ, respectively). Moreover, among tissues, the metabolic cost of somatic maintenance (**i**-**l**) has some of the most substantial total fitness effects even tough it is the smallest metabolic cost of maintenance (Fig. S12), perhaps due to the large size of somatic tissue. Total fitness costs also confound the action of selection and constraint as they depend on development rather than only on selection. That is, total fitness costs share components with genetic covariation.

## Data availability

No data was collected in this study. All data used were previously published in references provided in the main text or supplementary information.

### Code availability

All code is available in the supplementary information and has been deposited in ref. ^62^.

## Acknowledgments

I thank A. Gardner, K. Laland, R. Patchett, and C.R. Turner for comments on previous versions of the manuscript, A. Gardner for funding, S.D. Healy and C. Rutz for discussion, and R.I.M. Dunbar and 3 anonymous reviewers for detailed comments that helped to substantially improve the manuscript. A. Gardner suggested to randomly sample genotypic traits to evaluate the resulting brain-body allometry as in Fig. 2a. This work was funded by an European Research Council Consolidator Grant to A. Gardner (grant no. 771387).

## Author contributions

MGF conceived and carried out the research and wrote the paper.

---

**Extended Data Figure 1:**
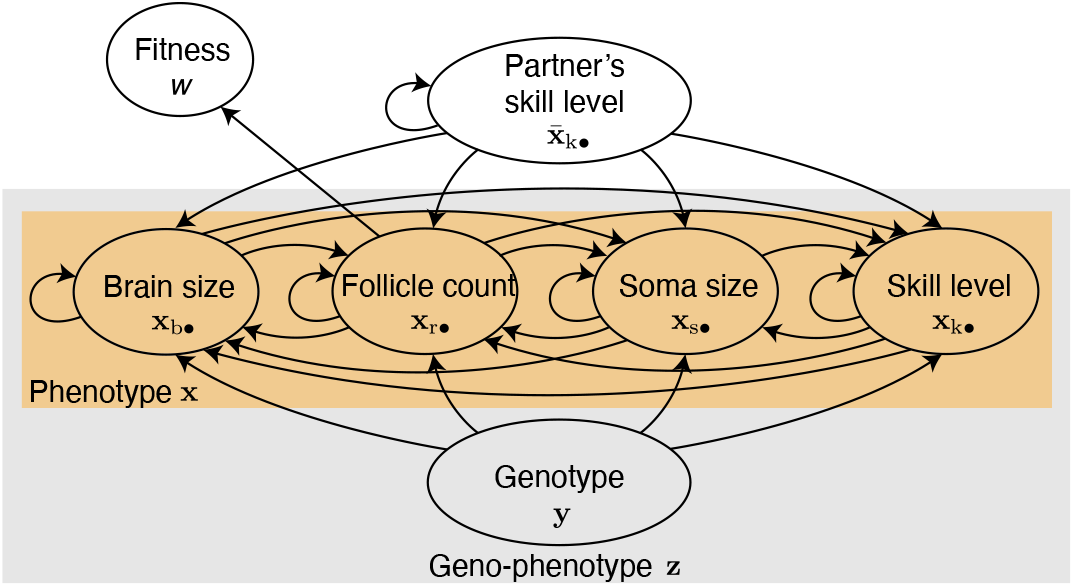
Causal diagram of the brain model analysed under the evo-devo dynamics framework. The evo-devo dynamics framework clarifies how to separate the direct and total effects of traits on fitness in the model. Variables have age-specific values that are not shown for clarity. The phenotype comprises brain size, follicle count, somatic tissue size, and skill level, all of which are constructed by a developmental process. Each arrow indicates the direct effect of a variable on another one. The total effect of a variable on another one is that across all the arrows directly or indirectly connecting the former to the latter. A mutant’s genotypic traits at a given age directly affect brain size, follicle count, somatic tissue size, and skill level at the immediately subsequent age (with the slope quantifying developmental bias from genotype). A mutant’s phenotypic traits at a given age affect themselves at the immediately subsequent age (quantifying developmental bias from the same phenotypic trait), thus the direct feedback loop from phenotypic traits to themselves. A mutant’s phenotypic traits at a given age also directly affect each other at the next age (quantifying developmental bias from immediately previous phenotypes). A mutant’s follicle count is the only trait directly affecting fitness (direct selection on follicle count). The social partner’s skill level at a given age directly affects its own development at an immediately subsequent age (quantifying developmental bias from the same phenotypic trait), thus the direct feedback loop. The social partner’s skill level at a given age also directly affects all the mutant’s phenotypic traits at the next age (quantifying indirect genetic effects from the phenotype). The genotype is assumed to be developmentally independent (i.e., controls **y** are open-loop), which means that there is no arrow towards the genotype. This diagram is a simplification of that considered by the evo-devo dynamics framework ^59^, so the brain model can be extended and the framework can still be used to analyse it.

**Extended Data Figure 2:**
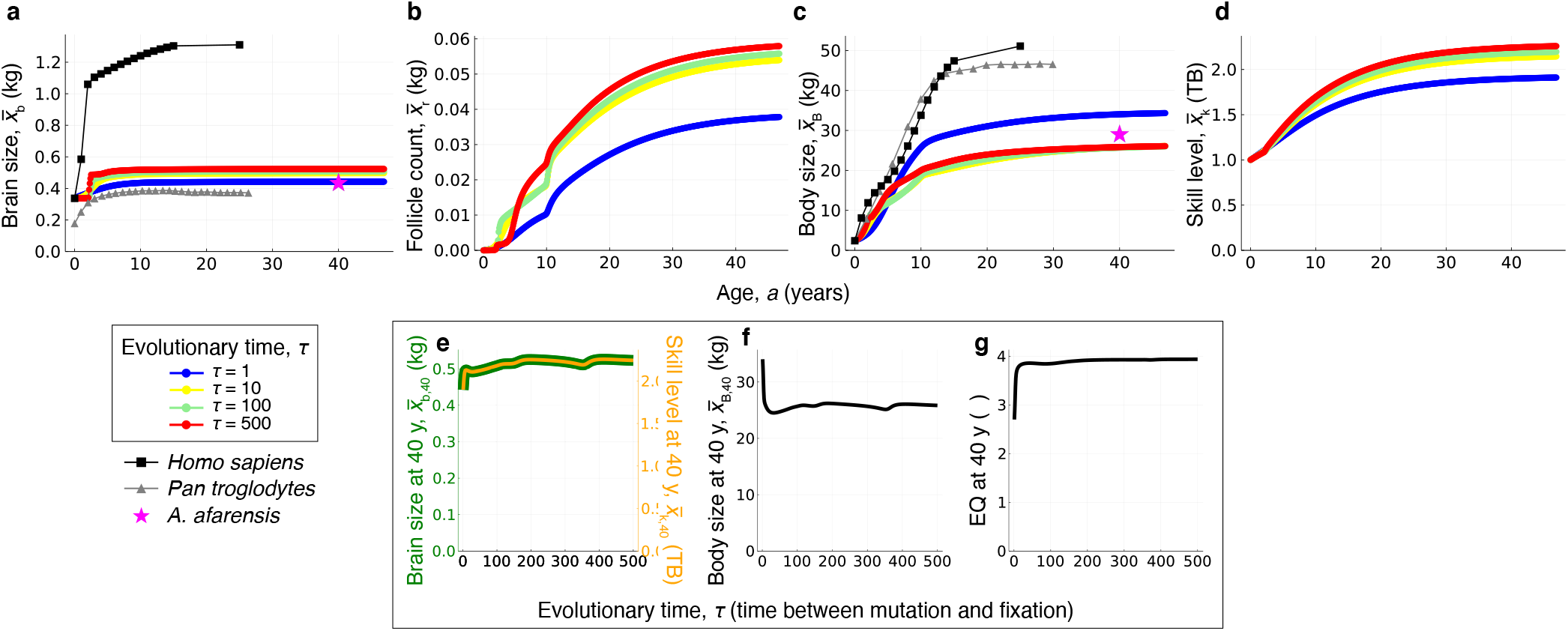
Evo-devo dynamics of brain size under *afarensis* scenario. Developmental dynamics are over age (e.g., horizontal axis in **a**) and evolutionary dynamics are over evolutionary time (differently coloured dots; center left label). Evo-devo dynamics of: **a**, brain size; **b**, follicle count; **c**, somatic tissue size; **d**, skill level; **e**, body size; and **f**, the yearly weight velocity showing the evolution of two growth spurts. Evolutionary dynamics of (**g**) brain size (green), (**g**) skill level (orange), (**h**) body size, and (**i**) encephalisation quotient (EQ) at 40 years of age. **a**,**e**,**f**, The mean observed values in a modern human female sample are shown in black squares (data from ref. ^39^ who fitted data from ref. ^69^). One evolutionary time unit is the time from mutation to fixation.

**Extended Data Figure 3:**
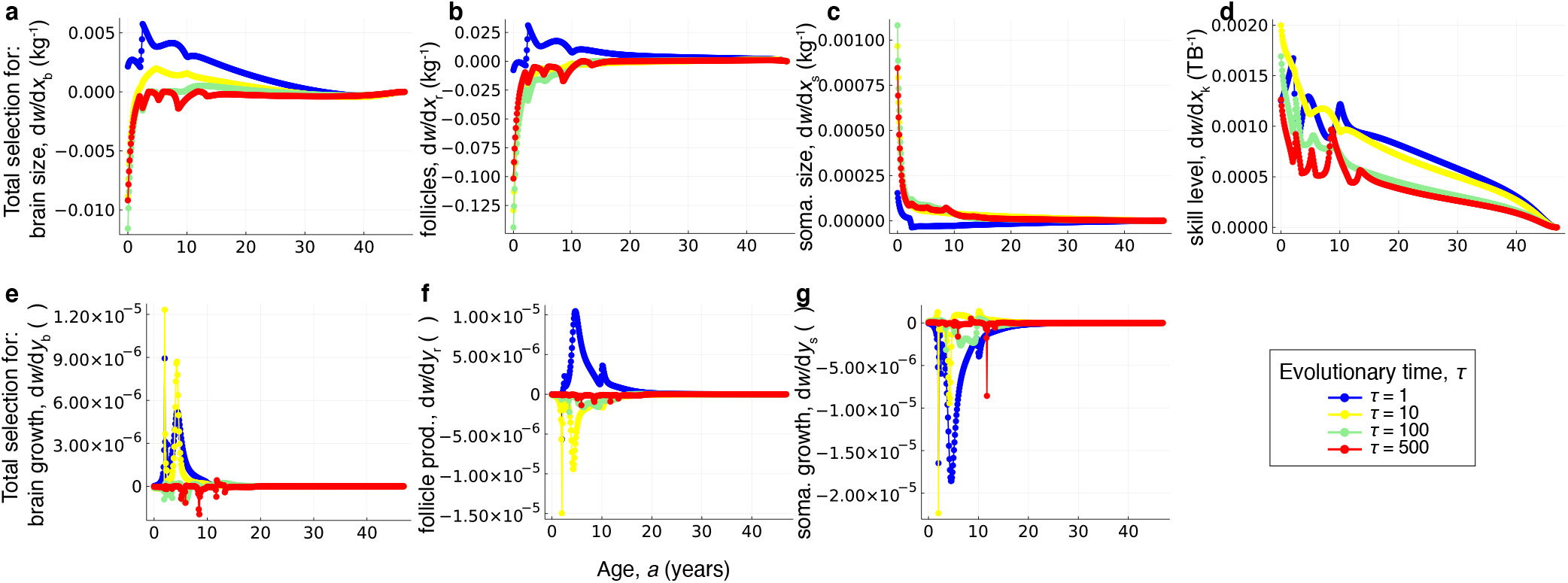
The action of total selection. **a-d**, Total selection on brain size, follicle count, somatic tissue size, and skill level at each age over evolutionary time. Total selection for skill level over life persists at evolutionary equilibrium (red dots in **d**). **e-g**, Total selection on effort for brain growth, follicle production, and somatic growth at each age over evolutionary time. Total selection for genotypic traits nearly vanish at evolutionary equilibrium (red dots in **e-g**), indicating that a path peak on the fitness landscape is reached. All plots are for the *sapiens* trajectory of Fig. 2b.

**Extended Data Figure 4:**
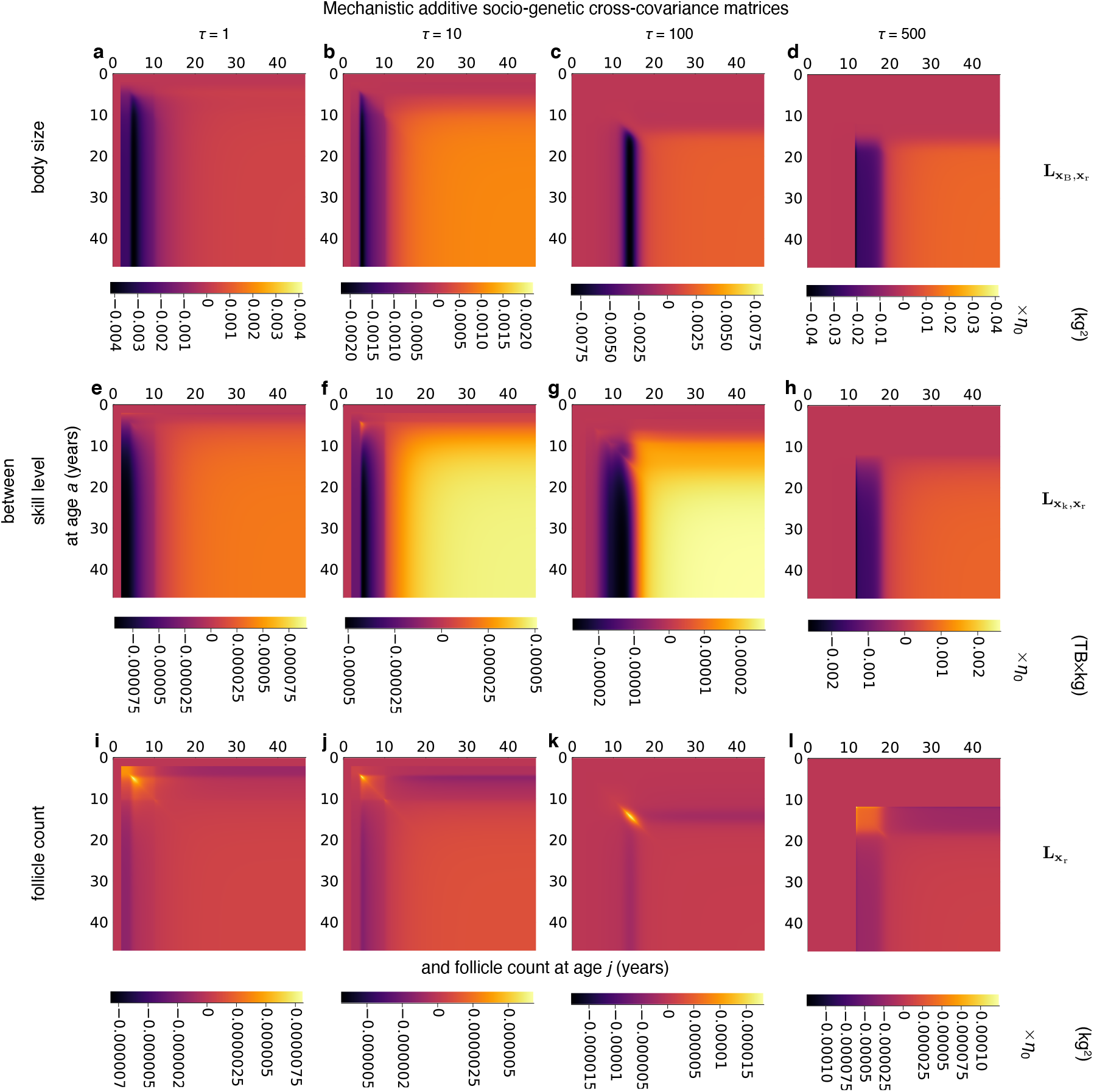
The action of constraint on body, skill, and follicle count expansion. Mechanistic sociogenetic cross-covariance matrix between: **a-d**, body size and follicle count, **e-h**, skill level and follicle count, and **i-l**, follicle count and itself. All plots are for the *sapiens* trajectory of Fig. 2b.

## References

1. Klein, R. G. The Human Career (The Univ. of Chicago Press, 2009), 3rd edn.

2. Brown, P. et al. A new small-bodied hominin from the Late Pleistocene of Flores, Indonesia. Nature 431, 1055–1061 (2004).

3. Garvin, H. M. et al. Body size, brain size, and sexual dimorphism in Homo naledi from the Dinaledi Chamber. J. Hum. Evol. 111, 119–138 (2017).

4. Darwin, C. The Descent of Man (John Murray, London, UK, 1871).

5. Jolly, A. Lemur social behavior and primate intelligence. Science 153, 501–506 (1966).

6. Humphrey, N. K. The social function of the intellect. In Bateson, P. P. G. & Hinde R. A. (eds.) Growing Points in Ethology, 303–317 (Cambridge Univ. Press, 1976).

7. Clutton-Brock, T. H. & Harvey, P. H. Primates, brains and ecology. J. Zool. 190, 309–323 (1980).

8. Hill, K. Hunting and human evolution. J. Hum. Evol. 11, 521–544 (1982).

9. Byrne, R. & Whiten A. (eds.) Machiavellian Intelligence (Oxford Univ. Press, 1988).

10. Alexander, R. How did humans evolve? Reflections on the uniquely unique species (University of Michigan Museum of Zoology, Ann Arbor, MI, 1990).

11. Aiello, L. C. & Wheeler, P. The expensive-tissue hypothesis. Curr. Anthropol. 36, 199–221 (1995).

12. Dunbar, R. I. M. The social brain hypothesis. Evol. Anthropol. 6, 178–190 (1998).

13. Potts, R. Variability selection in hominid evolution. Evol. Anthropol. 7, 81–96 (1998).

14. Wrangham, R. W., Jones, J. H., Laden, G., Pilbeam, D. & Conklin-Brittain, N. The raw and the stolen: cooking and the ecology of human origins. Curr. Anthropol. 40, 567–594 (1999).

15. Kaplan, H., Hill, K., Lancaster, J. & Hurtado, A. M. A theory of human life history evolution: diet, intelligence, and longevity. Evol. Anthropol. 9, 156–185 (2000).

16. Flinn, M. V., Geary, D. C. & Ward, C. V. Ecological dominance, social competition, and coalitionary arms races: Why humans evolved extraordinary intelligence. Evol. Hum. Behav. 26, 10–46 (2005).

17. Isler, K. & van Schaik, C. P. The expensive brain: a framework for explaining evolutionary changes in brain size. J. Hum. Evol. 57, 392–400 (2009).

18. van Schaik, C. P. & Burkart, J. M. Social learning and evolution: the cultural intelligence hypothesis. Phil. Trans. R. Soc. B 366, 1008–1016 (2011).

19. Henrich, J. The Secret of our Success (Princeton Univ. Press, 2016).

20. Dunbar, R. I. M. & Shultz, S. Why are there so many explanations for primate brain evolution? Phil. Trans. R. Soc. B 20160244 (2017).

21. Laland, K. N. Darwin’s Unfinished Symphony (Princeton Univ. Press, 2017).

22. Rosati, A. G. Foraging cognition: reviving the ecological intelligence hypothesis. Trends Cogn Sci 21, 691–702 (2017).

23. Will, M., Krapp, M., Stock, J. T. & Manica, A. Different environmental variables predict body size and brain size evolution in Homo. Nat. Comm. 12, 4116 (2021).

24. DeCasien, A. R., Barton, R. A. & Higham, J. P. Understanding the human brain: insights from comparative biology. Trends Cogn. Sci. 26, 432–445 (2022).

25. Lanuza, J. B., Collado, M. A., Sayol, F., Sol, D. & Bartomeus, I. Brain size predicts bees’ tolerance to urban environments. Biol. Lett. 19, 20230296 (2023).

26. Kotrschal, A. et al. Artificial selection on relative-brain size in the guppy reveals costs and benefits of evolving a larger brain. Curr. Biol. 23, 168–171 (2013).

27. Whiten, A. & van de Waal, E. Social learning, culture and the ‘socio-cultural brain’ of human and non-human primates. Neurosci. Biobehav. Rev. 82, 58–75 (2017).

28. Healy, S. D. Adaptation and the Brain (Oxford Univ. Press, Oxford, UK, 2021).

29. Hooper, R., Brett, B. & Thornton, A. Problems with using comparative analyses of avian brain size to test hypotheses of cognitive evolution. PLoS ONE 17, e0270771 (2022).

30. Pearl, J. Causality (Cambridge Univ. Press, 2009), 2nd edn.

31. Cranmer, K., Brehmer, J. & Louppe, G. The frontier of simulation-based inference. Proc. Natl. Acad. Sci. USA 117, 30055–30062 (2020).

32. Kaplan, H. & Robson, A. J. The emergence of humans: the coevolution of intelligence and longevity with intergenerational transfers. Proc. Natl. Acad. Sci. USA 99, 10221–10226 (2002).

33. Gavrilets, S. & Vose, A. The dynamics of Machiavellian intelligence. Proc. Natl. Acad. Sci. USA 103, 16823–16828 (2006).

34. dos Santos, M. & West, S. A. The coevolution of cooperation and cognition in humans. Proc. R. Soc. B 285, 20180723 (2018).

35. Muthukrishna, M., Doebeli, M., Chudek, M. & Henrich, J. The cultural brain hypothesis: how culture drives brain expansion, sociality, and life history. PLOS Comp. Biol. 14, e1006504 (2018).

36. González-Forero, M., Faulwasser, T. & Lehmann, L. A model for brain life history evolution. PLOS Comp. Biol. 13, e1005380 (2017).

37. González-Forero, M. & Gardner, A. Inference of ecological and social drivers of human brain-size evolution. Nature 557, 554–557 (2018).

38. West, G. B., Brown, J. H. & Enquist, B. J. A general model for ontogenetic growth. Nature 413, 628–631 (2001).

39. Kuzawa, C. W. et al. Metabolic costs and evolutionary implications of human brain development. Proc. Nat. Acad. Sci. USA 111, 13010–13015 (2014).

40. Nijhout, H. F. & Paulsen, S. M. Developmental models and polygenic characters. Am. Nat. 149, 394–405 (1997).

41. Pigliucci, M. Do we need an extended evolutionary synthesis? Evolution 61, 2743–2749 (2007).

42. Müller, G. B. Why an extended evolutionary synthesis is necessary. Interface Focus 7, 20170015 (2017).

43. Schaffer, W. M. The application of optimal control theory to the general life history problem. Am. Nat. 121, 418–431 (1983).

44. Roff, D. A. The Evolution of Life Histories (Chapman & Hall, New York, NY, USA, 1992).

45. Sydsæter, K., Hammond, P., Seierstad, A. & Strom, A. Further Mathematics for Economic Analysis (Prentice Hall, 2008), 2nd edn.

46. Kamien, M. I. & Schwartz, N. L. Dynamic Optimization (Dover, Mineola, NY, 2012), 2nd edn.

47. Dieckmann, U., Heino, M. & Parvinen, K. The adaptive dynamics of function-valued traits. J. Theor. Biol. 241, 370–389 (2006).

48. Parvinen, K., Heino, M. & Dieckmann, U. Function-valued adaptive dynamics and optimal control theory. J. Math. Biol. 67, 509–533 (2013).

49. Jerison, H. J. Evolution of the Brain and Intelligence (Academic Press, 1973).

50. Pilbeam, D. & Gould, S. J. Size and scaling in human evolution. Science 186, 892–901 (1974).

51. Gould, S. J. & Lewontin, R. C. The spandrels of San Marco and the Panglossian paradigm: a critique of the adaptationist programme. Proc. R. Soc. Lond. B 205, 581–598 (1979).

52. Montgomery, S. H., Mundy, N. I. & Barton, R. A. Brain evolution and development: adaptation, allometry and constraint. Proc. R. Soc. B 283, 20160433 (2016).

53. Tsuboi, M. et al. Breakdown of brain-body allometry and the encephalization of birds and mammals. Nat. Ecol. Evol. 2, 1492–1500 (2018).

54. Lande, R. Quantitative genetic analysis of multivariate evolution applied to brain: body size allometry. Evolution 34, 402–416 (1979).

55. Walsh, B. & Lynch, M. Evolution and Selection of Quantitative Traits (Oxford Univ. Press, Oxford, UK, 2018).

56. Brakefield, P. M. Evo-devo and constraints on selection. Trends Ecol. Evol. 21, 362–368 (2006).

57. Uller, T., Moczek, A. P., Watson, R. A., Brakefield, P. M. & Laland, K. N. Developmental bias and evolution: A regulatory network perspective. Genetics 209, 949–966 (2018).

58. Grabowski, M. Bigger brains led to bigger bodies?: the correlated evolution of human brain and body size. Curr. Anthropol. 57, 174–185 (2016).

59. González-Forero, M. A mathematical framework for evo-devo dynamics. Theor. Popul. Biol. 155, 24–50 (2024).

60. González-Forero, M. How development affects evolution. Evolution 77, 562–579 (2023).

61. Moore, A. J., Brodie III, E. D. & Wolf, J. B. Interacting phenotypes and the evolutionary process: I. direct and indirect genetic effects of social interactions. Evolution 51, 1352–1362 (1997).

62. González-Forero, M. Computer code for: Evo-devo dynamics of hominin brain size (3.0). Zenodo 10.5281/zenodo.10479516 (2024).

63. Bezanson, J., Edelman, A., Karpinski, S. & Shah, V. B. Julia: A fresh approach to numerical computing. SIAM Review 59, 65–98 (2017).

64. Smaers, J. B. et al. The evolution of mammalian brain size. Sci. Adv. 7, eabe2101 (2021).

65. Waddington, C. H. Canalization of development and the inheritance of acquired characters. Nature 150, 563–565 (1942).

66. Nijhout, H. F. & German, R. Z. Developmental causes of allometry: New models and implications for phenotypic plasticity and evolution. Integr. Comp. Biol. 52, 43–52 (2012).

67. Hansen, T. F. Stabilizing selection and the comparative analysis of adaptation. Evolution 51, 1341–1351 (1997).

68. Kirkpatrick, M. & Lofsvold, D. Measuring selection and constraint in the evolution of growth. Evolution 46, 954–971 (1992).

69. Dekaban, A. S. & Sadowsky, D. Changes in brain weights during the span of human life: Relation of brain weights to body heights and body weights. Ann. Neurol. 4, 345–356 (1978).

70. Leigh, S. R. Brain growth, life history, and cognition in primate and human evolution. Am. J. Primatol. 62, 139–164 (2004).

71. Bogin, B. Patterns of Human Growth (Cambridge Univ. Press, Cambridge, UK, 1988).

72. Bogin, B. & Smith, B. H. Evolution of the human life cycle. Am. J. Hum. Biol. 8, 703–716 (1996).

73. Gluckman, P. D. & Hanson, M. A. Evolution, development and timing of puberty. Trends Endocrinol. Metab. 17, 7–12 (2006).

74. Robson, S. L. & Wood, B. Hominin life history: reconstruction and evolution. J. Anat. 212, 394–425 (2008).

75. Jones, J. H. Primates and the evolution of long, slow life histories. Curr. Biol. 21, R708–R717 (2011).

76. Leigh, S. R. Evolution of human growth. Evol. Anthropol. 10, 223–236 (2001).

77. Tanner, J. M., Whitehouse, R. H. & Takaishi, M. Standards from birth to maturity for height, weight, height velocity, and weight velocity: British children, 1965. Arch. Dis. Childh. 41, 454–471 (1966).

78. Cameron, N. & Schell, L. M. (eds.) Human Growth and Development (Academic Press, London, UK, 2022), 3rd edn.

79. Howarth, C., Peppiatt-Wildman, C. M. & Attwell, D. The energy use associated with neural computation in the cerebellum. J. Cereb. Blood Flow Metab. 30, 403–414 (2010).

80. Pontzer, H. et al. Daily energy expenditure through the human life course. Science 373, 808–812 (2021).

81. Martin, R. D. Relative brain size and basal metabolic rate in terrestrial vertebrates. Nature 293, 57–60 (1981).

82. Leigh, S. R. & Shea, B. T. Ontogeny and the evolution of adult body size dimorphism in apes. Am. J. Primatol. 36, 37–60 (1995).

83. Hansen, T. F. & Houle, D. Measuring and comparing evolvability and constraint in multivariate characters. J. Evol. Biol. 21, 1201–1219 (2008).

84. Hamilton, W. D. The moulding of senescence by natural selection. J. Theor. Biol. 12, 12–45 (1966).

85. Maynard Smith, J. et al. Developmental constraints and evolution. Q. Rev. Biol. 60, 265–287 (1985).

86. Beldade, P., Koops, K. & Brakefield, P. M. Developmental constraints versus flexibility in morphological evolution. Nature 416, 844–847 (2002).

87. Gould, S. J. The Structure of Evolutionary Theory (Belknap Press, Cambridge, MA, USA, 2002).

88. Laland, K. N. et al. The extended evolutionary synthesis: its structure, assumptions and predictions. Proc. R. Soc. B 282, 20151019 (2015).

89. Dieckmann, U. & Law, R. The dynamical theory of coevolution: a derivation from stochastic ecological processes. J. Math. Biol. 34, 579–612 (1996).

90. Metz, J., Geritz, S., Meszéna, G., Jacobs, F. & van Heerwaarden, J. Adaptive dynamics, a geometrical study of the consequences of nearly faithful reproduction. In van Strien, S. & Lunel S. V. (eds.) Stochastic and spatial structures of dynamical systems, 183–231 (Konink. Nederl. Akad. Wetensch. Verh. Afd. Natuurk. Eerste Reeks, Amsterdam, Netherlands, 1996).

91. Broekmans, F. J., Faddy, M. J., Scheffer, G. & te Velde, E. R. Antral follicle counts are related to age at natural fertility loss and age at menopause. Menopause 11, 607–614 (2004).

92. Kleiber, M. The Fire of Life (Wiley, 1961).

93. Caswell, H. Sensitivity Analysis: Matrix Methods in Demography and Ecology (Springer Open, Cham, Switzerland, 2019).

94. Caswell, H. A general formula for the sensitivity of population growth rate to changes in life history parameters. Theor. Popul. Biol. 14, 215–230 (1978).

95. Baudisch, A. Hamilton’s indicators of the force of selection. Proc. Natl. Acad. Sci. USA 102, 8263–8268 (2005).

96. Morrissey, M. B. Selection and evolution of causally covarying traits. Evolution 68, 1748–1761 (2014).

97. Chugani, H. T., Phelps, M. E. & Mazziotta, J. C. Positron emission tomography study of human brain functional development. Ann. Neurol. 22, 487–497 (1987).

98. Scott, L. et al. Human oocyte respiration-rate measurement–potential to improve oocyte and embryo selection? Reprod. Biomed. Online 17, 461–469 (2008).

99. Froehle, A. W. & Churchill, S. E. Energetic competition between Neandertals and anatomically modern humans. PaleoAnthropology 96–116 (2009).

100. Ruff, C. B., Trinkaus, E. & Holliday, T. W. Body mass and encephalization in Pleistocene Homo. Nature 387, 173–176 (1997).

101. Rightmire, G. P. Brain size and encephalization in early to mid-pleistocene Homo. Am. J. Phys. Anthropol. 124, 109–123 (2004).

102. McHenry, H. M. Tempo and mode in human evolution. Proc. Natl. Acad. Sci. USA 91, 6780–6786 (1994).

103. McHenry, H. M. & Coffing, K. Australopithecus to Homo: transformations in body and mind. Annu. Rev. Anthropol. 29, 125–146 (2000).

104. Grabowski, M., Hatala, K. G., Jungers, W. L. & Richmond, B. G. Body mass estimates of hominin fossils and the evolution of human body size. J. Hum. Evol. 85, 75–93 (2015).

105. Kubo, D., Kono, R. T. & Kaifu, Y. Brain size of Homo floresiensis and its evolutionary implications. Proc. R. Soc. B 280, 20130338 (2013).

106. Leigh, S. R. & Shea, B. T. Ontogeny of body size variation in African apes. Am. J. Phys. Anthropol. 99, 43–65 (1996).

107. Wang, R. J., Al-Safar, S. I., Rogers, J. & Hahn, M. W. Human generation times across the past 250,000 years. Sci. Adv. 9, eabm7047 (2023).

