## Supplementary Information for "Evo-devo dynamics of hominin brain size"

### Supplementary Information for:
### Evo-devo dynamics of hominin brain size

Mauricio González-Forero*[1]

[1]School of Biology, University of St Andrews, St Andrews, UK

#### Contents

**Overview** 3

**S1 Brain model** 3
- S1.1 Developmental map, $\mathbf{g}_a$ 3
  - S1.1.1 Energy budget for tissue growth: growth metabolic rate, $B_{\text{syn}}$ 3
  - S1.1.2 Developmental map for tissue mass, $g_{ia}$ for $i \in \{\text{b}, \text{r}, \text{s}\}$ 4
  - S1.1.3 Brain's energy budget for learning: learning metabolic rate, $B_{\text{syn,k}}$ 5
  - S1.1.4 Developmental map for skill level, $g_{\text{k}a}$ 5
  - S1.1.5 Summary 5
- S1.2 Fertility and survival, $f_a$ and $p_a$ 5
  - S1.2.1 Their definition in the model 6
  - S1.2.2 Density dependence in fertility 6

**S2 Evo-devo dynamics** 7
- Layer 7. Evolutionary dynamics 7
- Layer 6. Genetic covariation 8
- Layer 5. Stabilized effects 9
- Layer 4. Total effects 9
- Layer 3. Total immediate effects 10
- Layer 2. Direct effects 10
- Layer 1. Elementary components 10

**S3 Calculation of direct effects** 10
- S3.1 Direct directional selection, $\partial w/\partial \zeta$ 10
- S3.2 Direct developmental bias of the phenotype, $\partial \mathbf{x}^\intercal/\partial \zeta$ 11
  - S3.2.1 From the phenotype, $\partial \mathbf{x}^\intercal/\partial \mathbf{x}$ 11
  - S3.2.2 From the genotype, $\partial \mathbf{x}^\intercal/\partial \mathbf{y}$ 13
- S3.3 Direct social developmental bias from the phenotype, $\partial \mathbf{x}^\intercal/\partial \bar{\mathbf{x}}$ 14

**S4 Parameter values and ancestral genotypic traits** 15
- S4.1 Parameter values 15
- S4.2 Ancestral genotypic traits 15
  - S4.2.1 Naive 16
  - S4.2.2 somewhatNaive 16
  - S4.2.3 ecoSols 17
  - S4.2.4 highlySpecified 17
  - S4.2.5 afarensisFromHighlySpecified 17
  - S4.2.6 afarensisFromSomewhatNaive 17
  - S4.2.7 somewhatNaive2 17
  - S4.2.8 afarensisFromNaive2 17
  - S4.2.9 afarensisFromEcoSols 17

*

**S5 Numerical implementation** 17

**S6 Brain and body size data** 18

**S7 Evolvability** 19

**S8 Total fitness effects of maintenance metabolic costs** 19

**S9 Supplementary figures** 21

**References** 33

### Overview

I model the evolutionary and developmental (evo-devo) dynamics of hominin brain size by applying the evo-devo dynamics framework[1] to the developmentally dynamic brain model[2;3] that was previously[2;3] analysed assuming evolutionary equilibrium. This application yields an evolutionary dynamic analysis of the brain model, and so a model of the evo-devo dynamics of brain size. In this Supplementary Information (SI) I describe the methods in detail, in the following order.

First, I describe the brain model and its equations in discrete age, which provides a specific developmental map, fertility, and survival functions to be used in the evo-devo dynamics framework (section S1). Second, I describe the evo-devo dynamics framework and its equations, which ultimately depend on partial derivatives, or direct effect matrices (section S2). Third, I derive the direct effect matrices of the brain model (section S3). Fourth, I list the parameter values and ancestral genotypic traits used (section S4). Fifth, I describe the numerical implementation of the model (section S5). Sixth, I describe the adult brain and body size data used to contrast the model with (section S6). Seventh, I write the formulas used to compute the evolvability angle and evolvability (section S7). Eighth, I write the formulas to compute the total fitness effects of the metabolic costs of maintenance (section S8). Ninth, I provide supplementary figures (section S9).

I use the following notation from matrix calculus[4] throughout. The Jacobian matrix of a vector $\mathbf{a} \in \mathbb{R}^{n\times 1}$ with respect to a vector $\mathbf{b} \in \mathbb{R}^{m\times 1}$ in its standard or transposed form is, respectively,

$$\frac{\partial \mathbf{a}}{\partial \mathbf{b}^\intercal} = \begin{pmatrix} \frac{\partial a_1}{\partial b_1} & \cdots & \frac{\partial a_1}{\partial b_m} \\ \vdots & \ddots & \vdots \\ \frac{\partial a_n}{\partial b_1} & \cdots & \frac{\partial a_n}{\partial b_m} \end{pmatrix} \in \mathbb{R}^{n\times m} \quad \text{or} \quad \frac{\partial \mathbf{a}^\intercal}{\partial \mathbf{b}} = \begin{pmatrix} \frac{\partial a_1}{\partial b_1} & \cdots & \frac{\partial a_n}{\partial b_1} \\ \vdots & \ddots & \vdots \\ \frac{\partial a_1}{\partial b_m} & \cdots & \frac{\partial a_n}{\partial b_m} \end{pmatrix} \in \mathbb{R}^{m\times n}. \tag{S1}$$

The transpose of $\partial\mathbf{a}/\partial\mathbf{b}^\intercal$ is $(\partial\mathbf{a}/\partial\mathbf{b}^\intercal)^\intercal = \partial\mathbf{a}^\intercal/\partial\mathbf{b}$. The analogous notation applies for total derivatives (i.e., with d rather than $\partial$).

### S1 Brain model

In the terminology of ref.[1], phenotypic traits have developmental dynamics modulated by the genotypic traits. In turn, genotypic traits have evolutionary dynamics modulated by mutation of genotypic traits, development of phenotypic traits, and selection on phenotypic and genotypic traits. The brain model considers four phenotypic traits (state variables; $N_\mathrm{p} = 4$) and three genotypic traits (control variables; $N_\mathrm{g} = 3$). A mutant's phenotypic trait $i$ at age $a$ is $x_{ia} \in \mathbb{R}_{\geq 0}$ for $i \in \{\mathrm{b,r,s,k}\}$ and $a \in \{1,2,\dots,N_\mathrm{a}\}$. The first three phenotypic traits, $x_{ia}$ for $i \in \{\mathrm{b,r,s}\}$, are the mutant's mass of brain, reproductive, and somatic tissue at age $a$, respectively. The fourth phenotypic trait, $x_{\mathrm{k}a}$, is the mutant's skill level at age $a$. In turn, a mutant's genotypic trait $i$ at age $a$ is $y_{ia} \in \mathbb{R}$ for $i \in \{\mathrm{b,r,s}\}$ and $a \in \{1,2,\dots,N_\mathrm{a}\}$, which is the mutant's effort to grow brain, reproductive, and somatic tissues at age $a$, respectively. To compute all the elements of the evolutionary dynamics, one must specify various elementary components (listed in section S2), including the developmental map $\mathbf{g}_a$, fertility $f_a$, and survival probability $p_a$ (for all $a \in \{1,\dots,N_\mathrm{a}\}$). In this section S1, I specify these elementary components of the brain model.

I specify the developmental map $\mathbf{g}_a$ by writing first the developmental maps for tissue mass and then the developmental map for skill level. To specify the developmental maps for tissue mass, I first write the energy budget for tissue production. Similarly, to specify the developmental map for skill level, I first write the brain's energy budget for learning.

#### S1.1 Developmental map, $\mathbf{g}_a$

##### S1.1.1 Energy budget for tissue growth: growth metabolic rate, $B_\mathrm{syn}$

Following West et al.[5], the brain model quantifies an individual's energy budget for tissue growth per unit time with the growth metabolic rate, which is the rate of heat release by the body at rest due to tissue production and equals the resting metabolic rate minus the metabolic rate due to tissue maintenance. Thus, a mutant's growth metabolic rate at age $a$ is

$$B_\mathrm{syn}(\mathbf{x}_a, \bar{x}_{\mathrm{k}a}, a) = Ke(x_{\mathrm{k}a}, \bar{x}_{\mathrm{k}a}, a)x_{\mathrm{B}a}^\beta - \sum_{i\in\{\mathrm{b,r,s}\}} x_{ia}B_i \tag{S2}$$

where the mutant's resting metabolic rate is $Ke(x_{\mathrm{k}a}, \bar{x}_{\mathrm{k}a}, a)x_{\mathrm{B}a}^\beta$ from Kleiber's law, with $K$ and $\beta$ being the Kleiber's law parameters. A mutant's body mass at age $a$ is $x_{\mathrm{B}a} = \sum_{i\in\{\mathrm{b,r,s}\}} x_{ia}$. The mutant's energy extraction efficiency (EEE) is $e(x_{\mathrm{k}a}, \bar{x}_{\mathrm{k}a}, a) \in [0,1]$, which depends on the mutant's skill level and on the skill level of social partners of the same age. EEE decreases exogenously with age $a$ to describe the effect of diminishing maternal care with age. The mutant's maintenance metabolic rate is $\sum_{i\in\{\mathrm{b,r,s}\}} x_{ia}B_i$, where $B_i$ is the mass-specific metabolic cost of maintenance of tissue $i$.

Refs.[2;3] make phenomenological considerations to obtain an expression for EEE as follows. At each age, a mutant faces energy extraction challenges of type $j \in \{1,2,3,4\}$ with probability $P_j$ (and $\sum_{j=1}^4 P_j = 1$), where the challenge types are:

ecological (me vs nature) if $j = 1$,
cooperative (us vs nature) if $j = 2$,
between-individual competitive (me vs you) if $j = 3$, or
between-group competitive (us vs them) if $j = 4$.

A mutant's energy extraction efficiency when facing type-$j$ challenges is $e_j(x_{\mathrm{k}a}, \bar{x}_{\mathrm{k}a}, a)$. Thus, a mutant's EEE is

$$e(x_{\mathrm{k}a}, \bar{x}_{\mathrm{k}a}, a) = \sum_{j=1}^{4} P_j e_j(x_{\mathrm{k}a}, \bar{x}_{\mathrm{k}a}, a).$$

A mutant succeeds at type-$j$ challenges a fraction $S_j(x_{\mathrm{k}a}, \bar{x}_{\mathrm{k}a})$ of the time. The EEE from maternal provisioning in case of individual failure at energy extraction at age $a$ is $\varphi(a)$. Hence, let a mutant's EEE when facing type-$j$ challenges be

$$e_j(x_{\mathrm{k}a}, \bar{x}_{\mathrm{k}a}, a) = S_j(x_{\mathrm{k}a}, \bar{x}_{\mathrm{k}a}) + [1 - S_j(x_{\mathrm{k}a}, \bar{x}_{\mathrm{k}a})]\varphi(a). \tag{S3}$$

Let the success proportion for type-$j$ challenges be

$$S_j(x_{\mathrm{k}a}, \bar{x}_{\mathrm{k}a}) = \frac{c_j(h_j(x_{\mathrm{k}a}, \bar{x}_{\mathrm{k}a}))}{c_j(h_j(x_{\mathrm{k}a}, \bar{x}_{\mathrm{k}a})) + d_j(\bar{x}_{\mathrm{k}a})}, \tag{S4}$$

where $c_j(h_j(x_{\mathrm{k}a}, \bar{x}_{\mathrm{k}a}))$ measures the individual's "competence" at type-$j$ challenges and $d_j(\bar{x}_{\mathrm{k}a})$ measures the type-$j$ challenge "difficulty" at age $a$. Let the competence take the same functional form across challenge types, so let us write $c_j(h_j(x_{\mathrm{k}a}, \bar{x}_{\mathrm{k}a})) = c(h_j(x_{\mathrm{k}a}, \bar{x}_{\mathrm{k}a}))$, where the competence function takes one of two forms:

$$c(h_j(x_{\mathrm{k}a}, \bar{x}_{\mathrm{k}a})) = \begin{cases} [h_j(x_{\mathrm{k}a}, \bar{x}_{\mathrm{k}a})]^{\gamma} & \text{with power competence} \\ \exp(\gamma h_j(x_{\mathrm{k}a}, \bar{x}_{\mathrm{k}a})) & \text{with exponential competence.} \end{cases}$$

The function $h_j(x_{\mathrm{k}a}, \bar{x}_{\mathrm{k}a})$ describes how the skills of cooperating partners interact to yield a "joint" competence $c(h_j(x_{\mathrm{k}a}, \bar{x}_{\mathrm{k}a}))$. Consider the following forms

$$h_j(x_{\mathrm{k}a}, \bar{x}_{\mathrm{k}a}) = \begin{cases} x_{\mathrm{k}a} & \text{if } j \in \{1,3\} \\ x_{\mathrm{k}a} + \bar{x}_{\mathrm{k}a} & \text{if } j \in \{2,4\} \text{ with additive cooperation} \\ x_{\mathrm{k}a}\bar{x}_{\mathrm{k}a} & \text{if } j \in \{2,4\} \text{ with multiplicative cooperation} \\ (x_{\mathrm{k}a}\bar{x}_{\mathrm{k}a})^{1/2} & \text{if } j \in \{2,4\} \text{ with sub-multiplicative cooperation.} \end{cases} \tag{S5}$$

Let environmental difficulty be

$$d_j(\bar{x}_{\mathrm{k}a}) = \begin{cases} \alpha & \text{if } j \in \{1,2\} \\ c(h_j(\bar{x}_{\mathrm{k}a}, \bar{x}_{\mathrm{k}a})) & \text{if } j \in \{3,4\}, \end{cases}$$

where $\alpha$ is a fixed parameter measuring the difficulty of ecological challenges. Finally, I let EEE from maternal provisioning be $\varphi(a) = \varphi_1 \exp(-\varphi_{\mathrm{r}}(a-1))$, with parameters $\varphi_1 \in [0,1]$ and $\varphi_{\mathrm{r}} \geq 0$ (I use this form of $\varphi(a)$ rather than $\varphi_1 \exp(-\varphi_{\mathrm{r}} a)$ as used by ref.[3] because the initial age bin referring to newborns is here 1 rather than 0, which is merely a convention used for reproductive values to take a classic form[6]).

##### S1.1.2 Developmental map for tissue mass, $g_{ia}$ for $i \in \{\mathrm{b,r,s}\}$

I now specify the developmental map for tissue mass, $g_{ia}$, for brain, reproductive, and somatic tissues, so $i \in \{\mathrm{b,r,s}\}$. The fraction of growth metabolic rate allocated to the production of tissue $i$ by a mutant individual of age $a$ is $q_{ia}(\mathbf{y}_a)$ for $i \in \{\mathrm{b,r,s}\}$. I let

$$q_{ia}(\mathbf{y}_a) = \frac{\exp(y_{ia})}{\sum_{i\in\{\mathrm{b,r,s}\}} \exp(y_{ia})}, \tag{S6}$$

which is a function of the effort $y_{ia}$ (relative to a baseline value) to grow tissue $i$ at age $a$ for $i \in \{\mathrm{b,r,s}\}$. Ref.[3] used $q_{ia}(\mathbf{y}_a)$ as control variables, but here I use $y_{ia}$ as control variables because numerical solution is easier as $y_{ia}$ has no constraints and can thus take any real value (the $y$'s do not have to be truncated to remain between zero and one, and their sum at a given age does not have to add to one). From Eq. (S6), it follows that

$$q_{ia}(\mathbf{y}_a) \in [0,1] \text{ for all } i \in \{\mathrm{b,r,s}\} \text{ and } \sum_{i\in\{\mathrm{b,r,s}\}} q_{ia}(\mathbf{y}_a) = 1 \tag{S7}$$

for all $a \in \{1,\dots,N_\mathrm{a}\}$, and similarly for the resident $q_{ia}(\bar{\mathbf{y}}_a)$. Let us now specify the developmental map for tissue mass. The size of tissue $i \in \{\mathrm{b,r,s}\}$ of a mutant individual at age $a+1$ is

$$x_{i,a+1} = g_{ia}(\mathbf{z}_a, \bar{x}_{\mathrm{k}a}, a) = \begin{cases} x_{ia} + \dfrac{q_{ia}(\mathbf{y}_a)}{E_i} B_\mathrm{syn}(\mathbf{x}_a, \bar{x}_{\mathrm{k}a}, a) & \text{if } x_{ia} + \dfrac{q_{ia}(\mathbf{y}_a)}{E_i} B_\mathrm{syn}(\mathbf{x}_a, \bar{x}_{\mathrm{k}a}, a) \geq 0 \\ 0 & \text{otherwise.} \end{cases} \tag{S8}$$

where $E_i$ is the heat released for producing a unit of mass of tissue $i \in \{\mathrm{b,r,s}\}$.

##### S1.1.3 Brain's energy budget for learning: learning metabolic rate, $B_{\mathrm{syn,k}}$

The brain model quantifies the brain's energy budget for learning with the learning metabolic rate, which is the rate of heat release by the brain at rest due to learning and equals the brain metabolic rate allocated to skills minus the brain metabolic rate due to skill maintenance (memory). For a mutant of age $a$, the learning metabolic rate is

$$B_\mathrm{syn,k}(\mathbf{z}_a, \bar{x}_{\mathrm{k}a}, a) = s_\mathrm{k} B_\mathrm{rest,b}(\mathbf{z}_a, \bar{x}_{\mathrm{k}a}, a) - x_{\mathrm{k}a} B_\mathrm{k}, \tag{S9a}$$

where $B_\mathrm{rest,b}(\mathbf{z}_a, \bar{x}_{\mathrm{k}a}, a)$ is the brain metabolic rate, $s_\mathrm{k}$ is the fraction of brain metabolic rate allocated to skills, and $B_\mathrm{k}$ is the metabolic cost of maintaining a unit of skill (metabolic cost of memory). From energy conservation, the brain metabolic rate is

$$B_\mathrm{rest,b}(\mathbf{z}_a, \bar{x}_{\mathrm{k}a}, a) = x_{\mathrm{b}a} B_\mathrm{b} + \left[g_{\mathrm{b}a}(\mathbf{z}_a, \bar{x}_{\mathrm{k}a}, a) - x_{\mathrm{b}a}\right] E_\mathrm{b}. \tag{S9b}$$

Indeed, $x_{\mathrm{b}a} B_\mathrm{b}$ is the rate of heat release by the brain due to maintenance of existing brain tissue, and $\left[g_{\mathrm{b}a}(\mathbf{z}_a, \bar{x}_{\mathrm{k}a}, a) - x_{\mathrm{b}a}\right] E_\mathrm{b}$ is the rate of heat release by the brain due to production of new brain tissue.

##### S1.1.4 Developmental map for skill level, $g_{\mathrm{k}a}$

I now specify the developmental map for skill level. The skill level of a mutant individual at age $a+1$ is

$$x_{\mathrm{k},a+1} = g_{\mathrm{k}a}(\mathbf{z}_a, \bar{x}_{\mathrm{k}a}, a) = \begin{cases} x_{\mathrm{k}a} + \dfrac{1}{E_\mathrm{k}} B_\mathrm{syn,k}(\mathbf{z}_a, \bar{x}_{\mathrm{k}a}, a) & \text{if } x_{\mathrm{k}a} + \dfrac{1}{E_\mathrm{k}} B_\mathrm{syn,k}(\mathbf{z}_a, \bar{x}_{\mathrm{k}a}, a) \geq 0 \\ 0 & \text{otherwise.} \end{cases} \tag{S10}$$

In words, a mutant's skill level at age $a+1$ is the individual's skill level at the previous age plus the brain's rate of heat release due to skill growth $B_\mathrm{syn,k}(\mathbf{z}_a, \bar{x}_{\mathrm{k}a}, a)$ divided by the metabolic cost $E_\mathrm{k}$ of learning a unit of skill.

##### S1.1.5 Summary

The mutant phenotype at age $a+1$ is thus given by the recurrence

$$\mathbf{x}_{a+1} = \mathbf{g}_a(\mathbf{z}_a, \bar{x}_{\mathrm{k}a}, a) = \left(g_{\mathrm{b}a}(\mathbf{z}_a, \bar{x}_{\mathrm{k}a}, a) \quad g_{\mathrm{r}a}(\mathbf{z}_a, \bar{x}_{\mathrm{k}a}, a) \quad g_{\mathrm{s}a}(\mathbf{z}_a, \bar{x}_{\mathrm{k}a}, a) \quad g_{\mathrm{k}a}(\mathbf{z}_a, \bar{x}_{\mathrm{k}a}, a)\right), \tag{S11}$$

with initial condition $\mathbf{x}_1 = \bar{\mathbf{x}}_1$ fixed, where $\mathbf{z}_a = (\mathbf{x}_a; \mathbf{y}_a)$ is the mutant geno-phenotype at age $a$, and with the developmental maps given by Eqs. (S8) and (S10).

In general, social development (i.e., when a mutant's developmental map depends on resident trait values) introduces a complication to evolutionary analysis, such that for example, evaluation of the developmental map at resident genotypic traits may not yield the resident phenotype, particularly if the developmental map at a given age depends on the resident phenotype at other ages (e.g., if individuals learn from older individuals). Ref.[1] handles this complication by computing a socio-devo stable resident phenotype, for which evaluation at resident genotypic traits yields the same phenotype (as explained in section S5). However, this complication is immaterial in the current brain model because the developmental map depends on the resident phenotype of the same age (i.e., individuals only compete or cooperate with peers). So in the current brain model, the resident phenotype is always socio-devo stable (section S5.1).

#### S1.2 Fertility and survival, $f_a$ and $p_a$

Now, I specify a mutant's fertility $f_a$ and survival $p_a$ at age $a \in \{1,\dots,N_\mathrm{a}\}$. First, I state how they are specified in the brain model, namely, as constant survival and density dependent fertility to satisfy the adaptive dynamics assumption of the evo-devo dynamics framework that the resident population dynamics asymptotically converges to a carrying capacity. Density dependence may be allowed to occur either on survival or fertility, but the brain model has so far considered density dependence via fertility, so to enable comparison with the original brain model I keep this approach. Second, I specify this density dependence to obtain a specific expression for fertility.

##### S1.2.1 Their definition in the model

To derive an expression for fertility, we use the same energy conservation reasoning as before. Specifically, the rate of heat release by the reproductive tissue for a mutant of age $a$ is $x_{ra}B_r + (x_{r,a+1} - x_{ra})E_r$, a fraction $s_o$ of which is due to offspring production and maintenance. Denoting by $x_{oa}$ the number of offspring a mutant has at age $a$, the rate of heat release by her reproductive tissue due to offspring maintenance and production is $x_{oa}B_o + (x_{o,a+1} - x_{oa})E_o$ where $B_o$ is the per-offspring metabolic cost of offspring maintenance incurred by the reproductive tissue and $E_o$ is the metabolic cost incurred by the reproductive tissue for producing an offspring. From energy conservation, it follows that $s_o[x_{ra}B_r + (x_{r,a+1} - x_{ra})E_r] = x_{oa}B_o + (x_{o,a+1} - x_{oa})E_o$. Assuming that reproductive tissue is narrowly defined, specifically as being preovulatory ovarian follicles, so offspring maintenance by reproductive tissue is negligible ($B_o \approx 0$), and assuming that maintenance of follicles is much costlier than production ($B_r \gg E_r/(\text{age unit})$), solving in the last expression for $(x_{o,a+1} - x_{oa})$ which is mutant fertility yields

$$x_{o,a+1} - x_{oa} = f_a(x_{ra}, \bar{\mathbf{x}}_{r\bullet}) = \frac{s_o B_r}{E_o(\bar{n}^*(\bar{\mathbf{x}}_{r\bullet}))} x_{ra}. \tag{S12}$$

Eq. (S12) introduces density dependence by letting the metabolic cost of offspring production $E_o(\bar{n}^*(\bar{\mathbf{x}}_{r\bullet}))$ depend on the resident population size

$$\bar{n}^*(\bar{\mathbf{x}}_{r\bullet}) = \sum_{a=1}^{N_a} \bar{n}_a^*(\bar{\mathbf{x}}_{r\bullet}), \tag{S13}$$

where $n_a^*(\bar{\mathbf{x}}_{r\bullet})$ is the density of age-$a$ residents at carrying capacity and $\mathbf{x}_{r\bullet} = (x_{r1}, \ldots, x_{rN_a})^\intercal$ is a mutant's follicle count over life. That the resident population size $\bar{n}^*$ is a direct function of the resident follicle count $\bar{\mathbf{x}}_{r\bullet}$ is shown below in Eq. (S17).

As explained in the main text, I let the survival probability at age $a$ be constant with age and evolutionary time, namely $p_a = p = 1 - m$ for all $a \in \{1, \ldots, N_a - 1\}$, where $m \in (0, 1]$ is a constant mortality rate. Also, I let individuals necessarily die at the last age, so $p_{N_a} = 0$.

##### S1.2.2 Density dependence in fertility

It remains to specify how density dependence affects fertility to obtain a specific expression for fertility. To do this, let the metabolic cost of offspring production $E_o(\bar{n}^*(\bar{\mathbf{x}}_{r\bullet}))$ be proportional to the resident population size $\bar{n}^*(\bar{\mathbf{x}}_{r\bullet})$, that is,

$$E_o(\bar{n}^*(\bar{\mathbf{x}}_{r\bullet})) = k\bar{n}^*(\bar{\mathbf{x}}_{r\bullet}), \tag{S14}$$

for a proportionality factor $k$ (the proportionality factor $k$ will be very small, on the order of $10^{-6}$ MJ/y, for the parameter values used which entail a large population size). Using Eq. (S14) in (S12), a mutant's fertility can then be written as

$$f_a(x_{ra}, \bar{\mathbf{x}}_{r\bullet}) = \frac{\tilde{f}_a(x_{ra})}{\bar{n}^*(\bar{\mathbf{x}}_{r\bullet})}, \tag{S15}$$

where a mutant's density-independent fertility is

$$\tilde{f}_a(x_{ra}) = \frac{s_o B_r}{k} x_{ra} = \tilde{f} x_{ra} \tag{S16}$$

with the summary factor $\tilde{f} = s_o B_r / k$. A resident population-density equilibrium $\bar{\mathbf{n}}^*(\bar{\mathbf{x}}_{r\bullet}) = (\bar{n}_1^*, \ldots, \bar{n}_{N_a}^*)^\intercal$ satisfies the Euler-Lotka equation evaluated at neutrality (Eq. S2.4.4 of ref.[1]) namely

$$\sum_{a=1}^{N_a} \ell_a^\circ f_a^\circ = 1,$$

where $\circ$ indicates evaluation at $\mathbf{y} = \bar{\mathbf{y}}$ and the survivorship to age $a$ of neutral mutants is $\ell_a^\circ = \prod_{i=1}^{a-1} p_i^\circ = p^{a-1}$ (which equals mutant survivorship $\ell_a$ since the survival probability is constant). Using Eq. (S15) in the Euler-Lotka equation yields the carrying capacity

$$\bar{n}^*(\bar{\mathbf{x}}_{r\bullet}) = \sum_{a=1}^{N_a} \ell_a^\circ \tilde{f}_a(\bar{x}_{ra}). \tag{S17}$$

Eqs. (S17) and (S16) show that the carrying capacity $\bar{n}^*(\bar{\mathbf{x}}_{r\bullet})$ directly depends on the resident follicle count across life $\bar{\mathbf{x}}_{r\bullet}$, which evolves over evolutionary time, even though the carrying capacity is constant in ecological time. Using Eqs. (S16) and (S17) in Eq. (S15) and simplifying yields the mutant fertility

$$f_a(x_{ra}, \bar{\mathbf{x}}_{r\bullet}) = \frac{x_{ra}}{\sum_{a=1}^{N_a} \ell_a^\circ \bar{x}_{ra}}. \tag{S18}$$

### S2 Evo-devo dynamics

Having specified the brain model, we can now obtain equations to compute its evo-devo dynamics. To do this, in this section, I provide the equations describing the evo-devo dynamics. Following ref.[1], I arrange in layers the equations describing the evo-devo process, such that the top layer 7 describes the evolutionary dynamics and underlying layers describe underlying processes.

#### Layer 7. Evolutionary dynamics

The top layer of the evo-devo process describes the evolutionary dynamics. In one arrangement, the evolutionary dynamics can be described as evo-devo dynamics (Eqs. 1 and 3 of ref.[1]). Specifically, the evolutionary dynamics of the resident genotypic traits are given by the canonical equation of adaptive dynamics[7;1]

$$\frac{\Delta \bar{\mathbf{y}}}{\Delta \tau} = \iota \mathbf{H}_{\mathbf{y}} \frac{\mathrm{d}w}{\mathrm{d}\mathbf{y}}, \tag{S19a}$$

over evolutionary time $\tau$ (one unit of evolutionary time includes the time from mutation to fixation; derivatives are evaluated at $\mathbf{y} = \bar{\mathbf{y}}$ throughout). Here $\iota$ is a scalar measuring mutational input, $\mathbf{H}_{\mathbf{y}}$ is the mutational covariance matrix, and $\mathrm{d}w/\mathrm{d}\mathbf{y}$ is the total selection gradient of the genotype, all of which will be specified in layers below. In turn, at each evolutionary time, the developmental dynamics of the resident phenotype (tissue mass and skill level) are given by

$$\bar{\mathbf{x}}_{a+1} = \mathbf{g}_a(\bar{\mathbf{z}}_a, \bar{x}_{\mathrm{k}a}, a) \tag{S19b}$$

The developmental map $\mathbf{g}_a$ was specified in Eq. (S11). Because development is social in the brain model (i.e., a mutant's developmental map depends on resident trait values), Eqs. (S19) assume that the resident phenotype is socio-devo stable[1]. As stated above, this assumption is met as the resident is always socio-devo stable in the brain model.

In another arrangement, the evolutionary dynamics of the resident phenotype can be understood as the climbing of a fitness landscape using the following equations. Since there is no niche construction and no exogenous environmental change in the brain model, the environment does not change. Hence, the evo-devo dynamics (S19) imply that the evolutionary dynamics of the resident geno-phenotype are given by

$$\frac{\mathrm{d}\bar{\mathbf{z}}}{\mathrm{d}\tau} = \iota \mathbf{L}_{\mathbf{z}} \frac{\partial w}{\partial \mathbf{z}}, \tag{S20}$$

where $\mathbf{L}_{\mathbf{z}}$ is the mechanistic additive socio-genetic cross-covarianze matrix of the geno-phenotype $\mathbf{z}$ and $\partial w/\partial \mathbf{z}$ is the direct selection gradient of the geno-phenotype (from Layer 7, Eq. 4 of ref.[1]). The matrix $\mathbf{L}_{\mathbf{z}}$ describes socio-genetic covariation and socio-genetic constraints.

In yet another arrangement, as the environment does not change, the evolutionary dynamics of the resident phenotype can be written as

$$\frac{\mathrm{d}\bar{\mathbf{z}}}{\mathrm{d}\tau} = \iota \mathbf{L}_{\mathbf{zy}} \frac{\mathrm{d}w}{\mathrm{d}\mathbf{y}}, \tag{S21}$$

where $\mathbf{L}_{\mathbf{zy}}$ is the mechanistic additive socio-genetic cross-covariance matrix between the geno-phenotype and the genotype (from Layer 7, Eq. 5 of ref.[1]).

Because of age structure and since survival $p_j = p$ is constant, using Eqs. 5-7 of ref.[1], a mutant's fitness is

$$w(\mathbf{x}_{\mathrm{r}\bullet}, \bar{\mathbf{x}}_{\mathrm{r}\bullet}) = \frac{1}{T(\bar{\mathbf{x}}_{\mathrm{r}\bullet})} \sum_{j=1}^{N_{\mathrm{a}}} \left[\phi_j f_j(x_{\mathrm{r}j}, \bar{\mathbf{x}}_{\mathrm{r}\bullet}) + \pi_j(\bar{\mathbf{x}}_{\mathrm{r}\bullet}) p\right], \tag{S22a}$$

where the force of selection[8–10] on fertility at age $j$ is

$$\phi_j = \ell_j^\circ = p^{j-1}, \tag{S22b}$$

on survival at age $j$ is

$$\pi_j(\bar{\mathbf{x}}_{\mathrm{r}\bullet}) = \frac{1}{p} \sum_{k=j+1}^{N_{\mathrm{a}}} \ell_k^\circ f_k(\bar{x}_{\mathrm{r}j}, \bar{\mathbf{x}}_{\mathrm{r}\bullet}), \tag{S22c}$$

and generation time[6] is

$$T(\bar{\mathbf{x}}_{\mathrm{r}\bullet}) = \sum_{j=1}^{N_{\mathrm{a}}} j \ell_j^\circ f_j(\bar{x}_{\mathrm{r}j}, \bar{\mathbf{x}}_{\mathrm{r}\bullet}). \tag{S22d}$$

Substituting the expression for fertility in Eq. (S18) into that of fitness in Eq. (S22) and simplifying, a mutant's fitness reduces to

$$w(\mathbf{x}_{\mathrm{r}\bullet}, \bar{\mathbf{x}}_{\mathrm{r}\bullet}) = \frac{1}{\sum_{a=1}^{N_\mathrm{a}} a p^{a-1} \bar{x}_{\mathrm{r}a}} \sum_{j=1}^{N_\mathrm{a}} \left( p^{j-1} x_{\mathrm{r}j} + \sum_{k=j+1}^{N_\mathrm{a}} p^{k-1} \bar{x}_{\mathrm{r}k} \right). \tag{S23}$$

This fitness function depends directly on the mutant's follicle count, but only indirectly on metabolic costs via the developmental constraint (i.e., after substituting $x_{\mathrm{r}j}$ for the corresponding entry of Eq. (S19b)).

I use Eqs. (S19) to obtain numerical solutions for the resident growth efforts over evolutionary time $\tau$ rather than using Eq. (S20) since the latter with socio-devo stabilisation dynamics effectively requires computing (S19) (section S5). Thus, using the numerical solutions obtained from Eqs. (S19), I compute the components of Eq. (S20) to analyse the evolutionary dynamics of the phenotype as the climbing of a fitness landscape.

#### Layer 6. Genetic covariation

The next layer of the evo-devo process describes genetic covariation. Eqs. (S19a) and (S20) depend on mutational input[7]

$$\iota = \frac{1}{2} \mu(\bar{\mathbf{z}}) \bar{n}^*(\bar{\mathbf{x}}_{\mathrm{r}\bullet}) \tag{S24}$$

where $\mu(\bar{\mathbf{z}}) \in [0,1]$ is the mutation rate as a function of the resident geno-phenotype. The mutation rate does not need to be specified as explained below.

In turn, from Eqs. (S16) and (S17), the carrying capacity is

$$\bar{n}^*(\bar{\mathbf{x}}_{\mathrm{r}\bullet}) = \frac{s_\mathrm{o} B_\mathrm{r}}{k} \sum_{a=1}^{N_\mathrm{a}} p^{a-1} \bar{x}_{\mathrm{r}a}. \tag{S25}$$

To eliminate free parameters and leave only one parameter ($\eta_1$) controlling the speed of the evolutionary dynamics, let the proportionality factor $k$ be

$$k = \frac{1}{2} \mu(\bar{\mathbf{z}}) s_\mathrm{o} B_\mathrm{r} \frac{\eta_0}{\eta_1}, \tag{S26}$$

with positive parameters $\eta_0$ and $\eta_1$. Hence, mutational input becomes

$$\iota = \frac{\eta_1}{\eta_0} \sum_{a=1}^{N_\mathrm{a}} p^{a-1} \bar{x}_{\mathrm{r}a}. \tag{S27}$$

The mutation rate does not need to be specified as $\iota$ now depends on parameters $\eta_0$ and $\eta_1$. I further eliminate consideration of $\eta_0$ as follows.

The evolutionary dynamics of the growth efforts in Eq. (S19a) depend on the mutational covariance matrix

$$\mathbf{H_y} = \mathrm{cov}[\mathbf{y}, \mathbf{y}]. \tag{S28}$$

I let the mutational covariance matrix $\mathbf{H_y}$ be diagonal, with diagonal entries equal to a constant $\eta_0 > 0$. As is standard in adaptive dynamics[7], ref.[1] assumes that mutational variance is marginally small, namely $0 < \mathrm{tr}(\mathbf{H_y}) \ll 1$, which yields $\sum_{i=1}^{N_\mathrm{g}} \sum_{a=1}^{N_\mathrm{a}} \eta_0 = N_\mathrm{g} N_\mathrm{a} \eta_0 \ll 1$. Hence, I assume $\eta_0 \ll 1/(N_\mathrm{g} N_\mathrm{a})$. Yet, the occurrence of $\eta_0$ in $\mathbf{H_y}$ cancels with its occurrence in $\iota$, so $\eta_0$ has no effect on the evolutionary dynamics and only scales mutational variation and population size. The fact that $\mathbf{H_y}$ is diagonal with only positive elements in its main diagonal means that there are no absolute mutational constraints (i.e., $\mathbf{H_y}$ is non-singular).

The evolutionary dynamics of the geno-phenotype in Eq. (S20) depend on the mechanistic socio-genetic cross-covariance matrix of the geno-phenotype, which is

$$\mathbf{L_z} = \frac{\mathrm{s}\mathbf{z}}{\mathrm{s}\mathbf{y}^\intercal} \mathbf{H_y} \frac{\mathrm{d}\mathbf{z}^\intercal}{\mathrm{d}\mathbf{y}} \tag{S29}$$

(Layer 6, Eq. 9 of ref.[1]). Biologically crucially, the matrix $\mathbf{L_z}$ is singular because $\mathrm{d}\mathbf{z}^\intercal/\mathrm{d}\mathbf{y}$ has fewer rows than columns. In turn, the evolutionary dynamics of the geno-phenotype in Eq. (S21) depend on the mechanistic socio-genetic cross-covariance matrix between the geno-phenotype and the genotype, which is

$$\mathbf{L_{zy}} = \frac{\mathrm{s}\mathbf{z}}{\mathrm{s}\mathbf{y}^\intercal} \mathbf{H_y} \tag{S30}$$

(Layer 6, Eq. 13 of ref.[1]).

#### Layer 5. Stabilized effects

The next layer of the evo-devo process describes stabilised effects, which are the total effects of perturbing a variable on another variable, after the effects of the perturbation have propagated through the population via social development and such effects have stabilised. The mechanistic socio-genetic cross-covariance matrix of the geno-phenotype in Eq. (S29) depends on the stabilized effects of the genotype on the geno-phenotype

$$\frac{\mathrm{s}\mathbf{z}}{\mathrm{s}\mathbf{y}^\intercal} = \begin{pmatrix} \frac{\mathrm{s}\mathbf{x}}{\mathrm{s}\mathbf{y}^\intercal} \\ \mathbf{I} \end{pmatrix}.$$

Because the developmental map in Eq. (S11) directly depends on the phenotype of social partners, there is direct social developmental bias from the phenotype ($\partial\mathbf{x}^\intercal/\partial\bar{\mathbf{x}} \neq \mathbf{0}$). Yet, the developmental map does not directly depend on the growth effort of social partners, so there is no direct social developmental bias from the genotype ($\partial\mathbf{x}^\intercal/\partial\bar{\mathbf{y}} = \mathbf{0}$). Then, from Layer 5, Eq. S2a of ref.[1] the stabilised effects of the genotype on the phenotype are

$$\frac{\mathrm{s}\mathbf{x}}{\mathrm{s}\mathbf{y}^\intercal} = \frac{\mathrm{s}\mathbf{x}}{\mathrm{s}\bar{\mathbf{x}}^\intercal}\frac{\mathrm{d}\mathbf{x}}{\mathrm{d}\mathbf{y}^\intercal}, \tag{S31}$$

where the stabilised effects of social partners' phenotypes on the mutant's phenotype are

$$\frac{\mathrm{s}\mathbf{x}}{\mathrm{s}\bar{\mathbf{x}}^\intercal} = \left(\mathbf{I} - \frac{\mathrm{d}\mathbf{x}}{\mathrm{d}\bar{\mathbf{x}}^\intercal}\right)^{-1} \tag{S32}$$

(Layer 5, Eq. S1 of ref.[1]).

#### Layer 4. Total effects

The next layer of the evo-devo process describes total effects, which are the total effects of a perturbing a variable on another variable at any age within the individual, before the effects of the perturbation have propagated through the population.

The evolutionary dynamics of the genotype (Eq. S19a) depends the total selection gradient of the genotype, that is, on the total effects of growth efforts on fitness. Since there is no niche construction, the total selection gradient of the genotype is

$$\frac{\mathrm{d}w}{\mathrm{d}\mathbf{y}} = \frac{\partial w}{\partial \mathbf{y}} + \frac{\mathrm{d}\mathbf{x}^\intercal}{\mathrm{d}\mathbf{y}}\frac{\partial w}{\partial \mathbf{x}} = \frac{\partial w}{\partial \mathbf{y}} + \frac{\partial\mathbf{x}^\intercal}{\partial\mathbf{y}}\frac{\mathrm{d}w}{\mathrm{d}\mathbf{x}} \tag{S33}$$

(from line 1 and 5 of Layer 4, Eq. S21 of ref.[1]; see also Eq. 9 of ref.[1]). Note that the first equality is in terms of the total effects of the genotype on the phenotype ($\mathrm{d}\mathbf{x}^\intercal/\mathrm{d}\mathbf{y}$) and direct phenotypic selection ($\partial w/\partial\mathbf{x}$); instead, the second equality is in terms of the direct effects of the genotype on the phenotype ($\partial\mathbf{x}^\intercal/\partial\mathbf{y}$) and total phenotypic selection ($\mathrm{d}w/\mathrm{d}\mathbf{x}$). Because there is no niche construction, the total selection gradient of the phenotype is

$$\frac{\mathrm{d}w}{\mathrm{d}\mathbf{x}} = \frac{\mathrm{d}\mathbf{x}^\intercal}{\mathrm{d}\mathbf{x}}\frac{\partial w}{\partial \mathbf{x}} \tag{S34}$$

(from Layer 4, Eq. 20 of ref.[1]), which is a mechanistic analogue of the extended selection gradient of ref.[11].

The total selection gradient of the genotype (Eq. S33) depends on the developmental matrix $\mathrm{d}\mathbf{x}^\intercal/\mathrm{d}\mathbf{y}$, which describes how the genotype affects the phenotype. The developmental matrix $\mathrm{d}\mathbf{x}^\intercal/\mathrm{d}\mathbf{y}$ is a mechanistic analogue of Wagner's[12] **B** matrix, whose entries contain Fisher's[13] additive effects of allelic substitution, which are the partial regression coefficients of phenotype on gene content. Since there is no niche construction, the developmental matrix is

$$\frac{\mathrm{d}\mathbf{x}^\intercal}{\mathrm{d}\mathbf{y}} = \frac{\partial\mathbf{x}^\intercal}{\partial\mathbf{y}}\frac{\mathrm{d}\mathbf{x}^\intercal}{\mathrm{d}\mathbf{x}} \tag{S35}$$

(from Layer 4, Eq. S2 of ref.[1]; see also Eq. 10 of ref.[1]). The developmental matrix can also be seen as quantifying the total developmental bias of the phenotype from the genotype. The developmental matrix depends on developmental feedback, which since there is no niche construction is

$$\frac{\mathrm{d}\mathbf{x}^\intercal}{\mathrm{d}\mathbf{x}} = \left(2\mathbf{I} - \frac{\partial\mathbf{x}^\intercal}{\partial\mathbf{x}}\right)^{-1} \tag{S36}$$

(from Layer 4, Eq. S1 of ref.[1]; see also Eq. 11 of ref.[1]) which is a mechanistic analogue of the matrix of total effects of variables on themselves as found in path analysis, where effects are regression coefficients[14]. The developmental feedback of the phenotype can also be seen as quantifying the total developmental bias of the phenotype from the phenotype.

The mechanistic socio-genetic cross-covariance matrix of the geno-phenotype (Eq. S29) also depends on the total developmental bias of the geno-phenotype from the genotype, which is

$$\frac{\mathrm{d}\mathbf{z}^\intercal}{\mathrm{d}\mathbf{y}} = \begin{pmatrix} \frac{\mathrm{d}\mathbf{x}^\intercal}{\mathrm{d}\mathbf{y}} & \mathbf{I} \end{pmatrix}$$

(Layer 4, Eq. S7 of ref. [1]).

The stabilised effects of the phenotype on the phenotype (Eq. S32) depend on the total social developmental bias of the phenotype from the phenotype

$$\frac{\mathrm{d}\mathbf{x}}{\mathrm{d}\bar{\mathbf{x}}^\intercal} = \frac{\mathrm{d}\mathbf{x}}{\mathrm{d}\mathbf{x}^\intercal}\frac{\partial\mathbf{x}}{\partial\bar{\mathbf{x}}^\intercal} \tag{S37}$$

(from Layer 4, Eq. S4 of ref. [1]; see also Eq. 28 of ref. [1]).

##### Layer 3. Total immediate effects

The next layer of the evo-devo process describes total immediate effects, which are the total effects that perturbing a variable has on another variable at the same age, without considering downstream effects over development. As in the brain model there is only developmental constraints but no same-age constraints (e.g., environmental constraints implementing niche construction or exogenous environmental change), total immediate effects reduce to direct effects in the brain model.

##### Layer 2. Direct effects

The next layer of the evo-devo process describes direct effects, which are the effects that perturbing a variable has directly on another variable, without considering constraints. I compute the direct effect matrices in section S3 below. Having the direct effect matrices, all the layers above can be computed.

##### Layer 1. Elementary components

The above layers, in particular Layer 2, can be computed from the following core elementary components. Since survival and the environment are constant, the core elementary components are fertility $f_a$, development $\mathbf{g}_a$, and mutational covariation $\mathbf{H}_\mathbf{y}$, which have already been specified.

#### S3 Calculation of direct effects

In this section, I compute the direct effect matrices which are the building blocks of the evo-devo process (Layer 2). In section S3.1, I compute the direct effects on fitness ($\partial w/\partial\boldsymbol{\zeta}$). In section S3.2, I compute the direct effects on the phenotype ($\partial\mathbf{x}^\intercal/\partial\boldsymbol{\zeta}$), of both the phenotype (where $\boldsymbol{\zeta} = \mathbf{x}$) and genotype (where $\boldsymbol{\zeta} = \mathbf{y}$). These direct effects ($\partial w/\partial\boldsymbol{\zeta}$ and $\partial\mathbf{x}^\intercal/\partial\boldsymbol{\zeta}$) are needed to compute the total selection gradient of the genotype and so the evo-devo dynamics (Eq. S19). In section S3.3, I compute the direct effects of social partners' phenotype on mutant's phenotype ($\partial\mathbf{x}^\intercal/\partial\bar{\mathbf{x}}$). These direct effects ($\partial\mathbf{x}^\intercal/\partial\bar{\mathbf{x}}$) are needed to compute the mechanistic socio-genetic cross-covariance matrix of the geno-phenotype $\mathbf{L}_\mathbf{z}$, and so to analyse socio-genetic constraints.

##### S3.1 Direct directional selection, $\partial w/\partial\zeta$

I now compute the direct effects of the genotype and phenotype on fitness. From Eq. (S23), there is no direct selection on genotypic traits:

$$\frac{\partial w}{\partial\mathbf{y}} = \mathbf{0}, \tag{S38a}$$

and no direct selection on brain, skill, or soma:

$$\frac{\partial w}{\partial\mathbf{x}_{\mathrm{b}\bullet}} = \frac{\partial w}{\partial\mathbf{x}_{\mathrm{k}\bullet}} = \frac{\partial w}{\partial\mathbf{x}_{\mathrm{s}\bullet}} = \mathbf{0}. \tag{S38b}$$

There is only direct selection on follicle count. Differentiating Eq. (S23), the direct selection on follicle count at age $a$ is

$$\frac{\partial w}{\partial x_{\mathrm{r}a}} = \frac{1}{\sum_{j=1}^{N_\mathrm{a}} j\ell_j^\circ\bar{x}_{\mathrm{r}j}}\ell_a^\circ,$$

which is positive provided that individuals can survive to age $a$ (i.e., $\ell^\circ_a > 0$). Hence, direct selection favours ever increasing follicle count. Direct selection on follicle count across ages is thus given by the vector

$$\frac{\partial w}{\partial \mathbf{x}_{\mathrm{r}\bullet}} = \frac{1}{\sum_{j=1}^{N_\mathrm{a}} j\ell^\circ_j \bar{x}_{\mathrm{r}j}}\boldsymbol{\ell}^\circ, \tag{S38c}$$

where $\boldsymbol{\ell}^\circ = (\ell^\circ_1,\ldots,\ell^\circ_{N_\mathrm{a}})^\intercal$.

To write these selection gradients as $\partial w/\partial \mathbf{z}$, note that

$$\frac{\partial w}{\partial \mathbf{z}} = \left(\frac{\partial w}{\partial \mathbf{x}};\frac{\partial w}{\partial \mathbf{y}}\right) = \left(\frac{\partial w}{\partial \mathbf{x}};\mathbf{0}\right)$$

where

$$\frac{\partial w}{\partial \mathbf{x}} = \left(\frac{\partial w}{\partial \mathbf{x}_1};\cdots;\frac{\partial w}{\partial \mathbf{x}_{N_\mathrm{a}}}\right)$$

and

$$\frac{\partial w}{\partial \mathbf{x}_a} = \left(\frac{\partial w}{\partial x_{\mathrm{b}a}};\frac{\partial w}{\partial x_{\mathrm{r}a}};\frac{\partial w}{\partial x_{\mathrm{s}a}};\frac{\partial w}{\partial x_{\mathrm{k}a}}\right) = \left(0;\frac{1}{\sum_{j=1}^{N_\mathrm{a}} j\ell^\circ_j \bar{x}_{\mathrm{r}j}}\ell^\circ_a;0;0\right).$$

#### S3.2 Direct developmental bias of the phenotype, $\partial \mathbf{x}^\intercal/\partial \boldsymbol{\zeta}$

##### S3.2.1 From the phenotype, $\partial \mathbf{x}^\intercal/\partial \mathbf{x}$

I now compute the direct effects of the phenotype on the phenotype. This is given by the matrix quantifying the direct developmental bias of the phenotype from the phenotype, which is

$$\left.\frac{\partial \mathbf{x}^\intercal}{\partial \mathbf{x}}\right|_{\mathbf{y}=\bar{\mathbf{y}}} \equiv \left.\begin{pmatrix} \frac{\partial \mathbf{x}_1^\intercal}{\partial \mathbf{x}_1} & \cdots & \frac{\partial \mathbf{x}_{N_\mathrm{a}}^\intercal}{\partial \mathbf{x}_1} \\ \vdots & \ddots & \vdots \\ \frac{\partial \mathbf{x}_1^\intercal}{\partial \mathbf{x}_{N_\mathrm{a}}} & \cdots & \frac{\partial \mathbf{x}_{N_\mathrm{a}}^\intercal}{\partial \mathbf{x}_{N_\mathrm{a}}} \end{pmatrix}\right|_{\mathbf{y}=\bar{\mathbf{y}}} = \left.\begin{pmatrix} \mathbf{I} & \frac{\partial \mathbf{x}_2^\intercal}{\partial \mathbf{x}_1} & \cdots & \mathbf{0} & \mathbf{0} \\ \mathbf{0} & \mathbf{I} & \cdots & \mathbf{0} & \mathbf{0} \\ \vdots & \vdots & \ddots & \vdots & \vdots \\ \mathbf{0} & \mathbf{0} & \cdots & \mathbf{I} & \frac{\partial \mathbf{x}_{N_\mathrm{a}}^\intercal}{\partial \mathbf{x}_{N_\mathrm{a}-1}} \\ \mathbf{0} & \mathbf{0} & \cdots & \mathbf{0} & \mathbf{I} \end{pmatrix}\right|_{\mathbf{y}=\bar{\mathbf{y}}} \in \mathbb{R}^{N_\mathrm{a}N_\mathrm{p}\times N_\mathrm{a}N_\mathrm{p}} \tag{S39}$$

(Layer 2, Eq. S2a of ref.[1]), where for all $a \in \{1,\ldots,N_\mathrm{a}-1\}$

$$\left.\frac{\partial \mathbf{x}_{a+1}^\intercal}{\partial \mathbf{x}_a}\right|_{\mathbf{y}=\bar{\mathbf{y}}} \equiv \left.\begin{pmatrix} \frac{\partial x_{\mathrm{b},a+1}}{\partial x_{\mathrm{b}a}} & \frac{\partial x_{\mathrm{r},a+1}}{\partial x_{\mathrm{b}a}} & \frac{\partial x_{\mathrm{s},a+1}}{\partial x_{\mathrm{b}a}} & \frac{\partial x_{\mathrm{k},a+1}}{\partial x_{\mathrm{b}a}} \\ \frac{\partial x_{\mathrm{b},a+1}}{\partial x_{\mathrm{r}a}} & \frac{\partial x_{\mathrm{r},a+1}}{\partial x_{\mathrm{r}a}} & \frac{\partial x_{\mathrm{s},a+1}}{\partial x_{\mathrm{r}a}} & \frac{\partial x_{\mathrm{k},a+1}}{\partial x_{\mathrm{r}a}} \\ \frac{\partial x_{\mathrm{b},a+1}}{\partial x_{\mathrm{s}a}} & \frac{\partial x_{\mathrm{r},a+1}}{\partial x_{\mathrm{s}a}} & \frac{\partial x_{\mathrm{s},a+1}}{\partial x_{\mathrm{s}a}} & \frac{\partial x_{\mathrm{k},a+1}}{\partial x_{\mathrm{s}a}} \\ \frac{\partial x_{\mathrm{b},a+1}}{\partial x_{\mathrm{k}a}} & \frac{\partial x_{\mathrm{r},a+1}}{\partial x_{\mathrm{k}a}} & \frac{\partial x_{\mathrm{s},a+1}}{\partial x_{\mathrm{k}a}} & \frac{\partial x_{\mathrm{k},a+1}}{\partial x_{\mathrm{k}a}} \end{pmatrix}\right|_{\mathbf{y}=\bar{\mathbf{y}}} \in \mathbb{R}^{N_\mathrm{p}\times N_\mathrm{p}} \tag{S40}$$

(see also Eqs. 18-19 of ref.[1]). So one only needs to calculate the matrix $\partial \mathbf{x}_{a+1}^\intercal/\partial \mathbf{x}_a$ for $a \in \{1,\ldots,N_\mathrm{a}-1\}$.

From Eq. (S8), the direct developmental bias of the mass of tissue $i \in \{\mathrm{b},\mathrm{r},\mathrm{s}\}$ from itself is

$$\frac{\partial x_{i,a+1}}{\partial x_{ia}} = \begin{cases} 1 + \frac{q_{ia}(\mathbf{y}_a)}{E_i}\frac{\partial B_{\mathrm{syn}}(\mathbf{x}_a,\bar{x}_{\mathrm{k}a},a)}{\partial x_{ia}} & \text{if } x_{ia} + \frac{q_{ia}(\mathbf{y}_a)}{E_i}B_{\mathrm{syn}}(\mathbf{x}_a,\bar{x}_{\mathrm{k}a},a) \geq 0 \\ 0 & \text{otherwise.} \end{cases} \tag{S41a}$$

Also from Eq. (S8), the direct developmental bias of the mass of tissue $j \in \{\mathrm{b},\mathrm{r},\mathrm{s}\}$ from the phenotypic trait $i \in \{\mathrm{b},\mathrm{r},\mathrm{s},\mathrm{k}\}$ with $i \neq j$ is

$$\frac{\partial x_{j,a+1}}{\partial x_{ia}} = \begin{cases} \frac{q_{ja}(\mathbf{y}_a)}{E_j}\frac{\partial B_{\mathrm{syn}}(\mathbf{x}_a,\bar{x}_{\mathrm{k}a},a)}{\partial x_{ia}} & \text{if } x_{ja} + \frac{q_{ja}(\mathbf{y}_a)}{E_j}B_{\mathrm{syn}}(\mathbf{x}_a,\bar{x}_{\mathrm{k}a},a) \geq 0 \\ 0 & \text{otherwise.} \end{cases} \tag{S41b}$$

Similarly, from Eq. (S10), the direct developmental bias of skill level from itself is

$$\frac{\partial x_{\mathrm{k},a+1}}{\partial x_{\mathrm{k}a}} = \begin{cases} 1+\dfrac{1}{E_\mathrm{k}}\dfrac{\partial B_{\mathrm{syn,k}}(\mathbf{z}_a,\bar{x}_{\mathrm{k}a},a)}{\partial x_{\mathrm{k}a}} & \text{if } x_{\mathrm{k}a}+\dfrac{1}{E_\mathrm{k}}B_{\mathrm{syn,k}}(\mathbf{z}_a,\bar{x}_{\mathrm{k}a},a)\geq 0 \\ 0 & \text{otherwise.} \end{cases}$$

Also from Eq. (S10), the direct developmental bias of skill level from the mass of tissue $i \in \{\mathrm{b},\mathrm{r},\mathrm{s}\}$ is

$$\frac{\partial x_{\mathrm{k},a+1}}{\partial x_{ia}} = \begin{cases} \dfrac{1}{E_\mathrm{k}}\dfrac{\partial B_{\mathrm{syn,k}}(\mathbf{z}_a,\bar{x}_{\mathrm{k}a},a)}{\partial x_{ia}} & \text{if } x_{\mathrm{k}a}+\dfrac{1}{E_\mathrm{k}}B_{\mathrm{syn,k}}(\mathbf{z}_a,\bar{x}_{\mathrm{k}a},a)\geq 0 \\ 0 & \text{otherwise.} \end{cases}$$

Thus, if $x_{ja}+\dfrac{q_{ja}(\mathbf{y}_a)}{E_j}B_{\mathrm{syn}}(\mathbf{x}_a,\bar{x}_{\mathrm{k}a},a)\geq 0$ for $j\in\{\mathrm{b},\mathrm{r},\mathrm{s}\}$ and $x_{\mathrm{k}a}+\dfrac{1}{E_\mathrm{k}}B_{\mathrm{syn,k}}(\mathbf{z}_a,\bar{x}_{\mathrm{k}a},a)\geq 0$, using Eq. (S40), the age-specific matrix of the direct developmental bias of the phenotype from the phenotype is

$$\frac{\partial \mathbf{x}_{a+1}^\intercal}{\partial \mathbf{x}_a} = \begin{pmatrix} 1+\dfrac{q_{\mathrm{b}a}}{E_\mathrm{b}}\dfrac{\partial B_{\mathrm{syn}}}{\partial x_{\mathrm{b}a}} & \dfrac{q_{\mathrm{r}a}}{E_\mathrm{r}}\dfrac{\partial B_{\mathrm{syn}}}{\partial x_{\mathrm{b}a}} & \dfrac{q_{\mathrm{s}a}}{E_\mathrm{s}}\dfrac{\partial B_{\mathrm{syn}}}{\partial x_{\mathrm{b}a}} & \dfrac{1}{E_\mathrm{k}}\dfrac{\partial B_{\mathrm{syn,k}}}{\partial x_{\mathrm{b}a}} \\ \dfrac{q_{\mathrm{b}a}}{E_\mathrm{b}}\dfrac{\partial B_{\mathrm{syn}}}{\partial x_{\mathrm{r}a}} & 1+\dfrac{q_{\mathrm{r}a}}{E_\mathrm{r}}\dfrac{\partial B_{\mathrm{syn}}}{\partial x_{\mathrm{r}a}} & \dfrac{q_{\mathrm{s}a}}{E_\mathrm{s}}\dfrac{\partial B_{\mathrm{syn}}}{\partial x_{\mathrm{r}a}} & \dfrac{1}{E_\mathrm{k}}\dfrac{\partial B_{\mathrm{syn,k}}}{\partial x_{\mathrm{r}a}} \\ \dfrac{q_{\mathrm{b}a}}{E_\mathrm{b}}\dfrac{\partial B_{\mathrm{syn}}}{\partial x_{\mathrm{s}a}} & \dfrac{q_{\mathrm{r}a}}{E_\mathrm{r}}\dfrac{\partial B_{\mathrm{syn}}}{\partial x_{\mathrm{s}a}} & 1+\dfrac{q_{\mathrm{s}a}}{E_\mathrm{s}}\dfrac{\partial B_{\mathrm{syn}}}{\partial x_{\mathrm{s}a}} & \dfrac{1}{E_\mathrm{k}}\dfrac{\partial B_{\mathrm{syn,k}}}{\partial x_{\mathrm{s}a}} \\ \dfrac{q_{\mathrm{b}a}}{E_\mathrm{b}}\dfrac{\partial B_{\mathrm{syn}}}{\partial x_{\mathrm{k}a}} & \dfrac{q_{\mathrm{r}a}}{E_\mathrm{r}}\dfrac{\partial B_{\mathrm{syn}}}{\partial x_{\mathrm{k}a}} & \dfrac{q_{\mathrm{s}a}}{E_\mathrm{s}}\dfrac{\partial B_{\mathrm{syn}}}{\partial x_{\mathrm{k}a}} & 1+\dfrac{1}{E_\mathrm{k}}\dfrac{\partial B_{\mathrm{syn,k}}}{\partial x_{\mathrm{k}a}} \end{pmatrix},$$

where the direct effect on the growth metabolic rate of perturbing the phenotype $i$ at age $a$ is

$$\frac{\partial B_{\mathrm{syn}}}{\partial x_{ia}} = \begin{cases} Ke(x_{\mathrm{k}a},\bar{x}_{\mathrm{k}a},a)\beta x_\mathrm{B}^{\beta-1}-B_i & \text{if } i\in\{\mathrm{b},\mathrm{r},\mathrm{s}\} \\ K\dfrac{\partial e}{\partial x_{\mathrm{k}a}}x_\mathrm{B}^{\beta} & \text{if } i=\mathrm{k}, \end{cases}$$

and the direct effect on the learning metabolic rate of perturbing the phenotype $i$ at age $a$ is

$$\frac{\partial B_{\mathrm{syn,k}}}{\partial x_{ia}} = \begin{cases} s_\mathrm{k}\dfrac{\partial B_{\mathrm{rest,b}}}{\partial x_{ia}} & \text{if } i\in\{\mathrm{b},\mathrm{r},\mathrm{s}\} \\ s_\mathrm{k}\dfrac{\partial B_{\mathrm{rest,b}}}{\partial x_{\mathrm{k}a}}-B_\mathrm{k} & \text{if } i=\mathrm{k}. \end{cases}$$

In turn, noting that $g_{\mathrm{b}a}=x_{\mathrm{b},a+1}$, the direct effect on the brain metabolic rate of the phenotype $i$ at age $a$ is

$$\frac{\partial B_{\mathrm{rest,b}}}{\partial x_{ia}} = \begin{cases} B_\mathrm{b}+\left(\dfrac{\partial x_{\mathrm{b},a+1}}{\partial x_{\mathrm{b}a}}-1\right)E_\mathrm{b} & \text{if } i=\mathrm{b} \\ \dfrac{\partial x_{\mathrm{b},a+1}}{\partial x_{ia}}E_\mathrm{b} & \text{if } i\in\{\mathrm{r},\mathrm{s},\mathrm{k}\}, \end{cases}$$

where $\partial x_{\mathrm{b},a+1}/\partial x_{ia}$ for $i\in\{\mathrm{b},\mathrm{r},\mathrm{s},\mathrm{k}\}$ is given by Eq. (S41).

Now, the direct effect on the EEE of the individual's skill level at age $a$ is

$$\frac{\partial e}{\partial x_{\mathrm{k}a}} = \sum_{j=1}^{4} P_j\frac{\partial e_j}{\partial x_{\mathrm{k}a}},$$

where the direct effect on the EEE for challenge type $j$ of the individual's skill level at age $a$ is

$$\frac{\partial e_j}{\partial x_{\mathrm{k}a}} = [1-\varphi(a)]\frac{\partial S_j}{\partial x_{\mathrm{k}a}},$$

and the direct effect on the success proportion for challenges of type $j$ from perturbing the individual's skill level at age $a$ is

$$\frac{\partial S_j}{\partial x_{\mathrm{k}a}} = (1-S_j)\frac{1}{c(h_j(x_{\mathrm{k}a},\bar{x}_{\mathrm{k}a}))+d_j(\bar{x}_{\mathrm{k}a})}\frac{\mathrm{d}c}{\mathrm{d}h_j}\frac{\partial h_j}{\partial x_{\mathrm{k}a}}.$$

In turn, the direct effect on the individual's competence from perturbing her skill level at age $a$ is

$$\frac{\mathrm{d}c}{\mathrm{d}h_j} = \begin{cases} \gamma h_j^{\gamma-1} & \text{with power competence} \\ \gamma \exp(\gamma h_j) & \text{with exponential competence,} \end{cases}$$

and the direct effect on the joint action of skills from perturbing the individual's skill level at age $a$ is

$$\frac{\partial h_j}{\partial x_{\mathrm{k}a}} = \begin{cases} 1 & \text{if } j \in \{1,3\} \\ 1 & \text{if } j \in \{2,4\} \text{ with additive cooperation} \\ \bar{x}_{\mathrm{k}a} & \text{if } j \in \{2,4\} \text{ with multiplicative cooperation} \\ \frac{1}{2}\left(\frac{\bar{x}_{\mathrm{k}a}}{x_{\mathrm{k}a}}\right)^{1/2} & \text{if } j \in \{2,4\} \text{ with sub-multiplicative cooperation.} \end{cases}$$

##### S3.2.2 From the genotype, $\partial\mathbf{x}^\intercal/\partial\mathbf{y}$

I now compute the direct effects of the genotype on the phenotype. This is given by the matrix quantifying the direct developmental bias of the phenotype from the genotype, which is

$$\left.\frac{\partial\mathbf{x}^\intercal}{\partial\mathbf{y}}\right|_{\mathbf{y}=\bar{\mathbf{y}}} \equiv \left.\begin{pmatrix} \frac{\partial\mathbf{x}_1^\intercal}{\partial\mathbf{y}_1} & \cdots & \frac{\partial\mathbf{x}_{N_\mathrm{a}}^\intercal}{\partial\mathbf{y}_1} \\ \vdots & \ddots & \vdots \\ \frac{\partial\mathbf{x}_1^\intercal}{\partial\mathbf{y}_{N_\mathrm{a}}} & \cdots & \frac{\partial\mathbf{x}_{N_\mathrm{a}}^\intercal}{\partial\mathbf{y}_{N_\mathrm{a}}} \end{pmatrix}\right|_{\mathbf{y}=\bar{\mathbf{y}}} = \left.\begin{pmatrix} \mathbf{0} & \frac{\partial\mathbf{x}_2^\intercal}{\partial\mathbf{y}_1} & \cdots & \mathbf{0} & \mathbf{0} \\ \mathbf{0} & \mathbf{0} & \cdots & \mathbf{0} & \mathbf{0} \\ \vdots & \vdots & \ddots & \vdots & \vdots \\ \mathbf{0} & \mathbf{0} & \cdots & \mathbf{0} & \frac{\partial\mathbf{x}_{N_\mathrm{a}}^\intercal}{\partial\mathbf{y}_{N_\mathrm{a}-1}} \\ \mathbf{0} & \mathbf{0} & \cdots & \mathbf{0} & \mathbf{0} \end{pmatrix}\right|_{\mathbf{y}=\bar{\mathbf{y}}} \in \mathbb{R}^{N_\mathrm{a}N_\mathrm{g}\times N_\mathrm{a}N_\mathrm{p}} \tag{S42}$$

(Layer 2, Eq. S2b of ref.[1]), where for all $a \in \{1,\ldots,N_\mathrm{a}-1\}$

$$\left.\frac{\partial\mathbf{x}_{a+1}^\intercal}{\partial\mathbf{y}_a}\right|_{\mathbf{y}=\bar{\mathbf{y}}} \equiv \left.\begin{pmatrix} \frac{\partial x_{\mathrm{b},a+1}}{\partial y_{\mathrm{b}a}} & \frac{\partial x_{\mathrm{r},a+1}}{\partial y_{\mathrm{b}a}} & \frac{\partial x_{\mathrm{s},a+1}}{\partial y_{\mathrm{b}a}} & \frac{\partial x_{\mathrm{k},a+1}}{\partial y_{\mathrm{b}a}} \\ \frac{\partial x_{\mathrm{b},a+1}}{\partial y_{\mathrm{r}a}} & \frac{\partial x_{\mathrm{r},a+1}}{\partial y_{\mathrm{r}a}} & \frac{\partial x_{\mathrm{s},a+1}}{\partial y_{\mathrm{r}a}} & \frac{\partial x_{\mathrm{k},a+1}}{\partial y_{\mathrm{r}a}} \\ \frac{\partial x_{\mathrm{b},a+1}}{\partial y_{\mathrm{s}a}} & \frac{\partial x_{\mathrm{r},a+1}}{\partial y_{\mathrm{s}a}} & \frac{\partial x_{\mathrm{s},a+1}}{\partial y_{\mathrm{s}a}} & \frac{\partial x_{\mathrm{k},a+1}}{\partial y_{\mathrm{s}a}} \end{pmatrix}\right|_{\mathbf{y}=\bar{\mathbf{y}}} \in \mathbb{R}^{N_\mathrm{g}\times N_\mathrm{p}}. \tag{S43}$$

So one only needs to calculate the matrix $\partial\mathbf{x}_{a+1}^\intercal/\partial\mathbf{y}_a$ for $a \in \{1,\ldots,N_\mathrm{a}-1\}$.

From Eq. (S8), the direct developmental bias of mass of tissue $j \in \{\mathrm{b},\mathrm{r},\mathrm{s}\}$ from genotypic trait $i \in \{\mathrm{b},\mathrm{r},\mathrm{s}\}$ is

$$\frac{\partial x_{j,a+1}}{\partial y_{ia}} = \begin{cases} \frac{\partial q_{ja}(\mathbf{y}_a)}{\partial y_{ia}}\frac{1}{E_j}B_{\mathrm{syn}}(\mathbf{x}_a,\bar{x}_{\mathrm{k}a},a) & \text{if } x_{ja} + \frac{q_{ja}(\mathbf{y}_a)}{E_j}B_{\mathrm{syn}}(\mathbf{x}_a,\bar{x}_{\mathrm{k}a},a) \geq 0 \\ 0 & \text{otherwise,} \end{cases} \tag{S44}$$

where the direct effect on the growth allocation of growth efforts is

$$\frac{\partial q_{ja}(\mathbf{y}_a)}{\partial y_{ia}} = \begin{cases} q_{ja}(\mathbf{y}_a)[1-q_{ja}(\mathbf{y}_a)] & \text{if } i = j \\ -q_{ja}(\mathbf{y}_a)q_{ia}(\mathbf{y}_a) & \text{if } i \neq j. \end{cases}$$

Also, from Eq. (S10), the direct developmental bias of skill level from genotypic trait $i \in \{\mathrm{b},\mathrm{r},\mathrm{s}\}$ is

$$\frac{\partial x_{\mathrm{k},a+1}}{\partial y_{ia}} = \begin{cases} \frac{1}{E_\mathrm{k}}\frac{\partial B_{\mathrm{syn,k}}(\mathbf{z}_a,\bar{x}_{\mathrm{k}a},a)}{\partial y_{ia}} & \text{if } x_{\mathrm{k}a} + \frac{1}{E_\mathrm{k}}B_{\mathrm{syn,k}}(\mathbf{z}_a,\bar{x}_{\mathrm{k}a},a) \geq 0 \\ 0 & \text{otherwise,} \end{cases}$$

where the direct effect on the learning metabolic rate of the growth efforts is

$$\frac{\partial B_{\mathrm{syn,k}}(\mathbf{z}_a,\bar{x}_{\mathrm{k}a},a)}{\partial y_{ia}} = s_\mathrm{k}\frac{\partial B_{\mathrm{rest,b}}(\mathbf{z}_a,\bar{x}_{\mathrm{k}a},a)}{\partial y_{ia}},$$

and the direct effect on the brain metabolic rate of the growth efforts is

$$\frac{\partial B_{\mathrm{rest,b}}(\mathbf{z}_a,\bar{x}_{\mathrm{k}a},a)}{\partial y_{ia}} = \frac{\partial g_{\mathrm{b}a}(\mathbf{z}_a,\bar{x}_{\mathrm{k}a},a)}{\partial y_{ia}}E_\mathrm{b} = \frac{\partial x_{\mathrm{b},a+1}}{\partial y_{ia}}E_\mathrm{b}.$$

Thus, the direct developmental bias of skill level from genotypic trait $i \in \{\mathrm{b},\mathrm{r},\mathrm{s}\}$ reduces to

$$\frac{\partial x_{\mathrm{k},a+1}}{\partial y_{ia}} = \begin{cases} \frac{1}{E_\mathrm{k}} s_\mathrm{k} \frac{\partial q_{\mathrm{b}a}(\mathbf{y}_a)}{\partial y_{ia}} B_\mathrm{syn}(\mathbf{x}_a, \bar{x}_{\mathrm{k}a}, a) & \text{if } x_{\mathrm{k}a} + \frac{1}{E_\mathrm{k}} B_{\mathrm{syn,k}}(\mathbf{z}_a, \bar{x}_{\mathrm{k}a}, a) \geq 0 \text{ and } x_{\mathrm{b}a} + \frac{q_{\mathrm{b}a}(\mathbf{y}_a)}{E_\mathrm{b}} B_\mathrm{syn}(\mathbf{x}_a, \bar{x}_{\mathrm{k}a}, a) \geq 0 \\ 0 & \text{otherwise.} \end{cases}$$

Hence, if $x_{ja} + \frac{q_{ja}(\mathbf{y}_a)}{E_j} B_\mathrm{syn}(\mathbf{x}_a, \bar{x}_{\mathrm{k}a}, a) \geq 0$ for $j \in \{\mathrm{b},\mathrm{r},\mathrm{s}\}$ and $x_{\mathrm{k}a} + \frac{1}{E_\mathrm{k}} B_{\mathrm{syn,k}}(\mathbf{z}_a, \bar{x}_{\mathrm{k}a}, a) \geq 0$, using Eq. (S40), the age-specific matrix of the direct developmental bias of the phenotype from the genotype is

$$\frac{\partial \mathbf{x}_{a+1}^\intercal}{\partial \mathbf{y}_a} = \begin{pmatrix} q_{\mathrm{b}a}(1-q_{\mathrm{b}a})\frac{1}{E_\mathrm{b}} & -q_{\mathrm{r}a}q_{\mathrm{b}a}\frac{1}{E_\mathrm{r}} & -q_{\mathrm{s}a}q_{\mathrm{b}a}\frac{1}{E_\mathrm{s}} & q_{\mathrm{b}a}(1-q_{\mathrm{b}a})\frac{s_\mathrm{k}}{E_\mathrm{k}} \\ -q_{\mathrm{b}a}q_{\mathrm{r}a}\frac{1}{E_\mathrm{b}} & q_{\mathrm{r}a}(1-q_{\mathrm{r}a})\frac{1}{E_\mathrm{r}} & -q_{\mathrm{s}a}q_{\mathrm{r}a}\frac{1}{E_\mathrm{s}} & -q_{\mathrm{b}a}q_{\mathrm{r}a}\frac{s_\mathrm{k}}{E_\mathrm{k}} \\ -q_{\mathrm{b}a}q_{\mathrm{s}a}\frac{1}{E_\mathrm{b}} & -q_{\mathrm{r}a}q_{\mathrm{s}a}\frac{1}{E_\mathrm{r}} & q_{\mathrm{s}a}(1-q_{\mathrm{s}a})\frac{1}{E_\mathrm{s}} & -q_{\mathrm{b}a}q_{\mathrm{s}a}\frac{s_\mathrm{k}}{E_\mathrm{k}} \end{pmatrix} B_\mathrm{syn}.$$

##### S3.3 Direct social developmental bias from the phenotype, $\partial \mathbf{x}^\intercal / \partial \bar{\mathbf{x}}$

I now compute the direct effects of the phenotype of social partners on the phenotype. This is given by the matrix quantifying the direct social developmental bias from the phenotype, which is

$$\left.\frac{\partial \mathbf{x}^\intercal}{\partial \bar{\mathbf{x}}}\right|_{\mathbf{y}=\bar{\mathbf{y}}} = \left.\begin{pmatrix} \frac{\partial \mathbf{x}_1^\intercal}{\partial \bar{\mathbf{x}}_1} & \cdots & \frac{\partial \mathbf{x}_{N_\mathrm{a}}^\intercal}{\partial \bar{\mathbf{x}}_1} \\ \vdots & \ddots & \vdots \\ \frac{\partial \mathbf{x}_1^\intercal}{\partial \bar{\mathbf{x}}_{N_\mathrm{a}}} & \cdots & \frac{\partial \mathbf{x}_{N_\mathrm{a}}^\intercal}{\partial \bar{\mathbf{x}}_{N_\mathrm{a}}} \end{pmatrix}\right|_{\mathbf{y}=\bar{\mathbf{y}}} = \left.\begin{pmatrix} \mathbf{0} & \frac{\partial \mathbf{x}_2^\intercal}{\partial \bar{\mathbf{x}}_1} & \cdots & \mathbf{0} & \mathbf{0} \\ \mathbf{0} & \mathbf{0} & \cdots & \mathbf{0} & \mathbf{0} \\ \vdots & \vdots & \ddots & \vdots & \vdots \\ \mathbf{0} & \mathbf{0} & \cdots & \mathbf{0} & \frac{\partial \mathbf{x}_{N_\mathrm{a}}^\intercal}{\partial \bar{\mathbf{x}}_{N_\mathrm{a}-1}} \\ \mathbf{0} & \mathbf{0} & \cdots & \mathbf{0} & \mathbf{0} \end{pmatrix}\right|_{\mathbf{y}=\bar{\mathbf{y}}} \in \mathbb{R}^{N_\mathrm{a}N_\mathrm{p} \times N_\mathrm{a}N_\mathrm{p}}$$

(Layer 2, Eq. S4 of ref.[1]). From Eqs. (S8) and (S10), the only direct dependence of a mutant's phenotype on social partners' phenotype is via the skill level of social partners of her same age, so for all $a \in \{1, \ldots, N_\mathrm{a}-1\}$

$$\left.\frac{\partial \mathbf{x}_{a+1}^\intercal}{\partial \bar{\mathbf{x}}_a}\right|_{\mathbf{y}=\bar{\mathbf{y}}} \equiv \left.\begin{pmatrix} 0 & 0 & 0 & 0 \\ 0 & 0 & 0 & 0 \\ 0 & 0 & 0 & 0 \\ \frac{\partial x_{\mathrm{b},a+1}}{\partial \bar{x}_{\mathrm{k}a}} & \frac{\partial x_{\mathrm{r},a+1}}{\partial \bar{x}_{\mathrm{k}a}} & \frac{\partial x_{\mathrm{s},a+1}}{\partial \bar{x}_{\mathrm{k}a}} & \frac{\partial x_{\mathrm{k},a+1}}{\partial \bar{x}_{\mathrm{k}a}} \end{pmatrix}\right|_{\mathbf{y}=\bar{\mathbf{y}}} \in \mathbb{R}^{N_\mathrm{p} \times N_\mathrm{p}}.$$

Hence, one only needs to consider the direct social developmental bias from social partners' skill level.

Consequently, using Eq. (S8), the direct social developmental bias of the mass of tissue $j \in \{\mathrm{b},\mathrm{r},\mathrm{s}\}$ from social partners' skill level is

$$\frac{\partial x_{j,a+1}}{\partial \bar{x}_{\mathrm{k}a}} = \begin{cases} \frac{q_{ja}(\mathbf{y}_a)}{E_j} \frac{\partial B_\mathrm{syn}(\mathbf{x}_a, \bar{x}_{\mathrm{k}a}, a)}{\partial \bar{x}_{\mathrm{k}a}} & \text{if } x_{ja} + \frac{q_{ja}(\mathbf{y}_a)}{E_j} B_\mathrm{syn}(\mathbf{x}_a, \bar{x}_{\mathrm{k}a}, a) \geq 0 \\ 0 & \text{otherwise,} \end{cases}$$

where the direct effect on the growth metabolic rate of perturbing a social partner's skill level at age $a$ is

$$\frac{\partial B_\mathrm{syn}(\mathbf{x}_a, \bar{x}_{\mathrm{k}a}, a)}{\partial \bar{x}_{\mathrm{k}a}} = K \frac{\partial e(x_{\mathrm{k}a}, \bar{x}_{\mathrm{k}a}, a)}{\partial \bar{x}_{\mathrm{k}a}} x_\mathrm{B}^\beta,$$

and the direct effect on the EEE of perturbing a social partner's skill level at age $a$ is

$$\frac{\partial e(x_{\mathrm{k}a}, \bar{x}_{\mathrm{k}a}, a)}{\partial \bar{x}_{\mathrm{k}a}} = \sum_{j=1}^{4} P_j \frac{\partial e_j(x_{\mathrm{k}a}, \bar{x}_{\mathrm{k}a}, a)}{\partial \bar{x}_{\mathrm{k}a}}.$$

The direct effect on the EEE for a challenge of type $j$ from perturbing a social partner's skill level at age $a$ is

$$\frac{\partial e_j}{\partial \bar{x}_{\mathrm{k}a}} = [1-\varphi(a)] \frac{\partial S_j}{\partial \bar{x}_{\mathrm{k}a}},$$

and the direct effect on the success proportion from perturbing a social partner's skill level at age $a$ is

$$\frac{\partial S_j}{\partial \bar{x}_{\mathrm{k}a}} = \frac{1}{c(h_j(x_{\mathrm{k}a}, \bar{x}_{\mathrm{k}a})) + d_j(\bar{x}_{\mathrm{k}a})} \left[ (1 - S_j) \frac{\partial c}{\partial h_j} \frac{\partial h_j}{\partial \bar{x}_{\mathrm{k}a}} - S_j \frac{\mathrm{d} d_j}{\mathrm{d} \bar{x}_{\mathrm{k}a}} \right].$$

In turn, the direct effect of the social partner's skill level on the joint action of the skills is

$$\frac{\partial h_j}{\partial \bar{x}_{\mathrm{k}a}} = \begin{cases} 0 & \text{if } j \in \{1,3\} \\ 1 & \text{if } j \in \{2,4\} \text{ with additive cooperation} \\ x_{\mathrm{k}a} & \text{if } j \in \{2,4\} \text{ with multiplicative cooperation} \\ \frac{1}{2}\left(\frac{x_{\mathrm{k}a}}{\bar{x}_{\mathrm{k}a}}\right)^{1/2} & \text{if } j \in \{2,4\} \text{ with sub-multiplicative cooperation,} \end{cases}$$

and the direct (and total) effect of the social partner's skill level on the difficulty of challenges of type $j$ is

$$\frac{\mathrm{d} d_j}{\mathrm{d} \bar{x}_{\mathrm{k}a}} = \begin{cases} 0 & \text{if } j \in \{1,2\} \\ \frac{\mathrm{d} c}{\mathrm{d} h_j} \left( \frac{\partial h_j}{\partial x_{\mathrm{k}a}} + \frac{\partial h_j}{\partial \bar{x}_{\mathrm{k}a}} \right) & \text{if } j \in \{3,4\}. \end{cases}$$

Similarly, using Eq. (S10), the direct social developmental bias of skill level from social partners' skill level is

$$\frac{\partial x_{\mathrm{k},a+1}}{\partial \bar{x}_{\mathrm{k}a}} = \begin{cases} \frac{1}{E_\mathrm{k}} \frac{\partial B_{\mathrm{syn,k}}(\mathbf{z}_a, \bar{x}_{\mathrm{k}a}, a)}{\partial \bar{x}_{\mathrm{k}a}} & \text{if } x_{\mathrm{k}a} + \frac{1}{E_\mathrm{k}} B_{\mathrm{syn,k}}(\mathbf{z}_a, \bar{x}_{\mathrm{k}a}, a) \geq 0 \\ 0 & \text{otherwise,} \end{cases}$$

where the direct effect on the learning metabolic rate from the perturbation of a social partner's skill level at age $a$ is

$$\frac{\partial B_{\mathrm{syn,k}}(\mathbf{z}_a, \bar{x}_{\mathrm{k}a}, a)}{\partial \bar{x}_{\mathrm{k}a}} = s_\mathrm{k} \frac{\partial B_{\mathrm{rest,b}}(\mathbf{z}_a, \bar{x}_{\mathrm{k}a}, a)}{\partial \bar{x}_{\mathrm{k}a}},$$

and the direct effect on the brain metabolic rate from the perturbation of a social partner's skill level at age $a$ is

$$\frac{\partial B_{\mathrm{rest,b}}(\mathbf{z}_a, \bar{x}_{\mathrm{k}a}, a)}{\partial \bar{x}_{\mathrm{k}a}} = \frac{\partial g_{\mathrm{b}a}(\mathbf{z}_a, \bar{x}_{\mathrm{k}a}, a)}{\partial \bar{x}_{\mathrm{k}a}} E_\mathrm{b} = \frac{\partial x_{\mathrm{b},a+1}}{\partial \bar{x}_{\mathrm{k}a}} E_\mathrm{b}.$$

Hence, if $x_{ja} + \frac{q_{ja}(\mathbf{y}_a)}{E_j} B_{\mathrm{syn}}(\mathbf{x}_a, \bar{x}_{\mathrm{k}a}, a) \geq 0$ for $j \in \{\mathrm{b,r,s}\}$ and $x_{\mathrm{k}a} + \frac{1}{E_\mathrm{k}} B_{\mathrm{syn,k}}(\mathbf{z}_a, \bar{x}_{\mathrm{k}a}, a) \geq 0$, using Eq. (S45a), the direct social developmental bias matrix of the phenotype from the phenotype is

$$\frac{\partial \mathbf{x}_{a+1}^\intercal}{\partial \bar{\mathbf{x}}_a} = \begin{pmatrix} 0 & 0 & 0 & 0 \\ 0 & 0 & 0 & 0 \\ 0 & 0 & 0 & 0 \\ \frac{q_{\mathrm{b}a}}{E_\mathrm{b}} & \frac{q_{\mathrm{r}a}}{E_\mathrm{r}} & \frac{q_{\mathrm{s}a}}{E_\mathrm{s}} & s_\mathrm{k}\frac{q_{\mathrm{b}a}}{E_\mathrm{k}} \end{pmatrix} \frac{\partial B_{\mathrm{syn}}}{\partial \bar{x}_{\mathrm{k}a}}.$$

### S4 Parameter values and ancestral genotypic traits

In this section, I list the parameter values and ancestral genotypic traits I used.

#### S4.1 Parameter values

I use the same parameter values as those in ref.[3] (Table S1) plus three additional parameter values that emerge from age discretisation and consideration of the evolutionary dynamics and socio-devo dynamics (Table S2). In contrast to the dynamic optimisation approach of ref.[3], in the present evo-devo dynamics approach the units of the phenotypic traits need not be rescaled to allow for convergence to an evolutionary equilibrium. However, as age is discrete, the unit of developmental time needs to be rescaled so that age bins are in the desired units (Table S2).

#### S4.2 Ancestral genotypic traits

I find that the ancestral conditions of the resident genotypic traits $\bar{y}_{ia}$ at the initial evolutionary time must be carefully provided for the system to evolve out of them. I considered the following nine different ancestral conditions.

| Initial age mass | | Tissue metabolism | | | | Demography | |
|---|---|---|---|---|---|---|---|
| | | $K$ | $132.7281\frac{\mathrm{MJ}}{\mathrm{y}}\mathrm{kg}^{-\beta}$ | $\beta$ | 0.7378 | | |
| $x_{\mathrm{b}1}$ | 0.3372 kg | $B_{\mathrm{b}}$ | $313.0962\frac{\mathrm{MJ}}{\mathrm{y}\times\mathrm{kg}}$ | $E_{\mathrm{b}}$ | $123.7584\frac{\mathrm{MJ}}{\mathrm{kg}}$ | $m$ | $0.034\frac{1}{\mathrm{y}}$ |
| $x_{\mathrm{r}1}$ | 0 kg | $B_{\mathrm{r}}$ | $2697.1179\frac{\mathrm{MJ}}{\mathrm{y}\times\mathrm{kg}}$ | $E_{\mathrm{r}}$ | $190.8196\frac{\mathrm{MJ}}{\mathrm{kg}}$ | $\tilde{f}$ | $1\frac{\#\mathrm{offspring}}{\mathrm{kg}\times\mathrm{y}}$ |
| $x_{\mathrm{s}1}$ | 2.0628 kg | $B_{\mathrm{s}}$ | $29.6891\frac{\mathrm{MJ}}{\mathrm{y}\times\mathrm{kg}}$ | $E_{\mathrm{s}}$ | $12.4594\frac{\mathrm{MJ}}{\mathrm{kg}}$ | $N_{\mathrm{a}}$ | 47 y |

Regime 1: power competence

| Skill metabolism | | Energy extraction | | Maternal provisioning | |
|---|---|---|---|---|---|
| $s_{\mathrm{k}}$ | 0.5 | $\alpha$ | $1\ \mathrm{TB}^{\gamma}$† | $\varphi_1$ | 0.4 |
| $B_{\mathrm{k}}$ | $36\frac{\mathrm{MJ}}{\mathrm{y}\times\mathrm{TB}}$ | $\gamma$ | 1.4 | $\varphi_{\mathrm{r}}$ | $0.2\frac{1}{\mathrm{y}}$ |
| $E_{\mathrm{k}}$ | $370\frac{\mathrm{MJ}}{\mathrm{TB}}$ | $x_{\mathrm{k}1}$ | 1 TB | | |

Regime 2: exponential competence

| Skill metabolism | | Energy extraction | | Maternal provisioning | |
|---|---|---|---|---|---|
| $s_{\mathrm{k}}$ | 0.5 | $\alpha$ | 1.15 | $\varphi_1$ | 0.5 |
| $B_{\mathrm{k}}$ | $50\frac{\mathrm{MJ}}{\mathrm{y}\times\mathrm{TB}}$ | $\gamma$ | $0.6\ \mathrm{TB}^{-1}$‡ | $\varphi_{\mathrm{r}}$ | $0.2\frac{1}{\mathrm{y}}$ |
| $E_{\mathrm{k}}$ | $250\frac{\mathrm{MJ}}{\mathrm{TB}}$ | $x_{\mathrm{k}1}$ | 0 TB | | |

Table S1: Parameter values of ref.[3], which I use here. †For the expression of $e_2$, the unit is $\mathrm{TB}^{2\gamma}$ with multiplicative cooperation but $\mathrm{TB}^{\gamma}$ with the other forms of cooperation (additive or submultiplicative). ‡For the expressions of $e_2$ and $e_4$, the unit is $\mathrm{TB}^{-2}$ with multiplicative cooperation but $\mathrm{TB}^{-1}$ with the other forms of cooperation. For these parameter values, power competence yields a strongly decelerating $S_1$ with increasing skill level (strongly diminishing returns of learning) whereas exponential competence yields a weakly decelerating $S_1$ (weakly diminishing returns of learning). Except when noted otherwise, all figures use exponential competence, submultiplicative cooperation, and the challenge proportions $P_1 = 0.6$, $P_2 = 0.3$, $P_3 = 0$, and $P_4 = 0.1$; this is the *sapiens* scenario found by ref.[3] to yield the evolution of adult brain and body sizes that best fitted those of *Homo sapiens*. The *afarensis* scenario of Fig. 1 uses power competence and submultiplicative cooperation with the challenge proportions indicated in the figure. The *habilis*, *ergaster*, and *erectus* scenarios of Fig. 1 use power competence and additive cooperation with the challenge proportions indicated in the figure. The *heidelbergensis*, *neanderthalensis*, and *sapiens* scenarios of Fig. 1 use exponential competence and submultiplicative cooperation with the challenge proportions indicated in the figure.

| Age unit | | Evolutionary speed | | Socio-devo error tolerance | |
|---|---|---|---|---|---|
| $a_{\mathrm{unit}}$ | 0.1 y | $\eta_1$ | $20000\frac{1}{\mathrm{y}\times\mathrm{kg}}$ | $\mathrm{SDS}_{\mathrm{error}}$ | $10^{-6}$ |

Table S2: Parameter values not present in ref.[3]. I set the age bin size to $a_{\mathrm{unit}} = 0.1$ y (a coarser bin size of 1 y yields much larger numerical error compared to the model's behaviour in continuous age[3]; time units in parameter values are then rescaled to the unit of the age bin size (e.g., $B_{\mathrm{s}} = 29.6891\frac{\mathrm{MJ}}{\mathrm{y}\times\mathrm{kg}}a_{\mathrm{unit}}$). From Eq. (S26), it follows that the units of $\eta_0/\eta_1$ are those of $k/B_{\mathrm{r}}$ and from Eq. (S14), it follows that the units of $k$ are MJ. Hence, the units of $\eta_0/\eta_1$ are y × kg and since $\eta_0$ is non-dimensional because $\mathbf{H}_{\mathbf{y}}$ is non-dimensional, the indicates units of $\eta_1$ follow.

##### S4.2.1 Naive

First, Naive ancestral conditions, where $\bar{y}_{ia} = 0$ for all $i \in \{\mathrm{b}, \mathrm{r}, \mathrm{s}\}$ and $a \in \{1, \ldots, N_{\mathrm{a}}\}$.

##### S4.2.2 somewhatNaive

Second, somewhatNaive ancestral conditions given in Table S3. These conditions are the blue dots in Fig. S4a-c and in Fig. S5a-c.

| $a$ | 1 | 2 | 3 | 4 | 5 | 6 | 7 | 8 | 9 | $10 \ldots N_{\mathrm{a}}$ |
|---|---|---|---|---|---|---|---|---|---|---|
| $\bar{y}_{\mathrm{b}a}$ | −0.1 | −0.1 | −0.1 | −0.1 | 0 | 0 | 0 | 0 | 0 | 0 |
| $\bar{y}_{\mathrm{r}a}$ | −1.2 | −1.1 | −1.0 | −0.9 | −0.7 | −0.5 | −0.4 | −0.2 | −0.1 | 0 |
| $\bar{y}_{\mathrm{s}a}$ | 4.3 | 4.2 | 4.1 | 3.9 | 3.8 | 3.6 | 3.4 | 3.2 | 3.1 | 3 |

Table S3: somewhatNaive ancestral conditions for resident genotypic traits.

##### S4.2.3 ecoSols

Third, ecoSols ancestral conditions given by the solutions of the evo-devo dynamics with $P_1 = 1$ starting from the somewhatNaive ancestral conditions in Table S3. These conditions are the red dots in Fig. S5a-c and the blue dots in Fig. S6a-c.

##### S4.2.4 highlySpecified

Fourth, highlySpecified ancestral conditions given by

$$\bar{y}_{\mathrm{b}a} = -6\exp(-0.2(a-1)a_{\mathrm{unit}}) \times \sin((a-1)a_{\mathrm{unit}} + 0.15) \tag{S45a}$$
$$\bar{y}_{\mathrm{r}a} = -6\exp(-0.2(a-1)a_{\mathrm{unit}}) \times [0.1\sin((a-1)a_{\mathrm{unit}}) + 0.3] \tag{S45b}$$
$$\bar{y}_{\mathrm{s}a} = 3\exp(-0.2(a-1)a_{\mathrm{unit}}) \times [0.5\sin((a-1)a_{\mathrm{unit}}) + 1] + 3. \tag{S45c}$$

Eqs. (S45) were manually obtained by seeking to roughly replicate with dampening sinusoidal curves the ecoSols ancestral conditions (red dots in Fig. S5a-c). These highlySpecified conditions yield the development of adult brain and body size of australopithecine scale and the evolution of adult brain and body sizes of modern human scale (not shown).

##### S4.2.5 afarensisFromHighlySpecified

Fifth, afarensisFromHighly specified ancestral conditions given by the solutions of the *afarensis* scenario started from the highlySpecified ancestral conditions.

##### S4.2.6 afarensisFromSomewhatNaive

Sixth, afarensisFromSomewhatNaive ancestral conditions given by the solutions of the *afarensis* scenario started from the somewhatNaive ancestral conditions.

##### S4.2.7 somewhatNaive2

Seventh, somewhatNaive2 ancestral conditions given by

$$\bar{y}_{\mathrm{b}a} = k_{1\mathrm{b}}\exp[-k_{2\mathrm{b}}(a-k_{3\mathrm{b}})] \tag{S46a}$$
$$\bar{y}_{\mathrm{r}a} = k_{1\mathrm{r}}\frac{1}{1+\exp[-k_{2\mathrm{r}}(a-k_{3\mathrm{r}})]} \tag{S46b}$$
$$\bar{y}_{\mathrm{s}a} = k_{1\mathrm{s}}\frac{1}{1+\exp[-k_{2\mathrm{s}}(a-k_{3\mathrm{s}})]}, \tag{S46c}$$

with

$$\begin{pmatrix} k_{1\mathrm{b}} & k_{1\mathrm{r}} & k_{1\mathrm{s}} \\ k_{2\mathrm{b}} & k_{2\mathrm{r}} & k_{2\mathrm{s}} \\ k_{3\mathrm{b}} & k_{3\mathrm{r}} & k_{3\mathrm{s}} \end{pmatrix} = \begin{pmatrix} 1 & 2 & 10 \\ 0.2\frac{1}{\mathrm{y}} & 10\frac{1}{\mathrm{y}} & 0\frac{1}{\mathrm{y}} \\ 7\,\mathrm{y} & 10\,\mathrm{y} & 10\,\mathrm{y} \end{pmatrix}. \tag{S46d}$$

These conditions are the blue dots in Fig. S1a-c.

##### S4.2.8 afarensisFromNaive2

Eighth, afarensisFromNaive2 ancestral conditions given by the solutions of the *afarensis* scenario started from the somewhatNaive2 ancestral conditions. These conditions are the red dots in Fig. S1a-c and the blue dots in Fig. S2a-c.

##### S4.2.9 afarensisFromEcoSols

Ninth, afarensisFromEcoSols ancestral conditions given by the solutions of the *afarensis* scenario started from the ecoSols ancestral conditions.

#### S5 Numerical implementation

I obtained numerical solutions as follows. Each evolutionary time $\tau$ has two phases: socio-devo dynamics and evolutionary dynamics. The phase of socio-devo dynamics finds a resident socio-devo stable geno-phenotype with the current resident genotype, whereas the phase of evolutionary dynamics determines the resident genotype for the next evolutionary time. For the brain model, the resident is always socio-devo stable so the socio-devo dynamics phase is unnecessary. I double check this numerically by implementing socio-devo stabilisation dynamics regardless. So I obtain socio-devo stable resident phenotypes via socio-devo stabilisation dynamics[1] at each $\tau$ before computing the change in resident genotype with Eqs. (S19).

##### S5.1 Socio-devo dynamics phase

I compute the socio-devo dynamics as follows. Recall that the resident phenotype across life is $\bar{\mathbf{x}} = (\bar{\mathbf{x}}_1;\ldots;\bar{\mathbf{x}}_{N_\mathrm{a}})$. During the socio-dynamics phase, I denote by $\bar{\mathbf{x}}(\theta)$ the resident phenotype at socio-devo time $\theta$.

First, I find the initial condition for the socio-devo dynamics by solving the recurrence

$$\bar{\mathbf{x}}_{a+1} = \mathbf{g}_a(\bar{\mathbf{x}}_a, \bar{\mathbf{y}}_a, \bar{x}_{\mathrm{k}a}, a) \tag{S47a}$$

for all $a \in \{1,\ldots,N_\mathrm{a}-1\}$ with initial condition $\bar{\mathbf{x}}_1$. I use this solution as the initial socio-devo resident phenotype, so $\bar{\mathbf{x}} = \bar{\mathbf{x}}(1)$.

Then, I solve the socio-devo dynamics by solving the recurrence

$$\bar{\mathbf{x}}_{a+1}(\theta+1) = \mathbf{g}_a(\bar{\mathbf{x}}_a(\theta+1), \bar{\mathbf{y}}_a, \bar{x}_{\mathrm{k}a}(\theta), a) \tag{S47b}$$

for all $a \in \{1,\ldots,N_\mathrm{a}-1\}$ and all $\theta \in \{1,2,\ldots,\theta_\mathrm{end}\}$ with initial condition $\bar{\mathbf{x}}_1$ and $\bar{\mathbf{x}}(1)$.

I consider the socio-devo dynamics to have converged to a socio-devo stable equilibrium $\bar{\mathbf{x}}^{**} = \bar{\mathbf{x}}(\theta+1)$ at $\theta+1$ if the maximum entry of $|\bar{\mathbf{x}}(\theta+1) - \bar{\mathbf{x}}(\theta)|$ is smaller than $\mathrm{SDS}_\mathrm{error}$. I use $\bar{\mathbf{x}}^{**}$ as the (socio-devo stable) resident in the evolutionary dynamics phase, dropping $^{**}$ for simplicity.

It can be checked that the initial conditions (S47a) for the socio-devo dynamics immediately yield a socio-devo stable resident for the brain model. This is possible because the developmental map at a given age depends on the resident phenotype at the same age. This is not generally the case if the developmental map depends on the resident phenotype at other ages in which case the initial conditions for the socio-devo dynamics are not so straightforward[1] (for instance, if development depends on immediately older residents, then $\bar{\mathbf{x}}_{a+1} = \mathbf{g}_a(\bar{\mathbf{x}}_a, \bar{\mathbf{y}}_a, \bar{x}_{\mathrm{k},a+1}, a)$ is not a recurrence for $\bar{\mathbf{x}}_{a+1}$ over $a$). It is in such cases that the socio-devo dynamics phase is useful to find a socio-devo stable resident.

##### S5.2 Evolutionary dynamics phase

During the evolutionary dynamics phase at evolutionary time $\tau$, I denote by $\bar{\mathbf{y}}(\tau)$ the resident genotypic traits and by $\bar{\mathbf{x}}(\tau)$ the socio-devo stable resident phenotype. I find the resident genotype at the next evolutionary time step via

$$\bar{\mathbf{y}}(\tau+1) = \bar{\mathbf{y}}(\tau) + \iota \mathbf{H}_\mathbf{y} \frac{\mathrm{d}w}{\mathrm{d}\mathbf{y}}. \tag{S48}$$

A typical run took from 2 to 4 minutes to complete, in contrast to the 2 to 4 days it took with the previous optimal control approach[3]. Runs were implemented in Julia 1.7.2.

##### S5.3 Brain-body allometry from random growth efforts

To generate the dots in Fig. 2a, I sampled $\bar{y}_{ia}$ from the normal distribution with mean 0 and standard deviation 4 for all $i \in \{\mathrm{b},\mathrm{r},\mathrm{s}\}$ and all $a \in \{1,\ldots,N_\mathrm{a}\}$. Then, I used the sampled growth efforts to solve the recurrence (S47a). If the resulting $\bar{x}_{\mathrm{b},40\mathrm{y}} \neq \bar{x}_{\mathrm{B},40\mathrm{y}}$, I plotted these values in log-log scale. I repeated this procedure $10^6$ times.

#### S6 Brain and body size data

The coloured regions in Fig. 2a describe the ranges of variation of brain and body sizes for fossil and extant primate species. Data for hominins are listed in Table S4, where as in ref.[3], since the model considers females only, I used only female data when available. Data for non-hominin primates are from ref.[15], which comprise 56 species of lemurs and lorises, 94 species of lesser apes and cercopithecines, and 7 species of great apes. These non-hominin data are listed in the file PrimateData.jl in the Computer Code. Three outlier, fossil species of cercopithecines listed in ref.[15] are not used here to be able to see the allometric trend with the shaded regions, namely *Aegyptopithecus zeuxis*, *Catopithecus browni*, and *Parapithecus grangeri*.

| Species | brain size in kg | body size in kg |
|---|---|---|
| *H. sapiens* | 1.31 (females) [16;17] | 51.1 (females) [16;17] |
| *H. neanderthalensis* | 1.442 (mixed) [18;19] | 66.4 (females) [18;19] |
| *H. heidelbergensis* | 1.16 (mixed) [20] | 54.24 (mixed) [20] |
| *H. erectus* | 0.98 (mixed) [21] | 55 (females) [21] |
| *H. ergaster* | 0.849 (mixed) [22] | 56 (females) [22] |
| *A. sediba* | 0.42 (mixed) [15] | 26.7 (mixed) [15] |
| *H. habilis* | 0.601 (mixed) [22] | 32 (females) [22] |
| *P. boisei* | 0.49 (mixed) [15] | 35.3 (mixed) [15] |
| *P. robustus* | 0.53 (mixed) [15] | 24.0 (females) [23] |
| *A. africanus* | 0.43 (mixed) [15] | 25.08 (females) [23] |
| *H. floresiensis* | 0.4 (mixed) [24;25] | 25 (mixed) [24;25] |
| *H. naledi* | 0.5 (mixed) [26] | 37.4 (mixed) [26] |
| *A. afarensis* | 0.434 (mixed) [22] | 29 (mixed) [22] |

Table S4: Data used for adult brain and body sizes of hominins. Sexes are indicated in parentheses.

### S7 Evolvability

In Fig. 4e, I plot the angle between the vector of evolutionary change of the geno-phenotype ($\mathrm{d}\bar{\mathbf{z}}/\mathrm{d}\tau = \iota \mathbf{L}_{\mathbf{z}} \partial w/\partial \mathbf{z}$) and the vector of direct selection on the geno-phenotype ($\partial w/\partial \mathbf{z}$), which is given by

$$\cos^{-1}\left( \frac{\frac{\mathrm{d}\bar{\mathbf{z}}^\intercal}{\mathrm{d}\tau} \frac{\partial w}{\partial \mathbf{z}}}{\left\| \frac{\mathrm{d}\bar{\mathbf{z}}}{\mathrm{d}\tau} \right\| \left\| \frac{\partial w}{\partial \mathbf{z}} \right\|} \right). \tag{S49}$$

Also, in Fig. 4f, I plot evolvability sensu Hansen and Houle[27] (their Eq. 1) defined as the scalar projection of the vector of selection response on the vector of direct selection normalized by the strength of selection, which for the evo-devo brain model is given by

$$\frac{\iota \frac{\partial w}{\partial \mathbf{z}^\intercal} \mathbf{L}_{\mathbf{z}}^\intercal \frac{\partial w}{\partial \mathbf{z}}}{\frac{\partial w}{\partial \mathbf{z}^\intercal} \frac{\partial w}{\partial \mathbf{z}}}. \tag{S50}$$

(recall that $\mathbf{L}_{\mathbf{z}}$ is asymmetric so $\mathbf{L}_{\mathbf{z}} \neq \mathbf{L}_{\mathbf{z}}^\intercal$, whereas $\mathbf{G}$ in Eq. 1 of Hansen and Houle[27] is symmetric so $\mathbf{G} = \mathbf{G}^\intercal$). I do not standardize vectors in this measure of evolvability as recommended by Hansen and Houle[27] because doing so changes the magnitude and direction of the vectors and my interest is in the scalar projection of the unmodified selection response on the unmodified selection gradient. Consequently, evolvability so defined is in mixed units as noted by Hansen and Houle[27]. With this definition, if evolvability equals 1 (with mixed units), then the magnitude of the selection response in the direction of direct selection is the same as the magnitude of direct selection; if evolvability equals 0, then either selection response or direct selection is zero or they are orthogonal; if evolvability is greater than 1, then selection response has a magnitude in the direction of direct selection that is greater than the magnitude of direct selection; finally, if evolvability is negative, then selection response has a component opposite to the direction of direct selection.

### S8 Total fitness effects of maintenance metabolic costs

The maintenance metabolic costs $B_i$ are not direct fitness costs in the brain model as fitness does not directly depend on them (Eq. S23). However, they may be total fitness costs. A total fitness cost is defined as a negative total fitness effect on fitness. Total fitness effects of parameter values can be computed with the formulas of the evo-devo dynamics framework for the total effects of environmental variables on fitness.

Consider the column vector $\mathbf{B}_a = (B_\mathrm{b}; B_\mathrm{r}; B_\mathrm{s}; B_\mathrm{k})$ of maintenance metabolic costs experienced at age $a$. This is to consider the effect on fitness of perturbing maintenance metabolic costs at a given age. Now consider the block column vector $\mathbf{B} = (\mathbf{B}_1; \ldots; \mathbf{B}_{N_\mathrm{a}})$ of maintenance metabolic costs experienced at all ages. The total effects of maintenance metabolic costs on fitness are given by

$$\frac{\mathrm{d}w}{\mathrm{d}\mathbf{B}} = \frac{\mathrm{d}\mathbf{x}^\intercal}{\mathrm{d}\mathbf{B}} \frac{\partial w}{\partial \mathbf{x}} \tag{S51}$$

(Layer 4, Eq. S22 of ref. [1]), where the total effects of the maintenance metabolic costs on the phenotype are given by

$$\frac{\mathrm{d}\mathbf{x}^{\intercal}}{\mathrm{d}\mathbf{B}} = \frac{\partial \mathbf{x}^{\intercal}}{\partial \mathbf{B}} \frac{\mathrm{d}\mathbf{x}^{\intercal}}{\mathrm{d}\mathbf{x}} \tag{S52}$$

(Layer 4, Eq. S3 of ref. [1]). As we already have the matrix of developmental feedback (Eq. S36), we only need to compute the matrix of the direct plasticity of the phenotype to change in maintenance costs, which is

$$\left.\frac{\partial \mathbf{x}^{\intercal}}{\partial \mathbf{B}}\right|_{\mathbf{y}=\bar{\mathbf{y}}} = \left.\begin{pmatrix} \frac{\partial \mathbf{x}_1^{\intercal}}{\partial \mathbf{B}_1} & \cdots & \frac{\partial \mathbf{x}_{N_\mathrm{a}}^{\intercal}}{\partial \mathbf{B}_1} \\ \vdots & \ddots & \vdots \\ \frac{\partial \mathbf{x}_1^{\intercal}}{\partial \mathbf{B}_{N_\mathrm{a}}} & \cdots & \frac{\partial \mathbf{x}_{N_\mathrm{a}}^{\intercal}}{\partial \mathbf{B}_{N_\mathrm{a}}} \end{pmatrix}\right|_{\mathbf{y}=\bar{\mathbf{y}}} = \left.\begin{pmatrix} \mathbf{0} & \frac{\partial \mathbf{x}_2^{\intercal}}{\partial \mathbf{B}_1} & \cdots & \mathbf{0} & \mathbf{0} \\ \mathbf{0} & \mathbf{0} & \cdots & \mathbf{0} & \mathbf{0} \\ \vdots & \vdots & \ddots & \vdots & \vdots \\ \mathbf{0} & \mathbf{0} & \cdots & \mathbf{0} & \frac{\partial \mathbf{x}_{N_\mathrm{a}}^{\intercal}}{\partial \mathbf{B}_{N_\mathrm{a}-1}} \\ \mathbf{0} & \mathbf{0} & \cdots & \mathbf{0} & \mathbf{0} \end{pmatrix}\right|_{\mathbf{y}=\bar{\mathbf{y}}} \in \mathbb{R}^{4N_\mathrm{a}\times 4N_\mathrm{a}} \tag{S53}$$

(Layer 2, Eq. S2c of ref. [1]), where direct calculation shows that

$$\frac{\partial \mathbf{x}_{a+1}^{\intercal}}{\partial \mathbf{B}_a} = \begin{pmatrix} -\frac{q_{\mathrm{b}a}}{E_\mathrm{b}} x_{\mathrm{b}a} & -\frac{q_{\mathrm{r}a}}{E_\mathrm{r}} x_{\mathrm{b}a} & -\frac{q_{\mathrm{s}a}}{E_\mathrm{s}} x_{\mathrm{b}a} & \frac{s_\mathrm{k}}{E_\mathrm{k}} x_{\mathrm{b}a}(1-q_{\mathrm{b}a}) \\ -\frac{q_{\mathrm{b}a}}{E_\mathrm{b}} x_{\mathrm{r}a} & -\frac{q_{\mathrm{r}a}}{E_\mathrm{r}} x_{\mathrm{r}a} & -\frac{q_{\mathrm{s}a}}{E_\mathrm{s}} x_{\mathrm{r}a} & -\frac{s_\mathrm{k}}{E_\mathrm{k}} x_{\mathrm{r}a} q_{\mathrm{b}a} \\ -\frac{q_{\mathrm{b}a}}{E_\mathrm{b}} x_{\mathrm{s}a} & -\frac{q_{\mathrm{r}a}}{E_\mathrm{r}} x_{\mathrm{s}a} & -\frac{q_{\mathrm{s}a}}{E_\mathrm{s}} x_{\mathrm{s}a} & -\frac{s_\mathrm{k}}{E_\mathrm{k}} x_{\mathrm{s}a} q_{\mathrm{b}a} \\ 0 & 0 & 0 & -\frac{1}{E_\mathrm{k}} x_{\mathrm{k}a} \end{pmatrix} \in \mathbb{R}^{4\times 4}. \tag{S54}$$

The negative entries in this matrix give the direct “phenotypic costs” of the metabolic costs of maintenance. In turn, the non-negative entry gives the direct “skill benefit” of the metabolic cost of brain maintenance. The zero entries arise because the metabolic cost of memory $B_\mathrm{k}$ does not directly affect tissue mass.

Using these equations, Fig. S12 plots the total fitness effects of maintenance metabolic costs $\mathrm{d}w/\mathrm{d}\mathbf{B}$.

### S9 Supplementary figures

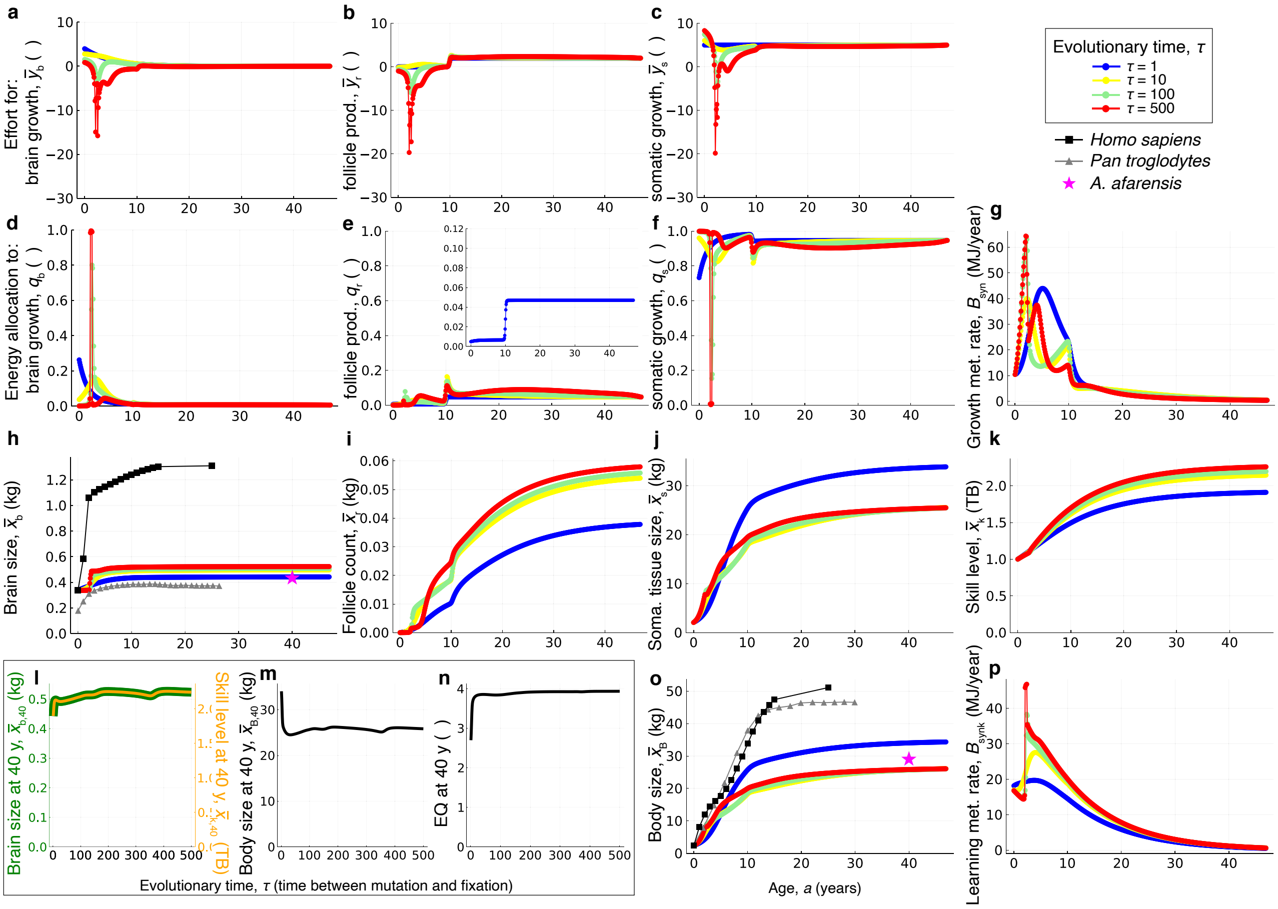

Figure S1: Detailed evo-devo dynamics of brain size for the *afarensis* trajectory in Fig. 4b of the main text. Thus, it is the *afarensis* scenario with the somewhatNaive2 ancestral genotypic traits. Evo-devo dynamics of: **a-c,** genotypic traits, specifically, brain growth, follicle production, and somatic growth; **d-f,** energy allocation to brain growth, follicle production, and somatic growth (**e** inset is a zoom to show the ancestral allocation to follicle production); **g,** the growth metabolic rate (i.e., energy budget for growth, which is allocated according to **d-f**); **h-k,** phenotypic traits, specifically, brain size, follicle count, somatic tissue size, and skill level; **o,** body size; and **p,** the learning metabolic rate. **l-n,** Evolutionary dynamics of (**l**) brain size (green), (**l**) skill level (orange), (**m**) body size, and (**n**) encephalisation quotient (EQ) at 40 years of age.

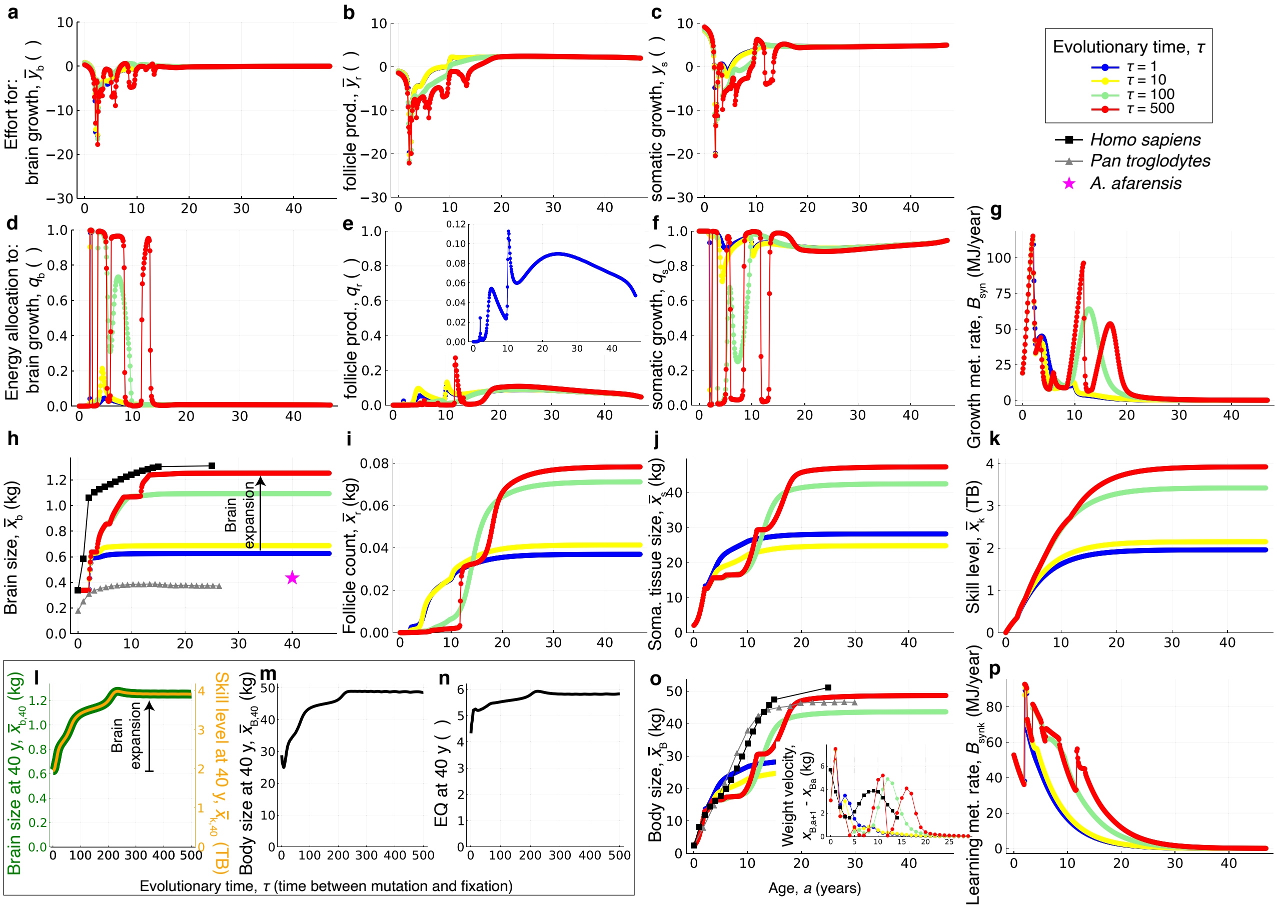

Figure S2: Detailed evo-devo dynamics of brain size for the *sapiens* trajectory in Fig. 2b of the main text. Thus, it is the *sapiens* scenario with the afarensisFromNaive2 ancestral genotypic traits. Evo-devo dynamics of: **a-c,** genotypic traits, specifically, brain growth, follicle production, and somatic growth; **d-f,** energy allocation to brain growth, follicle production, and somatic growth (**e** inset is a zoom to show the ancestral allocation to follicle production); **g,** the growth metabolic rate (i.e., energy budget for growth, which is allocated according to **d-f**); **h-k,** phenotypic traits, specifically, brain size, follicle count, somatic tissue size, and skill level; **o,** body size (**o** inset shows the evolution of weight velocities contrasted to that observed (black squares) in human females); and **p,** the learning metabolic rate. **l-n,** Evolutionary dynamics of (**l**) brain size (green), (**l**) skill level (orange), (**m**) body size, and (**n**) encephalisation quotient (EQ) at 40 years of age.

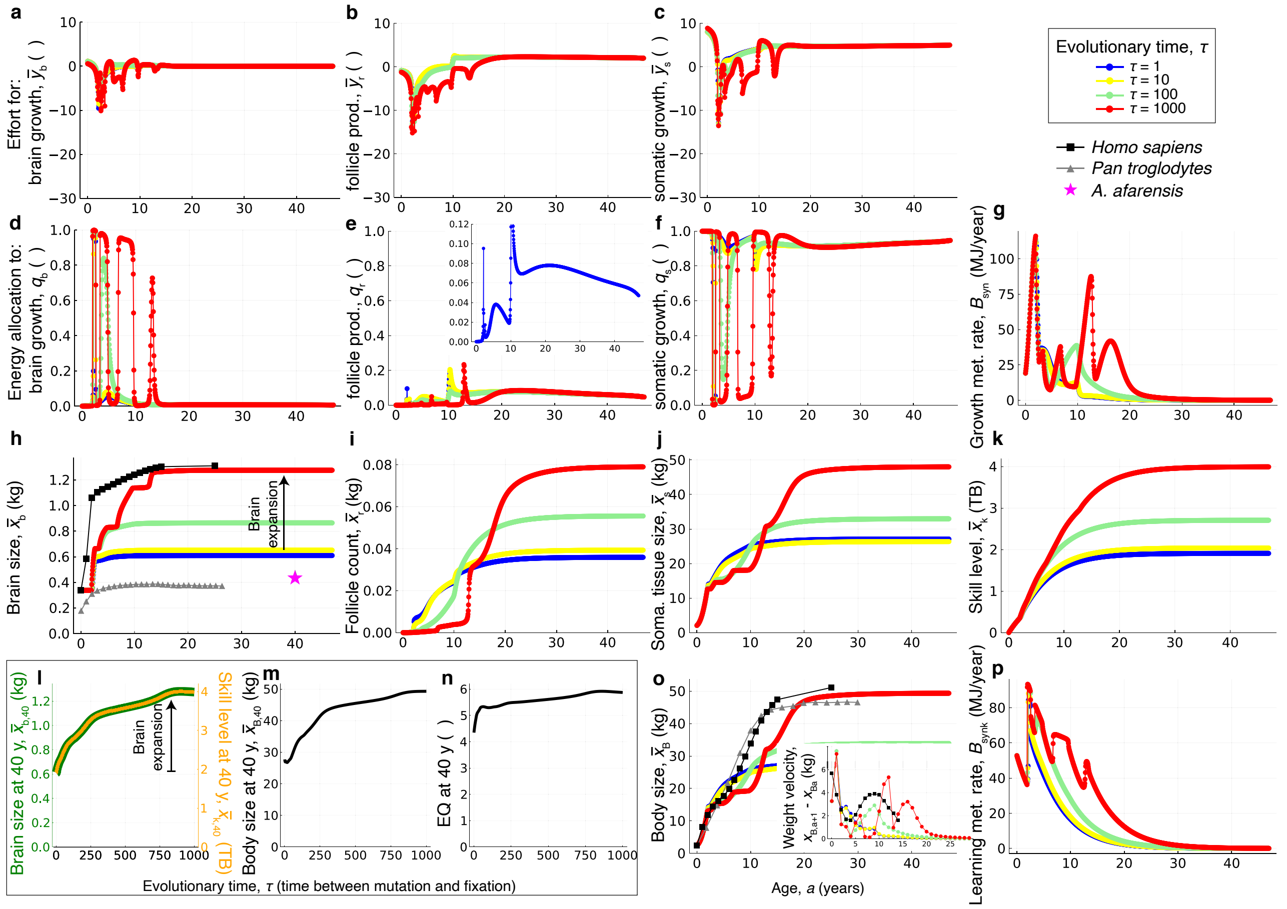

Figure S3: Halving the age bin size does not affect the evolved adult brain and body sizes. The scenario is the same as in Fig. 3 in the main text, except that the age bin is of 1/20 of a year and the final evolutionary time is 1000. The evolutionary dynamics are more than twice as slow as in Fig. 3 (shown in detail in Fig. S2).

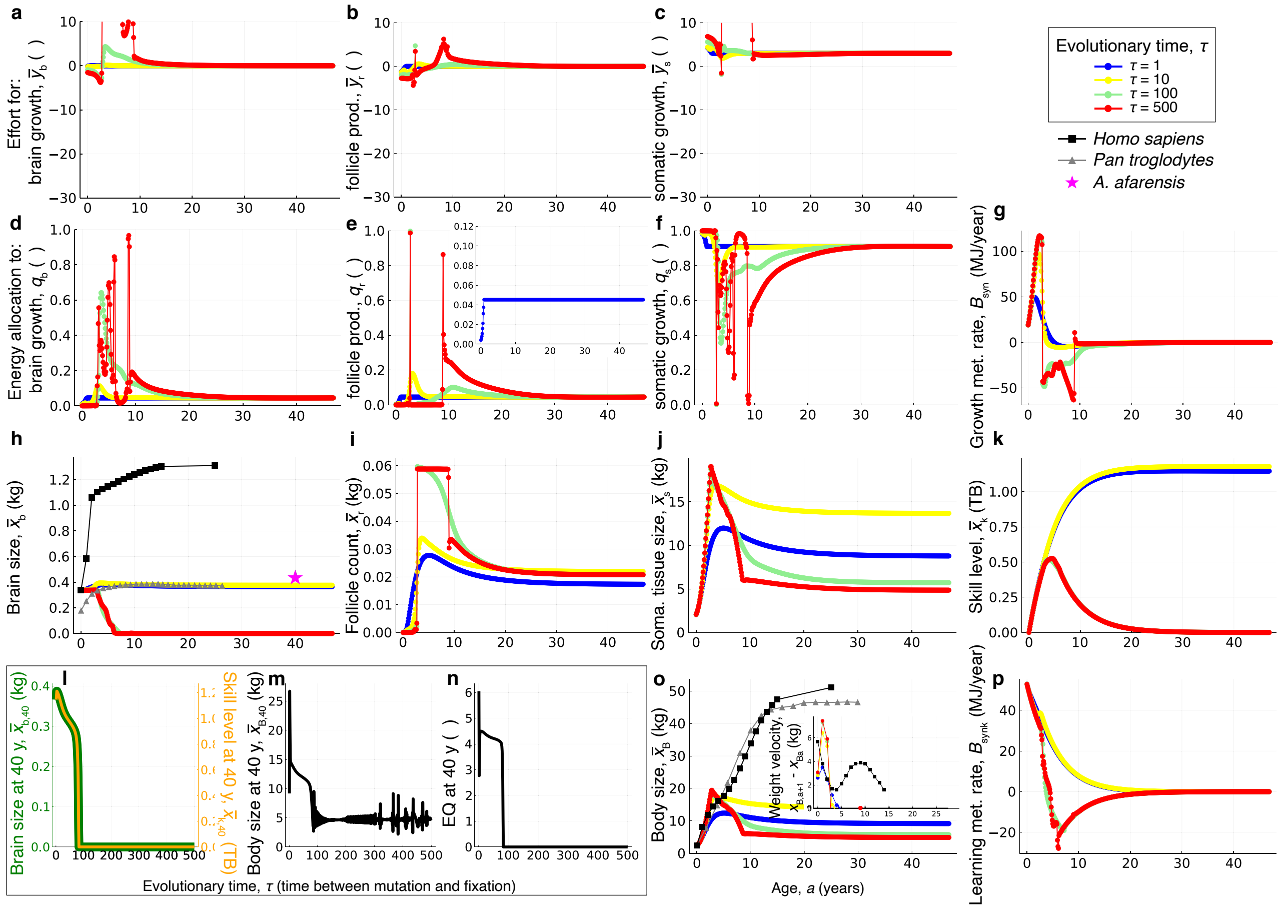

Figure S4: Evo-devo dynamics of brain size with the somewhatNaive ancestral conditions. The scenario is the *sapiens* scenario as in Fig. 3 in the main text, except that the ancestral genotypic traits are the somewhatNaive (blue dots in **a-c**). **l,** Brain size collapses over evolution. These same somewhatNaive conditions cause brain expansion under the ecological scenario as shown in Fig. S5.

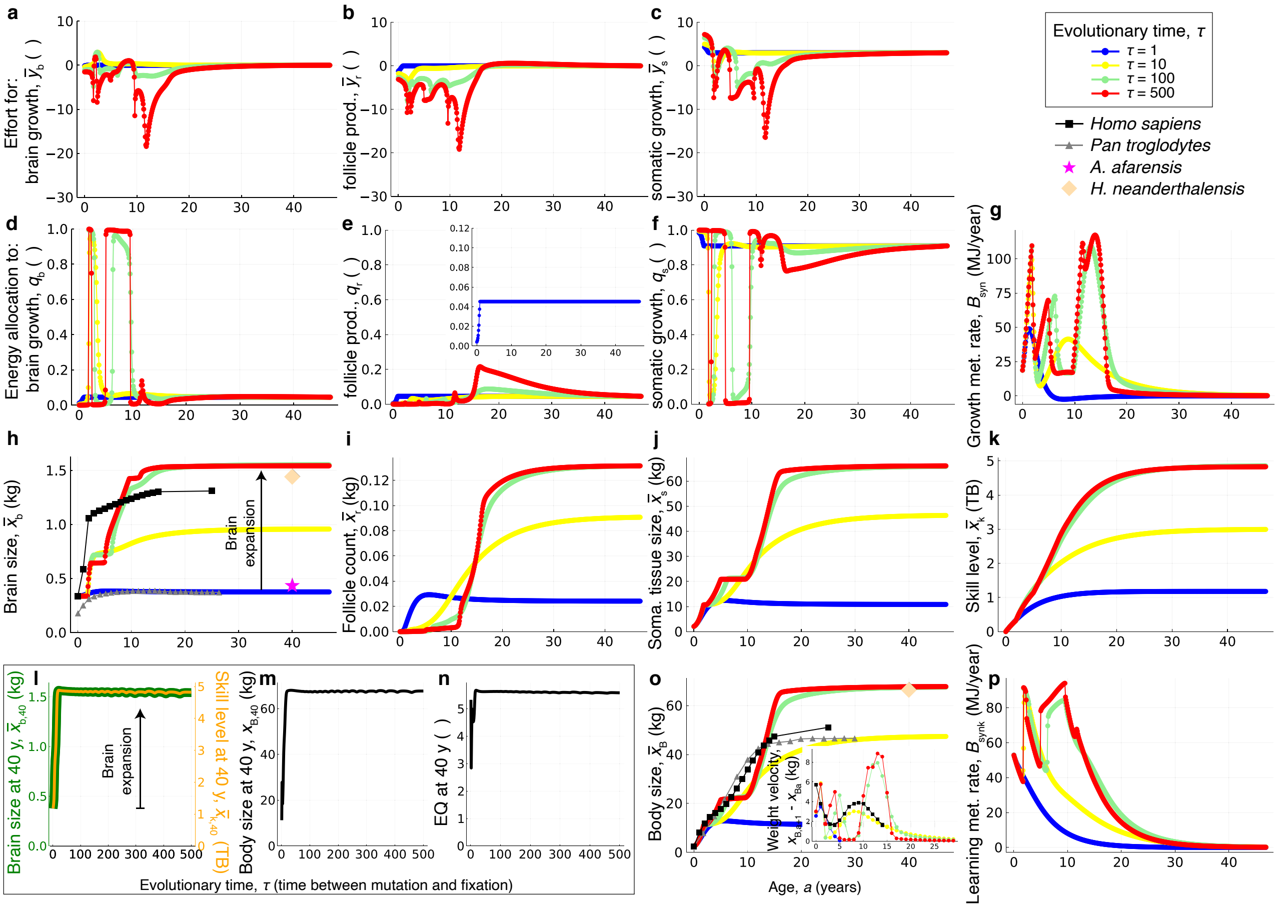

Figure S5: Evo-devo dynamics of brain size with the somewhatNaive ancestral conditions under the ecological scenario. **l,** Brain size more than triples over evolution. The scenario is the same as in Fig. 3 in the main text, except that there are only ecological challenges ($P_1 = 1$) and the ancestral genotypic traits are the somewhatNaive (blue dots in **a**-**c**). These same ancestral genotypic traits cause brain collapse under the *sapiens* scenario as shown in Fig. S4. The red dots in **d**-**k** and **o**,**p** broadly but not exactly correspond to the solutions found by ref.[2] (their Fig. 3E-H) who studied the ecological scenario at evolutionary equilibrium using optimal control.

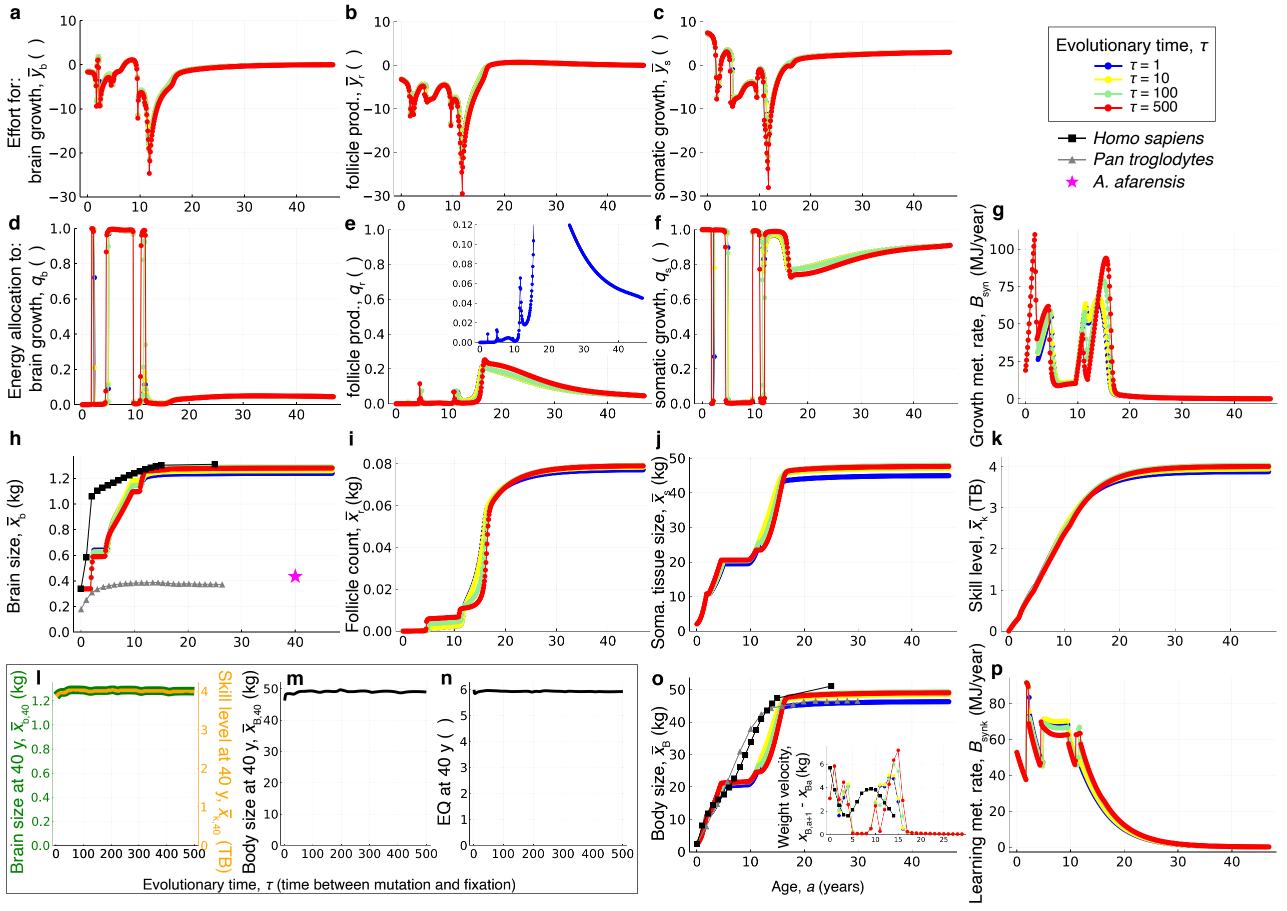

Figure S6: Evo-devo dynamics of brain size under the *sapiens* scenario with ecologically quasi-optimal ancestral genotype. The scenario is the same as in Fig. 3 in the main text, except that the ancestral genotypic traits are the evolved ones under the ecological scenario (blue dots in **a-c** are the red dots in Fig. S5a-c). **l,** Brain size is already large at the start of the evolutionary process.

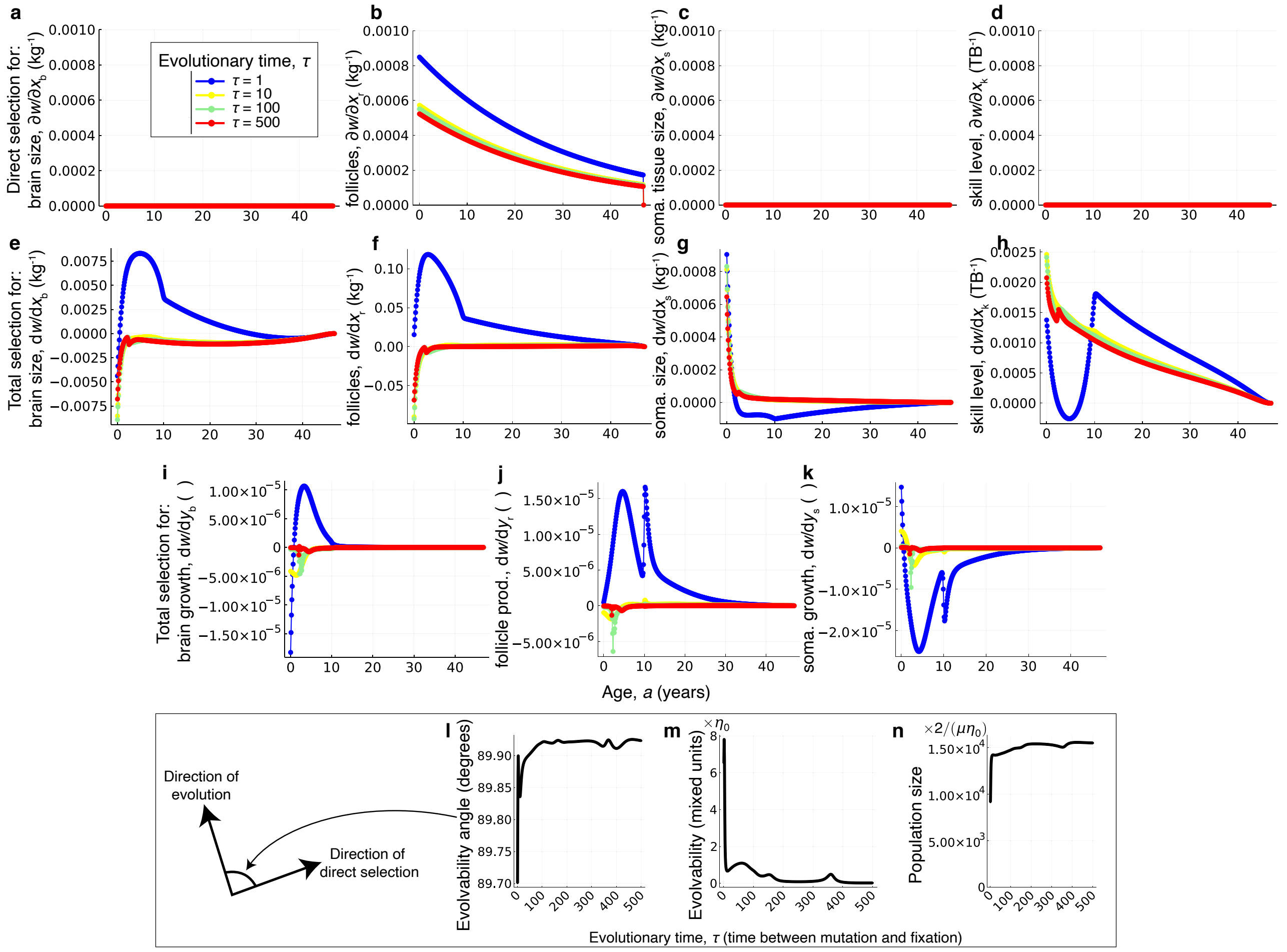

Figure S7: The action of selection for the *afarensis* trajectory in Fig. 2b. **a-d,** Direct selection on brain size, follicle count, somatic tissue size, and skill level at each age over evolutionary time. **e-h,** Total selection on brain size, follicle count, somatic tissue size, and skill level at each age over evolutionary time. **i-k,** Total selection on effort for brain growth, follicle production, and somatic growth at each age over evolutionary time. **l,** Angle between the direction of evolution and direct selection, both of the geno-phenotype, over evolutionary time. **m,** Evolvability over evolutionary time: evolvability equal to 0 here means no evolution despite selection; SI section S7; Eq. 1 of ref.[27]). **n,** Population size (plot of $\frac{1}{2}\mu\bar{n}^*\eta_0$, so the indicated multiplication yields population size). All plots are for the evolutionary process of Fig. S1.

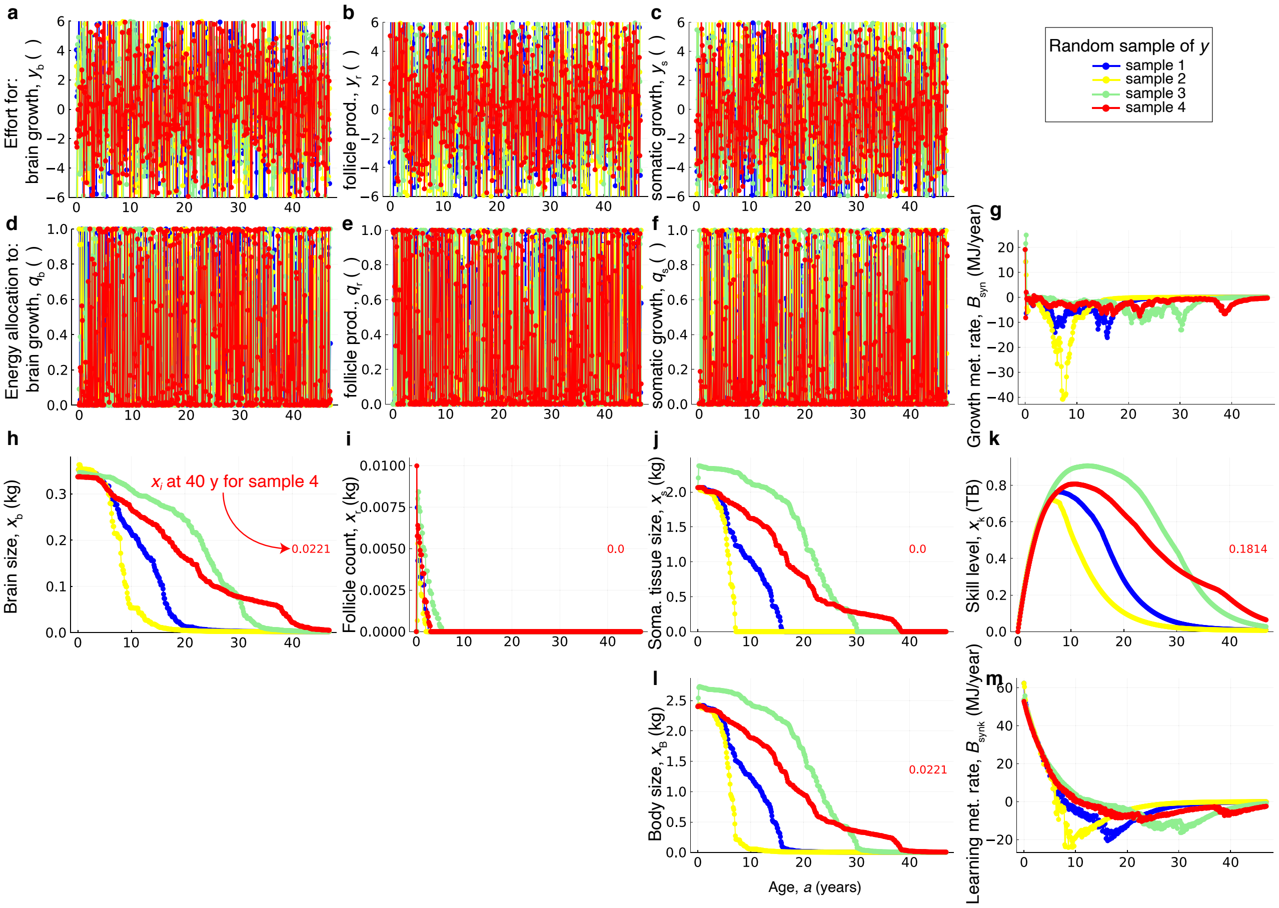

Figure S8: Failed organisms with random growth efforts. **a-c,** Four samples of growth efforts drawn from a normal distribution with mean 0 and standard deviation 4. **h-k,** With these growth efforts, tissues and skill collapse over development. **l,** Body size also collapses over development. Numbers in red in panels **h-l** give the value of $x_{i40}$ at 40 years of age for sample 4. In the four samples shown, all body is composed of brain at age 40 (e.g., red numbers in **h** and **l** are equal). This is because the growth metabolic rate $B_{\text{syn}}$ is negative (panel **g**), so tissues shrink and from the developmental map of tissues (Eq. S8), the most expensive tissue to grow (i.e., that with the highest $E_i$) shrinks the slowest. As the brain is the second most expensive tissue to grow and follicles are more expensive but of smaller total mass, brain size takes longer to reach a zero value. Consequently, the body of these organisms is entirely composed of brain at 40 y of age.

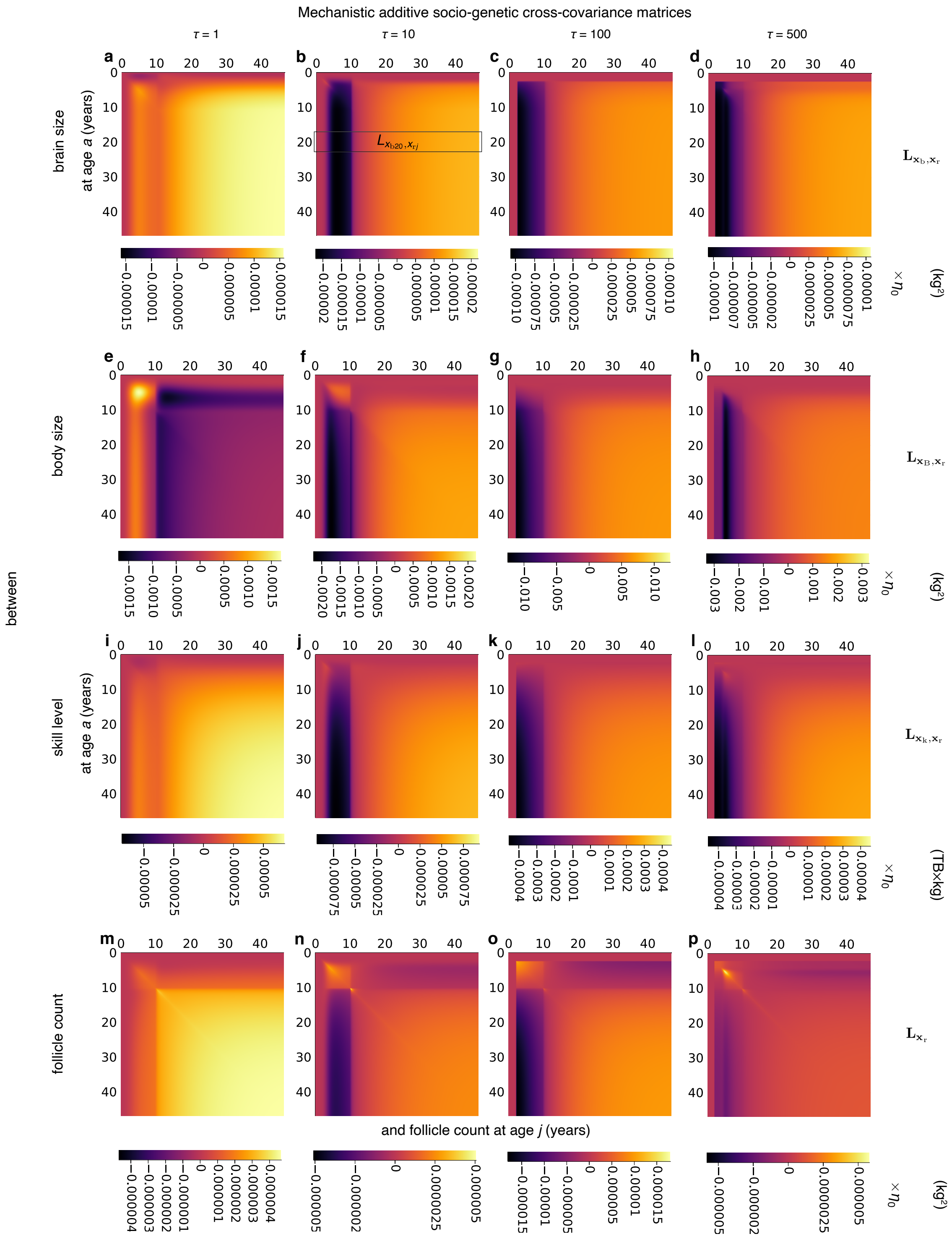

Figure S9: The action of constraint in the *afarensis* trajectory in Fig. 2b. Mechanistic socio-genetic cross-covariance matrix between **a-d,** brain size and follicle count, **a-d,** body size and follicle count, **e-h,** skill level and follicle count, and **i-l,** follicle count and itself. All plots are for the evolutionary process of Fig. S1. Bar legends have different limits so that patterns are visible (bar legend limits are $\{-l, l\}$, where $l = \max(|L_{x_{ba}, x_{rj}}|)$ over $a$ and $j$ for each $\tau$).

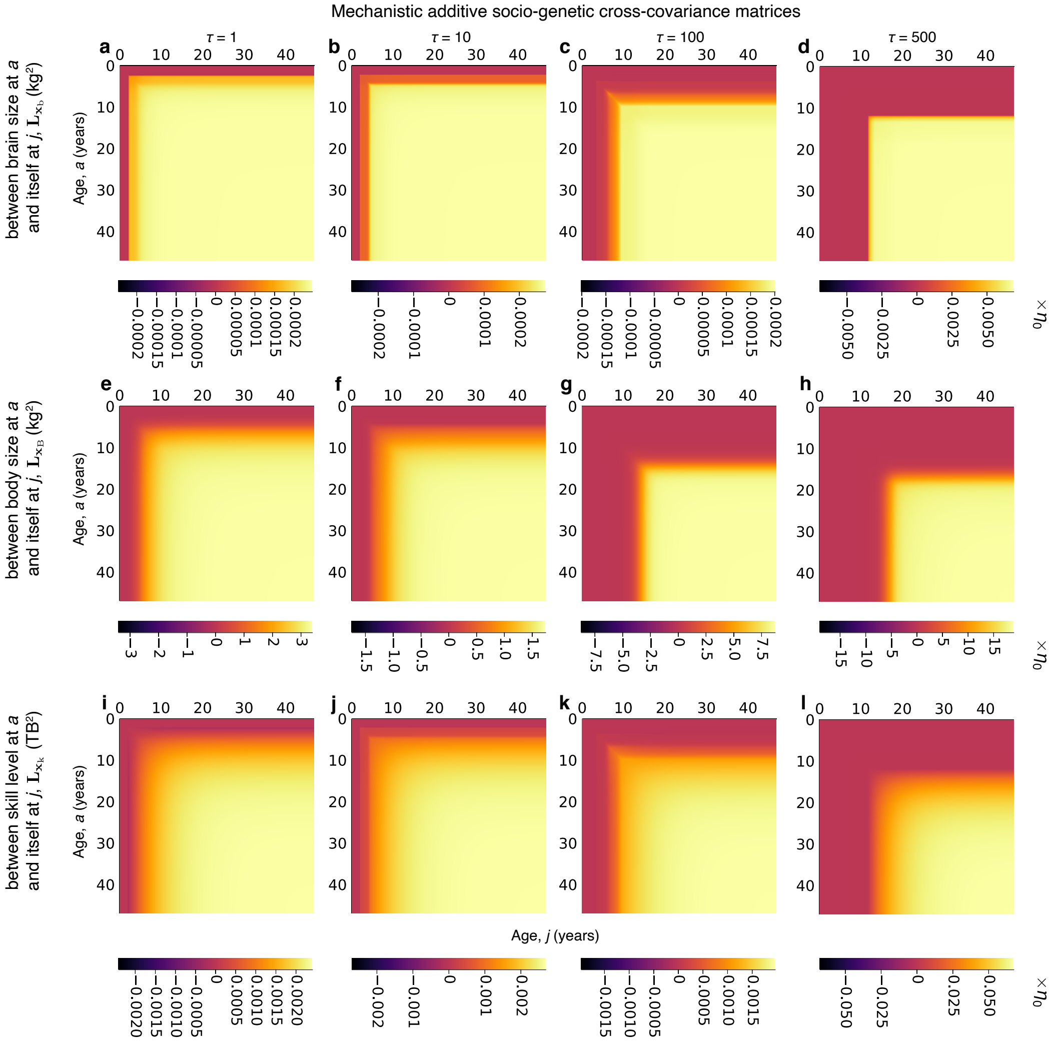

Figure S10: Further mechanistic socio-genetic covariance matrices between a phenotype and itself for the *sapiens* trajectory in Fig. 2b. **a-d,** Mechanistic socio-genetic cross-covariance matrix between brain size (at the ages on vertical axes) and itself (at the ages on horizontal axes) over evolutionary time. **e-h,** Mechanistic socio-genetic cross-covariance matrix between body size (at the ages on vertical axes) and itself (at the ages on horizontal axes) over evolutionary time. **i-l,** Mechanistic socio-genetic cross-covariance matrix between skill level (at the ages on vertical axes) and itself (at the ages on horizontal axes) over evolutionary time. All plots are for the evolutionary process of Fig. 3 in the main text.

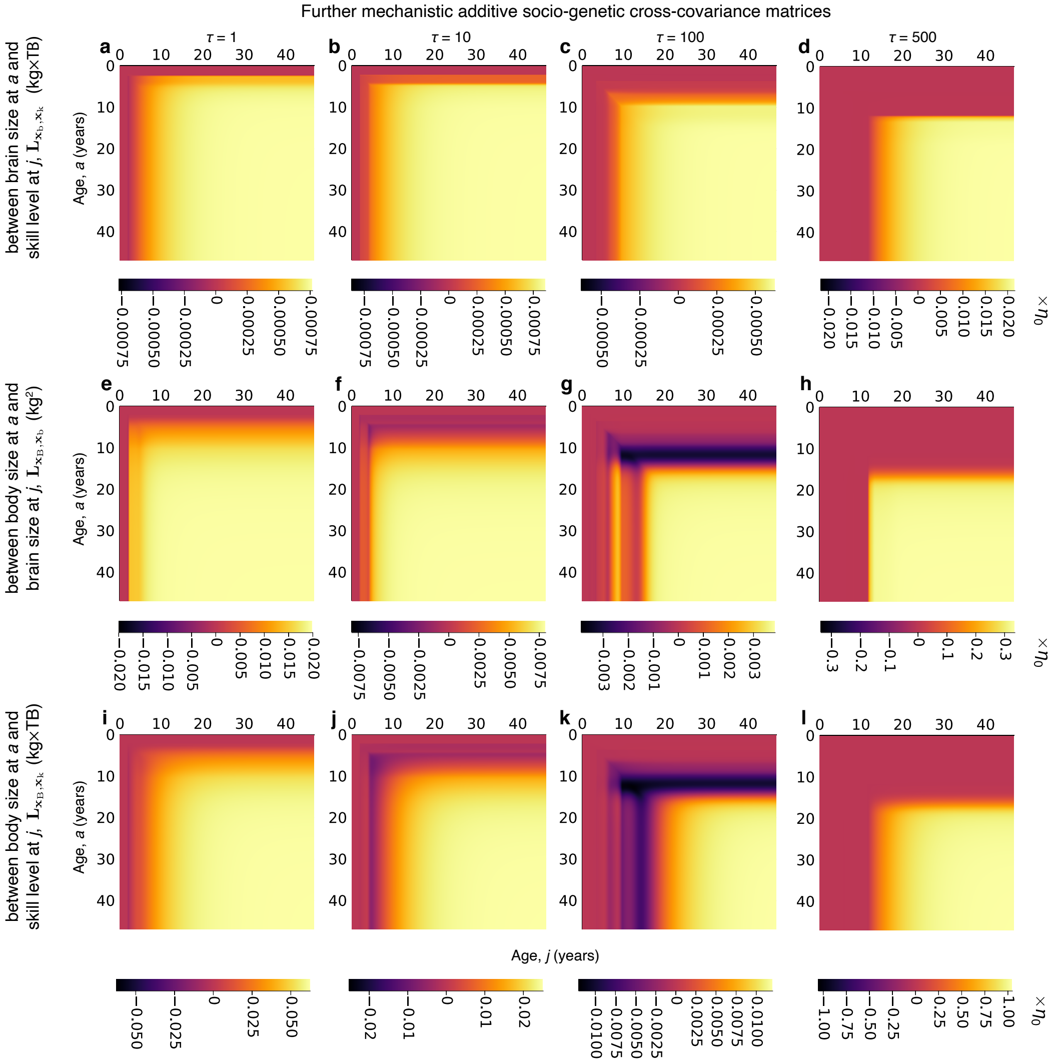

Figure S11: Further mechanistic socio-genetic covariance matrices between different phenotypes for the *sapiens* trajectory in Fig. 2b. **a-d,** Mechanistic socio-genetic cross-covariance matrix between brain size (at the ages on vertical axes) and skill level (at the ages on horizontal axes) over evolutionary time. **e-h,** Mechanistic socio-genetic cross-covariance matrix between body size (at the ages on vertical axes) and brain size (at the ages on horizontal axes) over evolutionary time. **i-l,** Mechanistic socio-genetic cross-covariance matrix between body size (at the ages on vertical axes) and skill level (at the ages on horizontal axes) over evolutionary time. All plots are for the evolutionary process of Fig. 3 in the main text.

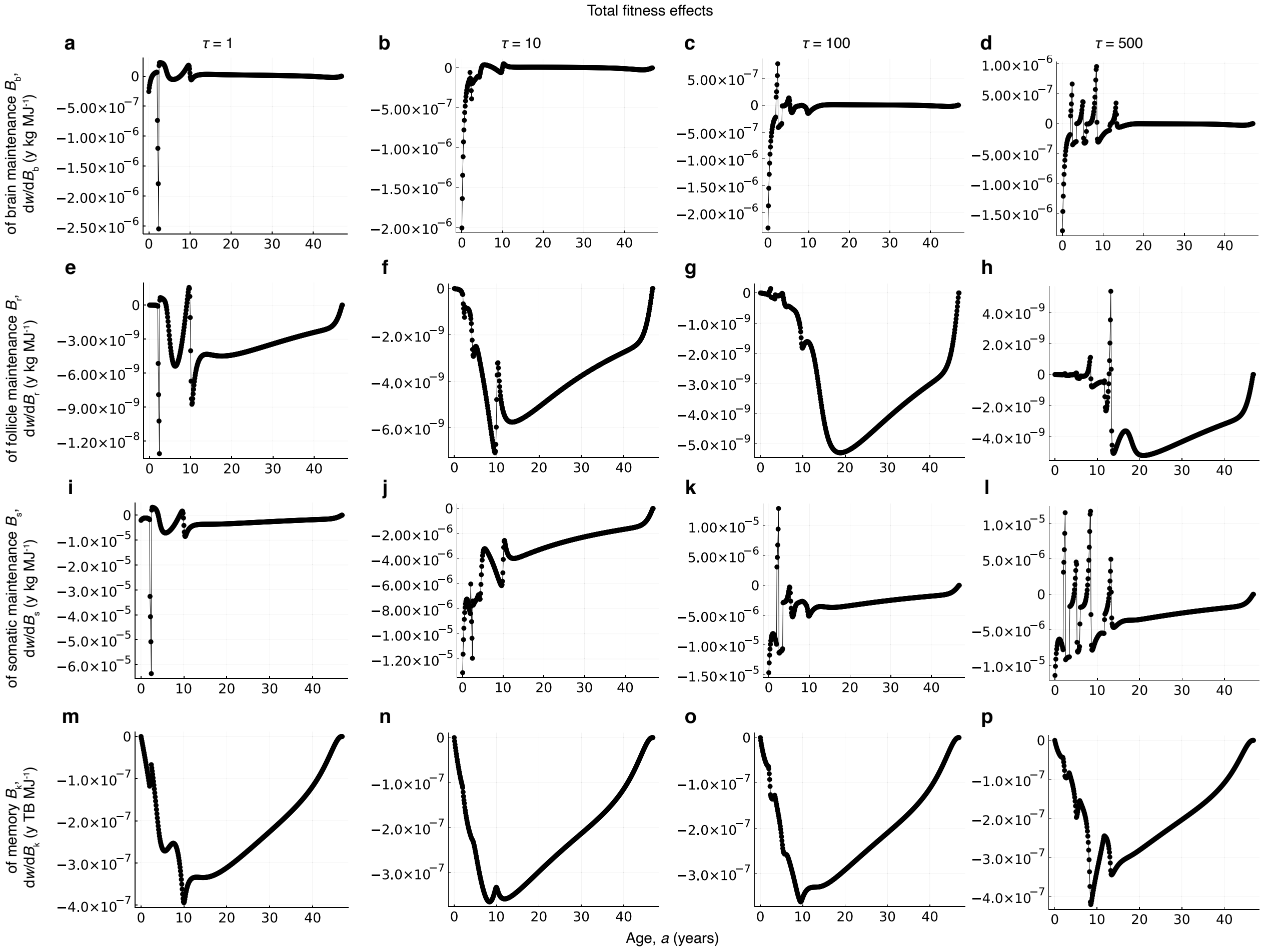

Figure S12: Total fitness effects of maintenance metabolic costs for the *sapiens* trajectory in Fig. 2b. Each panel shows the total effect on fitness of perturbing the indicated maintenance metabolic cost at the indicated age. When the effect is negative, the corresponding metabolic cost is a total fitness cost at that age. When the effect is positive, the corresponding metabolic cost is a total fitness benefit at that age. The metabolic costs of brain maintenance (**a**-**d**) and follicle maintenance (**e**-**h**) are occasionally a total fitness benefit. Among tissues, the metabolic cost of somatic maintenance (**i**-**l**) has some of the most substantial total fitness effects (compare vertical axes in **a**-**l**). The metabolic cost of memory (**m**-**p**) is a total fitness cost over development and evolution.

#### References

1. González-Forero, M. A mathematical framework for evo-devo dynamics. *Theor. Popul. Biol.* **155**, 24–50 (2024).
2. González-Forero, M., Faulwasser, T. & Lehmann, L. A model for brain life history evolution. *PLOS Comp. Biol.* **13**, e1005380 (2017).
3. González-Forero, M. & Gardner, A. Inference of ecological and social drivers of human brain-size evolution. *Nature* **557**, 554–557 (2018).
4. Caswell, H. *Sensitivity Analysis: Matrix Methods in Demography and Ecology* (Springer Open, Cham, Switzerland, 2019).
5. West, G. B., Brown, J. H. & Enquist, B. J. A general model for ontogenetic growth. *Nature* **413**, 628–631 (2001).
6. Charlesworth, B. *Evolution in age-structured populations* (Cambridge Univ. Press, 1994), 2nd edn.
7. Dieckmann, U. & Law, R. The dynamical theory of coevolution: a derivation from stochastic ecological processes. *J. Math. Biol.* **34**, 579–612 (1996).
8. Hamilton, W. D. The moulding of senescence by natural selection. *J. Theor. Biol.* **12**, 12–45 (1966).
9. Caswell, H. A general formula for the sensitivity of population growth rate to changes in life history parameters. *Theor. Popul. Biol.* **14**, 215–230 (1978).
10. Baudisch, A. Hamilton's indicators of the force of selection. *Proc. Natl. Acad. Sci. USA* **102**, 8263–8268 (2005).
11. Morrissey, M. B. Selection and evolution of causally covarying traits. *Evolution* **68**, 1748–1761 (2014).
12. Wagner, G. P. On the eigenvalue distribution of genetic and phenotypic dispersion matrices: Evidence for a non-random organization of quantitative character variation. *J. Math. Biol.* **21**, 77–95 (1984).
13. Fisher, R. A. XV.—The correlation between relatives on the supposition of Mendelian inheritance. *Trans. Roy. Soc. Edinb.* **52**, 399–433 (1918).
14. Greene, V. L. An algorithm for total and indirect causal effects. *Political Methodology* **4**, 369–381 (1977).
15. Smaers, J. B. *et al.* The evolution of mammalian brain size. *Sci. Adv.* **7**, eabe2101 (2021).
16. Kuzawa, C. W. *et al.* Metabolic costs and evolutionary implications of human brain development. *Proc. Nat. Acad. Sci. USA* **111**, 13010–13015 (2014).
17. Dekaban, A. S. & Sadowsky, D. Changes in brain weights during the span of human life: Relation of brain weights to body heights and body weights. *Ann. Neurol.* **4**, 345–356 (1978).
18. Froehle, A. W. & Churchill, S. E. Energetic competition between Neandertals and anatomically modern humans. *PaleoAnthropology* 96–116 (2009).
19. Ruff, C. B., Trinkaus, E. & Holliday, T. W. Body mass and encephalization in Pleistocene *Homo*. *Nature* **387**, 173–176 (1997).
20. Rightmire, G. P. Brain size and encephalization in early to mid-pleistocene *Homo*. *Am. J. Phys. Anthropol.* **124**, 109–123 (2004).
21. McHenry, H. M. Tempo and mode in human evolution. *Proc. Natl. Acad. Sci. USA* **91**, 6780–6786 (1994).
22. McHenry, H. M. & Coffing, K. *Australopithecus* to *Homo*: transformations in body and mind. *Annu. Rev. Anthropol.* **29**, 125–146 (2000).
23. Grabowski, M., Hatala, K. G., Jungers, W. L. & Richmond, B. G. Body mass estimates of hominin fossils and the evolution of human body size. *J. Hum. Evol.* **85**, 75–93 (2015).
24. Brown, P. *et al.* A new small-bodied hominin from the Late Pleistocene of Flores, Indonesia. *Nature* **431**, 1055–1061 (2004).
25. Kubo, D., Kono, R. T. & Kaifu, Y. Brain size of *Homo floresiensis* and its evolutionary implications. *Proc. R. Soc. B* **280**, 20130338 (2013).
26. Garvin, H. M. *et al.* Body size, brain size, and sexual dimorphism in *Homo naledi* from the Dinaledi Chamber. *J. Hum. Evol.* **111**, 119–138 (2017).
27. Hansen, T. F. & Houle, D. Measuring and comparing evolvability and constraint in multivariate characters. *J. Evol. Biol.* **21**, 1201–1219 (2008).
